# THREAD: a Bayesian nonparametric framework for resolving polygenic risk into multicellular programs

**DOI:** 10.64898/2026.11.25.769208

**Authors:** Jiacheng Lou, Teng Li, Xuyuan Chen, Ruizhi Zhou, Han Wang, Yan Dora Zhang

## Abstract

Genome-wide association studies have linked thousands of loci to complex disease, yet translating these associations into the cellular mechanisms that generate risk remains difficult. Existing methods relate genetic risk to individual cell types or, at single-cell resolution, to individual cells, yet how that risk is organized across coordinated cellular programs remains less explored. We present THREAD, a Bayesian nonparametric framework that integrates GWAS summary statistics with single-cell expression: a Dirichlet process mixture over genes partitions heritable risk into a data-driven number of latent modules, each with its own cell-subtype coefficients and a likelihood whose heteroscedasticity is fixed by local linkage disequilibrium. Modules consistently prioritized coordinated sets of cell subtypes, which we interpret as multicellular programs. In simulations, single-coefficient models misattribute heterogeneous risk rather than merely losing power, whereas THREAD recovers the underlying programs across a broad range of heterogeneous architectures. Across Alzheimer’s disease, schizophrenia and bipolar disorder, colorectal cancer and breast cancer, THREAD resolves disease-specific multicellular programs that reproduce in independent cohorts and align with clinical progression, molecular subtype and survival, shifting genetic mapping from identifying individual risk-associated cell subtypes to resolving the multicellular programs through which polygenic risk is organized.

## Introduction

Genome-wide association studies (GWAS) have linked thousands of genetic loci to complex human traits and diseases, yet because most risk variants lie in non-coding regions, and translating statistical associations into biological mechanisms remains a central challenge. Because regulatory variants act in a cell-type-dependent manner, a natural route is to locate genetic risk in the cellular contexts in which it is realised. Single-cell RNA sequencing (scRNA-seq) now resolves the cellular composition of tissues at fine granularity, and integrating GWAS summary statistics with single-cell expression offers a route to map genetic susceptibility onto the cell populations in which risk variants act, connecting a macroscopic phenotype to its cellular origins.

Over the past decade, substantial progress has been made in linking genetic risk to cellular context through the integration of GWAS and transcriptomic data. Polygenic regression frameworks such as LDSC-SEG^1^, MAGMA gene-property analysis^2^ and RolyPoly^3^ have demonstrated that heritable risk is enriched in specific tissues and cell types, providing important biological insights into the cellular origins of complex disease. More recently, approaches including scPagwas^4^ and scDRS^5^ have extended this effort to single-cell resolution, enabling the identification of disease-associated cellular populations and states. Despite their differences, these methods share a common formulation that the translation of cellular expression into genetic risk is the same for every gene: that a given cell state contributes to heritability in a fixed manner regardless of which gene it is acting through.

This formulation leaves an important aspect of genetic architecture unaddressed. Two genes expressed with identical specificity in the same microglial, neuronal or stromal state can carry profoundly different genetic burden, because risk is not conferred by expression per se but by the function a gene performs within a particular biological process. The contribution of a cell state to heritability is therefore conditional on the set of genes through which it acts, and a single cell state may be a dominant conduit of risk for one group of genes while being inert for another. Conversely, a coordinated biological process typically recruits several cell states at once, so that the genetic risk carried by a functionally coherent group of genes is distributed across a constellation of cellular contexts rather than concentrated in one. Accumulating evidence from neurodegenerative disorders, psychiatric disease and cancer also suggests that disease risk is distributed across multiple interacting cell populations engaged in distinct but coordinated processes ^6,7^. Collapsing this gene-dependent structure into a single ranking of cell types averages over mechanistically distinct pathways, dilutes the signal of cell populations that are relevant to only a subset of the risk architecture, and obscures the multicellular organisation of disease.

Here we present THREAD (Trait Heterogeneity Regression through Expression Annotation and Dirichlet processes), a Bayesian nonparametric framework that decomposes polygenic risk into interpretable multicellular programs. Rather than asking which cell types are relevant to a trait, it asks through which groups of genes, acting through which constellations of cell states, polygenic risk is transmitted. The framework is built on a generative account in which the genetic variance at each gene arises from an unobserved functional context—a latent program—that dictates how the gene’s expression across cell states is converted into heritable risk. Because the precision with which genetic contributions can be estimated varies markedly across genes, partly owing to local linkage disequilibrium (LD), THREAD derives the sampling uncertainty of each gene’s evidence analytically and incorporates it into inference. Finally, since single-cell measurements are sparse and related cell states are often collinear, THREAD reports only those program– cell-state associations that are supported by complementary Bayesian and frequentist criteria. The resulting programs are defined jointly by coherent sets of genes, cell states, pathways and regulatory mechanisms, providing an interpretable unit for studying how genetic risk is organised across complex cellular systems.

We benchmarked THREAD on more than a thousand simulated datasets spanning homogeneous to highly heterogeneous architectures, including regimes dominated by rare programs and by non-informative genes, and found that a single shared cell-type relationship does not merely lose power under heterogeneity but misattributes risk across programs. Applied to Alzheimer’s disease, schizophrenia, bipolar disorder, colorectal cancer and breast cancer, THREAD resolved genetic risk into coherent programs that revealed disease-specific architectures not apparent from global cell-type rankings. In every case the programs were reproducible in independent cohorts and aligned with clinically meaningful axes, from druggable-target distribution and tumour stage to molecular subtype and patient survival. Together these results indicate that, across neurodegeneration, psychiatric disorders and cancer, polygenic risk is organised into biologically coherent, multicellular programs that THREAD can recover from data.

## Results

### Overview of the THREAD framework

THREAD is built on a generative account in which the genetic variance at each gene arises from an unobserved functional context—a latent program—that determines how the gene’s expression across cell states is translated into heritable risk (Fig. 1).

**Figure 1.**
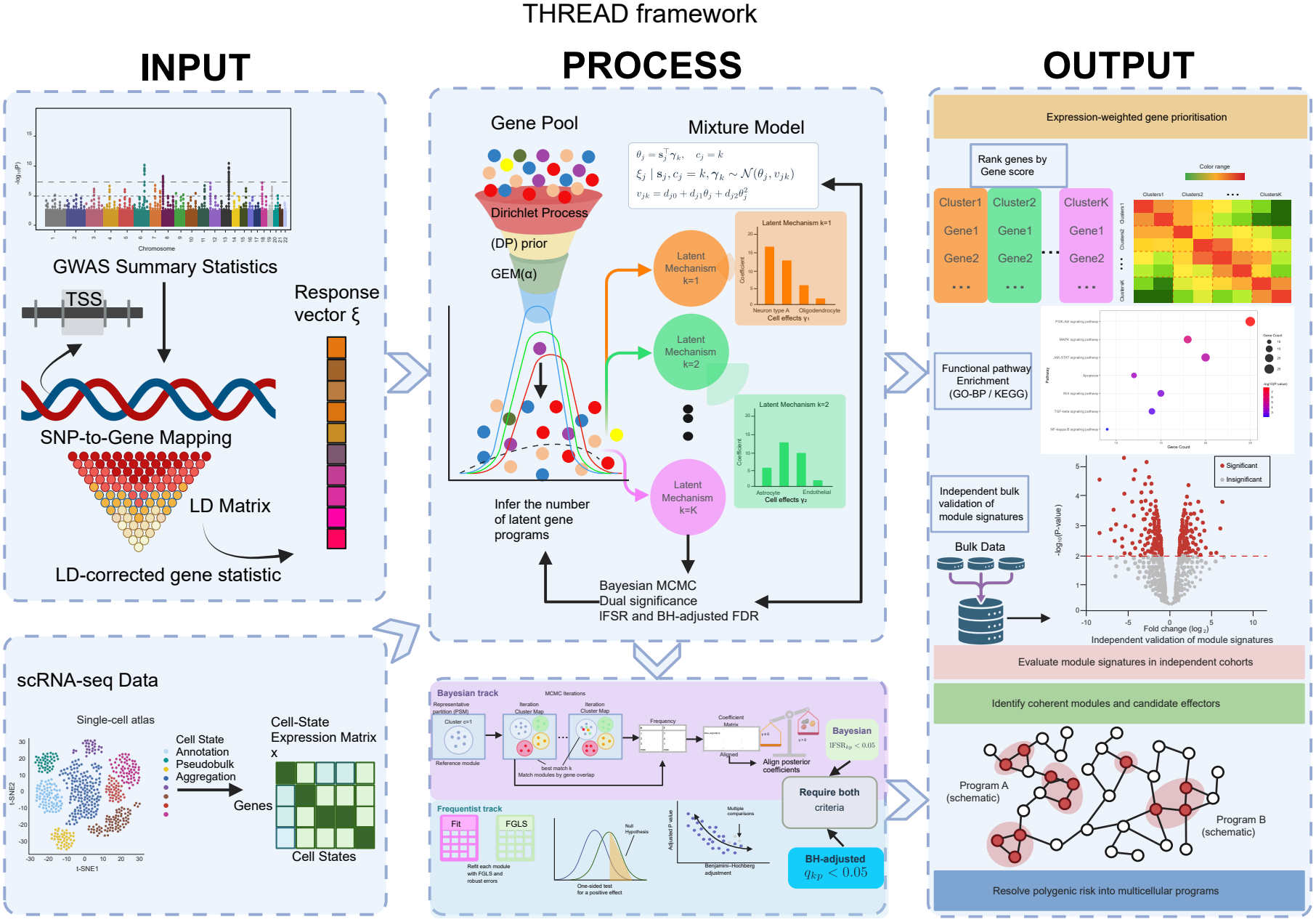
Overview of the THREAD framework. **a, Input**. Variant-level GWAS summary statistics are mapped to cis-regions around transcription start sites (TSS) accounting for linkage disequilibrium (LD) to derive gene-level association statistics (***y***). Single-cell RNA-seq (scRNA-seq) profiles are aggregated into a standardized gene-by-cell-state expression specificity matrix (***X***). **b, Process**. A Dirichlet Process Mixture (DPM) prior with a GEM(*α*) construction partitions genes into latent mechanistic modules (*k* = 1, …, *K*) via MCMC sampling. Candidate associations are filtered through a dual-track criterion requiring both Bayesian credibility (lFSR < 0.05) and frequentist significance (FGLS Benjamini–Hochberg-adjusted *q* < 0.05). **c, Output**. THREAD outputs gene co-clustering matrices, module-specific cell-state profiles, and prioritized driver genes for downstream pathway enrichment, bulk transcriptomic validation, and therapeutic target identification.

For gene *j*, let *θ*_*j*_ denote the per-SNP effect-size variance at its locus and ***s***_*j*_ its expression-specificity profile across *P* cell states. Conditional on membership in program *k*, THREAD models 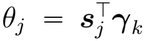, so that the vector ***γ***_*k*_ encodes the cellular basis of risk shared by the genes of that program, and the same cell state may carry a strong effect in one program and none in another. Programs are not specified in advance: the translation rules ***γ***_*k*_ are modelled as draws from a random probability measure with a Dirichlet process prior, whose discreteness causes genes governed by the same rule to coalesce while the number, size and composition of programs emerge from the data. Within each program, a hierarchical Laplace prior on cell-state effects isolates the few cellular contexts that constitute its substrate from the many that do not.

The evidence entering this model is derived from GWAS summary statistics without reference to any external gene set. For each gene, THREAD summarises the marginal effect estimates of SNPs mapped to its locus as an unbiased, LD-corrected estimate *ξ*_*j*_ of *θ*_*j*_. Rather than estimating the precision of *ξ*_*j*_ empirically, we derive its sampling variance under the polygenic model and show that it is a quadratic function of *θ*_*j*_ with coefficients fixed by local LD structure, GWAS sample size and residual variance (Methods and Supplementary Note). The reliability of each gene’s evidence is therefore known rather than assumed, and a precisely estimated locus and an imprecisely estimated one of equal magnitude are weighed differently by construction.

Posterior inference over programs, gene memberships and cell-state effects is performed jointly by Markov chain Monte Carlo (MCMC). Because program labels carry no intrinsic identity, THREAD summarises the posterior through co-clustering probabilities, from which a representative partition of genes is extracted, and quantifies the directional certainty of each program–cell-state effect by its local false sign rate. To guard against prior-induced shrinkage and the collinearity of related cell states, each candidate effect must also survive an unpenalised within-program refit by feasible generalised least squares with false-discovery control; only effects supported on both grounds are reported, with the unshrunk estimate serving as the effect size. We refer to each inferred group of genes as a gene module and to the module, together with its prioritised cell states and the biological processes its genes implicate, as a multicellular program. Building on these programs, THREAD ranks constituent genes by an expression-weighted prioritisation score and links each program to upstream regulators, protein–protein interaction networks, druggable targets and clinical phenotypes in independent cohorts (Fig. 1).

### THREAD recovers cell-subtype programs that pooled models misattribute in simulation

Before applying THREAD to disease, we asked a more basic question: under what genetic architectures does explicitly modeling the cellular origin of heritability change the answer rather than merely refine it? To address it we simulated GWAS data from a specified cell-subtype architecture (Methods). For each of *G* genes we drew an expression-specificity profile across five cell subtypes—with no built-in cell-subtype structure, so that all program structure enters through the coefficients alone—and assigned the gene to one of several latent programs, each defined by a distinct, sparse subset of causal cell subtypes that set the gene’s genetic variance. From these per-gene variances we simulated GWAS summary statistics for 500 SNPs per gene under linkage disequilibrium and summarised them into an LD-corrected per-gene heritability—the quantity THREAD regresses on expression specificity in real data (Methods). Critically, this estimate is not observed cleanly: its sampling variance is, by derivation rather than assumption, a quadratic function of its own mean, with coefficients fixed entirely by the local LD (Methods), so the noise grows with the heritability itself—the heteroscedastic structure carried by real GWAS summary statistics. Once LD and this heteroscedastic noise are imposed, the cleanly separated program structure is heavily obscured (Figure 2a), making recovery genuinely hard rather than a linear toy.

**Figure 2.**
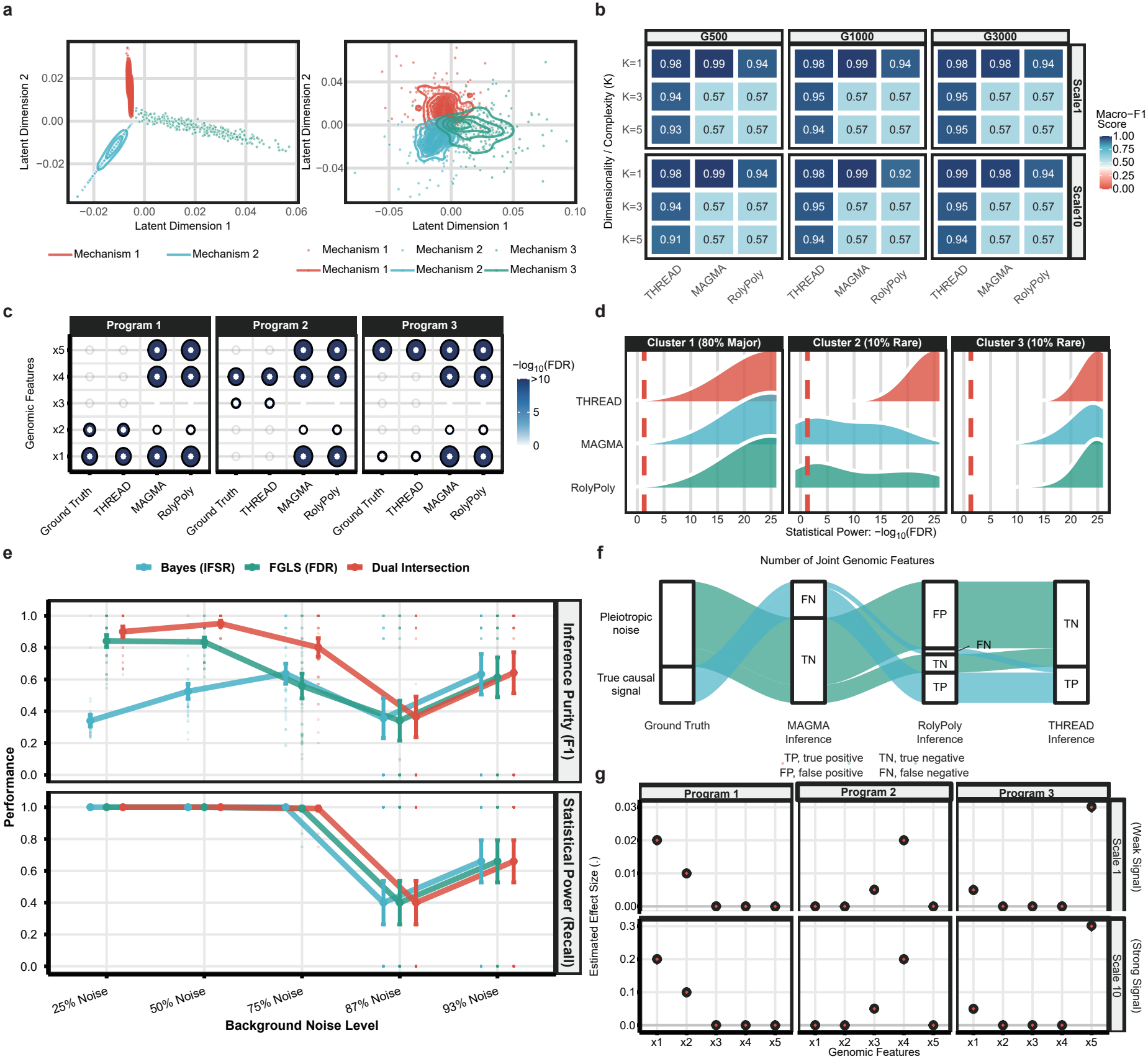
Simulation benchmarks of THREAD. **a**, Latent representations of simulated gene-level associations under two-(left) and three-mechanism (right) architectures. **b**, Partition recovery accuracy (Macro-F1) of THREAD, MAGMA, and RolyPoly across varying gene set sizes (*G*), module numbers (*K*), and effect-size scales (Scale 1, weak; Scale 10, strong). **c**, Recovery of true genomic features (*x*_1_–*x*_5_) across Programs 1–3. Dot size and colour indicate −log_10_(FDR). **d**, Statistical power distributions (−log_10_(FDR)) under imbalanced mixture proportions (Cluster 1, 80%; Clusters 2 and 3, 10% each). Red dashed lines indicate the significance threshold. **e**, Inference purity (F1, top) and statistical power (recall, bottom) across background noise levels (25%–93%) for Bayesian-only (lFSR), frequentist-only (FGLS FDR), and the dual-intersection criterion. Error bars represent ±1 s.d. **f**, Alluvial diagram of signal classification across methods (TP, true positive; TN, true negative; FP, false positive; FN, false negative). **g**, Estimated effect sizes (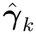, black circles) versus ground truth (red crosses) across features *x*_1_–*x*_5_ under Scale 1 (top) and Scale 10 (bottom).

We therefore examined architectures that differed along three biologically meaningful axes: the number of underlying programs, the strength of their cellular signals, and the proportion of non-informative genes. These spanned a homogeneous, tissue-wide architecture through risk distributed across increasingly many distinct programs; a weak and a markedly weaker per-subtype signal, both small relative to the simulated noise; and gene counts matching the range that real GWAS recover across traits of differing polygenicity (exact values, replication and the LD model in Methods). Each configuration was replicated 50 times to give more than a thousand datasets, analysed with THREAD and with two methods applicable to synthetic data, MAGMA and RolyPoly. By construction, these three axes isolate the features of real genetic architecture that a single-coefficient model handles poorly and that THREAD’s three components are built to address: risk split across multiple programs, a per-gene signal of varying reliability, and a majority of genes carrying no cell-subtype signal at all.

The three methods agreed when risk arose from a single homogeneous program (macro-F1 0.92–0.99) and diverged only once the architecture became heterogeneous— the regime in which a cellular decomposition is actually needed. With three or more programs, THREAD held a cell-subtype-resolved macro-F1 of 0.91–0.99 across every effect size and gene-set size, whereas MAGMA and RolyPoly fell to 0.57 and stayed there (Figure 2b). This was not a graded loss of power but a structural failure. A single-coefficient model fits one set of cell-subtype coefficients to the entire transcriptome; when different programs are driven by different cell subtypes, no single vector can represent them, and any signal the model does detect is propagated across all programs at once, producing systematic false positives. The plateau in Figure 2b is therefore not a matter of sample size but of misspecification: no amount of data recovers program-specific effects from a model that admits only one global coefficient vector. Figure 2c shows this at the level of individual programs—because MAGMA and RolyPoly return a single genome-wide set of significant cell subtypes and do not resolve genes into programs, that one verdict is necessarily correct where a cell subtype is causal and a false positive in every program where it is not, whereas THREAD assigns each cell subtype only to the program(s) that carry it, reproducing the sparse, block-structured truth. This is what the nonparametric mixture provides: it infers the program partition jointly with the coefficients and concentrates its posterior on the correct number of programs.

The same program resolution is what rescues rare cell populations, which are common in disease—infiltrating immune cells and transient developmental states among them. We generated architectures in which a dominant program accounted for 80% of genes and two minor programs for 10% each, the two minor programs deliberately differing in effect size. All methods recovered the dominant program, but the minor ones behaved according to their strength: the program whose cell-subtype effect was individually large was recovered by every method, even when its genes were pooled with the majority, whereas the weaker one fell below significance for MAGMA and RolyPoly—whose single estimate averages it into the dominant program—while THREAD, by isolating the program, retained it (Figure 2g). Dilution therefore bites precisely where a rare population is also weakly specified, the case most relevant to small disease-associated subpopulations.

A second feature of real data is the overwhelming excess of genes that participate in no disease-relevant program: in gene-level analyses only a minority carry detectable cell-subtype-linked heritability, the rest forming a largely non-informative background. We reproduced this by introducing an explicit null program whose coefficient vector was zero—its genes carrying no cell-subtype-linked heritability and generating GWAS effects from LD alone—and raising the proportion of null genes from 25% to 93%. Under this growing background, neither a purely Bayesian criterion (lFSR) nor a purely frequentist one (FDR) alone retained good operating characteristics, whereas requiring both held specificity near its ceiling at only a modest cost in sensitivity (Figure 2e). At the most extreme null proportion, THREAD routed most null-program genes to true negatives while MAGMA and RolyPoly converted a substantial fraction into false positives (Figure 2f). This motivates the dual-significance criterion used throughout, which calls a cell-subtype association credible only when it satisfies both lFSR < 0.05 and FDR < 0.05.

Detection, however, is only half of what the downstream analyses require: the effect sizes must also be trustworthy, and this is where the heteroscedastic likelihood becomes decisive. Because genes differ markedly in the precision with which their per-gene heritability can be estimated—each gene’s sampling variance is fixed by its own local LD (Methods)—treating every gene as equally reliable would bias the estimated coefficients. Weighting each gene by the precision of its own estimate instead recovered the generating coefficients without appreciable bias across both effect-size levels and all programs (Figure 2g). Because the Bayesian posterior mean is additionally shrunk toward zero by the Lasso prior and underestimates the true coefficient, we read effect sizes from the frequentist fit; the same prior withheld significance from a non-causal cell subtype even when it was strongly collinear with a causal one. We therefore report the dual-significant frequentist coefficient as the final effect size—the natural weight for the cell-subtype gene scores used in the applications that follow.

These simulations establish that when genetic risk is organized into multiple cellular programs, a pooled, single-coefficient model does not merely underperform but misrepresents the architecture: it collapses program-specific signal into one global effect, averages away rare cell populations, and inflates false positives once most genes are null. By jointly inferring program membership, modeling the gene-specific uncertainty of each heritability estimate, and controlling false discoveries under heavy null backgrounds, THREAD recovers the latent architecture of multicellular programs from which those cell-subtype associations arise.

### THREAD identifies disease-associated cell subtypes and functional modules in Alzheimer’s disease

We first applied THREAD to late-onset Alzheimer’s disease (AD), integrating publicly available GWAS summary statistics^8^ with a single-nucleus RNA-seq (snRNA-seq) atlas comprising 11,884 nuclei and 31 cell subtypes across six lineages from 6 AD and 6 control donors. This atlas profiles the human entorhinal cortex, one of the earliest brain regions affected by AD pathology^9^ (Fig. 3a). The snRNA-seq dataset is highly dominated by oligodendrocytes (57%) and astrocytes (17%), while microglia represent only ~3% of the captured nuclei (Fig. 3b), a cellular distribution that inherently constrains the statistical resolution available for prioritizing less abundant lineages, such as microglia. After standard preprocessing (Methods), THREAD partitioned the heritable risk of AD into six stable gene programs (C1–C6), containing 3,532, 3,900, 1,685, 185, 675, and 27 genes, respectively.

**Figure 3.**
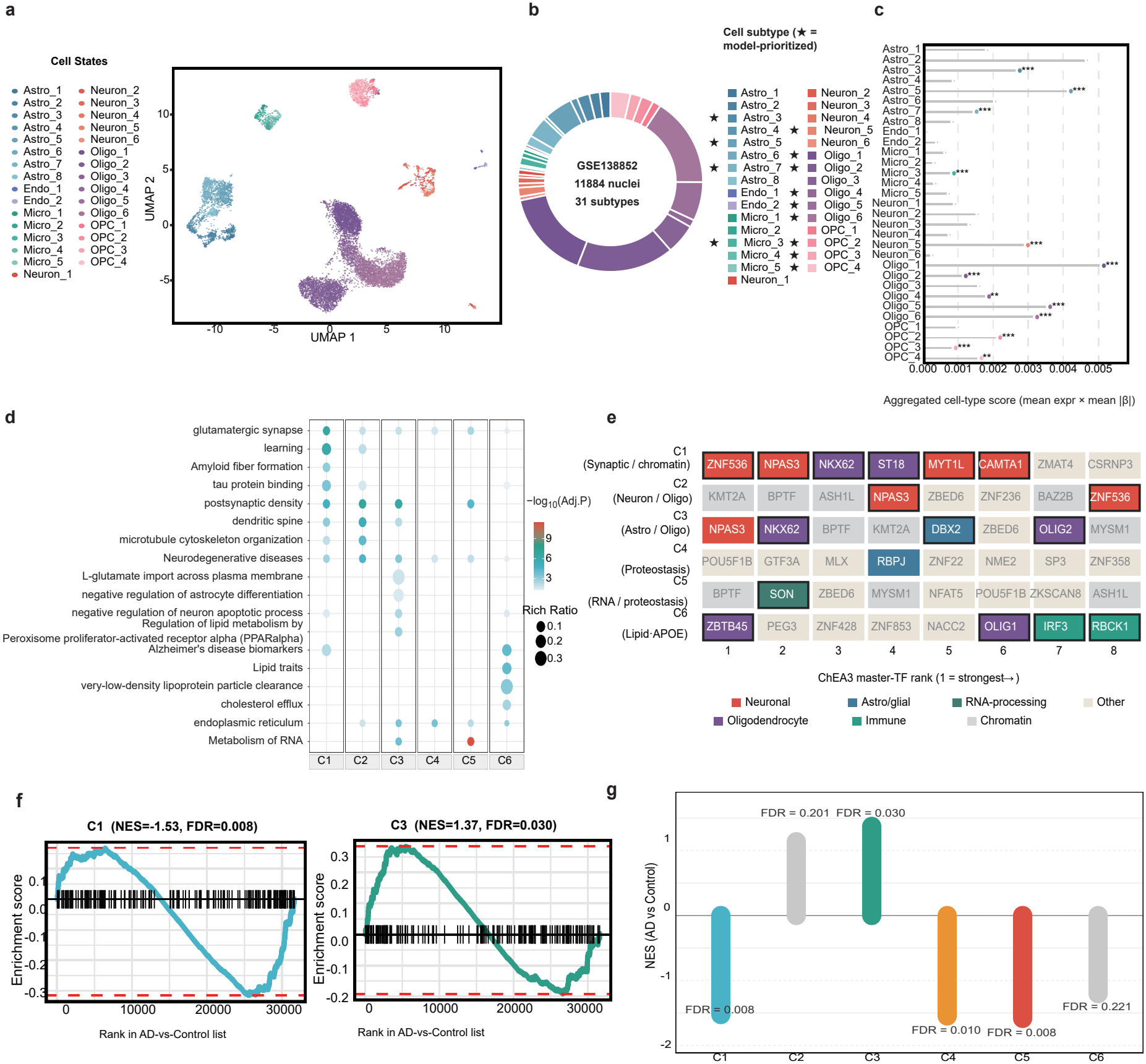
Dissection of Alzheimer’s disease (AD) heritability across entorhinal cortex cell states. **a**, UMAP of 11,884 nuclei from the human entorhinal cortex (GSE138852) across 31 subtypes. **b**, Nuclei proportions across the 31 subtypes; stars (*) indicate the 13 THREAD-prioritized subtypes. **c**, Aggregated cell-type association scores (mean expressionxmean 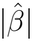). Asterisks denote dual-track significance (^*^*q* < 0.05, ^**^*q* < 0.01, ^***^*q* < 0.001). **d**, GO and Reactome pathway enrichment across AD modules C1–C6. Dot size, rich ratio; colour, −log_10_(adjusted *P*). **e**, Top eight master transcription factors (TFs) per module ranked by ChEA3, coloured by lineage. **f**, <monospace>fgsea </monospace>enrichment plots in the external GSE109887 cohort (*n* = 46 AD vs. *n* = 32 controls) for modules C1 (left) and C3 (right). **g**, Normalized enrichment scores (NES, AD vs. Control) and adjusted FDR values across modules C1–C6 in GSE109887.

Beyond simply nominating a set of risk-enriched cell states, THREAD assigned these cell states to distinct programs, each aggregating multiple significant cell states that may underlie a separable mechanistic axis of AD heritability. Thirteen of the 31 cell cell states passed the dual-significance criterion (Methods), drawn mainly from the oligodendrocyte, astrocyte and OPC lineages, with a neuronal (Neuron-5) and a microglial (Micro-3) subtype also selected (Fig. 3C and Supplementary Table). Consistent with the atlas of Grubman et al.^9^, in which astrocytic and oligodendrocytic states harboured most AD-associated transcriptomic change in this region, a shared oligodendroglial–astrocytic backbone (Oligo-1, Oligo-6 and Astro-7) recurred across nearly every program. Crucially, however, THREAD resolved distinct cell-type associations for each program (Supplementary Table). We designated C1 as a neuro-axonal program (“Neuron / Oligo”) due to its dual prioritization of neuronal and oligodendroglial lineages, and C3 as a glial metabolic program (“Astro / Oligo”) based on its astrocytic enrichment. C2 was labeled a synaptic-regulatory program (“Synaptic / Chromatin”) owing to its focus on chromatin modifiers. C4 and C5 were defined as housekeeping programs implicated in protein and RNA homeostatic machinery (“Proteostasis” and “RNA / Proteostasis”, respectively), while C6 was classified as a cell-type-agnostic lipid program (“Lipid-APOE”).

The two cell-subtype-specific programs, C1 and C3, represent biologically distinct processes corroborated by convergent pathway and upstream-regulator evidence (Fig. 3d,e). Program C1 (Neuron / Oligo) was strongly enriched for glutamatergic synapse (*q* = 2.6 × 10^−7^) and learning terms alongside classical AD pathology, such as tau-protein binding and amyloid-fibre formation (Fig. 3d). Its top-ranked genes spanned critical neuro-synaptic effectors (*PCDH*9, *DLG*2, *NRXN* 1, *NLGN* 1, *ERBB*4, *NPAS*3) together with myelin-related/oligodendrocyte-lineage genes (*MBP, QKI, LINGO*1, *CLDN* 11; Supplementary Table). Upstream regulator analysis (ChEA3; Supplementary Table) revealed a coordinated neuro-glial regulatory axis, combining neuronal transcription factors (*ZNF* 536, *NPAS*3, *MY T* 1*L*^10^) with oligodendrocyte-lineage factors (*NKX*6-2, *ST* 18)^11^ (Fig. 3e). This convergence implicates a coordinated pathogenic program operating at the neuron–oligodendrocyte interface. Program C3 (“Astro / Oligo”) was enriched for astrocyte differentiation, glutamate transport, and cholesterol biology. This program was anchored by the genome-wide-significant risk gene *CLU* ^12^, the reactive astrogliosis marker *GFAP*, and a suite of cholesterol regulators (*SREBF* 2, *NPC*1, *FDFT* 1, *PSAP*) that mediate lipoprotein and sterol metabolism. Additionally, C3 captured the astrocytic glutamate transporters *SLC*1*A*2/EAAT2 and *SLC*1*A*3/EAAT1; the AD-associated downregulation of these transporters is known to drive excitotoxic injury^13^ (Fig. 3d and Supplementary Table). Analysis of its upstream regulators highlighted the glial-lineage transcription factors *OLIG*2 and *NKX*6-2^14^ (Fig. 3e), recapitulating the astrocyte–oligodendrocyte transcriptional continuum characteristic of the degenerating brain.

Program C6 (Lipid-APOE; 27 genes) exhibited no cell-subtype expression specificity but was heavily enriched for AD GWAS-catalogue terms (*q* = 7.3 × 10^−5^) and systemic lipid traits, mapping directly to lipoprotein-clearance and cholesterol-efflux networks (Fig. 3d). The program was anchored by core genes such as *APOE, APOC*1, and *TOMM* 40, with *ZBTB*45 and the innate-immune regulator *IRF* 3 identified as its leading inferred upstream regulators (Fig. 3e). By grouping these functionally convergent lipid-transport genes into a single distinct program, THREAD recovered a compact, cell-type-agnostic risk axis that mirrors the cross-lineage lipid biology of the *APOE* locus^15^—signal that would be diluted when risk is resolved through cellular specificity alone.

The remaining programs resolved complementary, yet equally vital, pathological layers. Program C2 (Synaptic / Chromatin) prioritized excitatory neurons alongside glial subtypes but was uniquely distinguished by upstream chromatin remodellers (*KMT* 2*A, BPTF, ASH*1*L, BAZ*2*B*; Fig. 3e) and synaptic risk genes (*GRIA*2, *CNTNAP* 2), pointing to epigenetic dysregulation of synaptic plasticity. Programs C4 (Proteostasis) and C5 (RNA / Proteostasis) were dominated by broadly expressed cellular machinery. C4 was enriched for ubiquitin-proteasome and ER-associated degradation pathways, containing key proteostatic genes (*RPL*35, *FAU, V CP, UBE*2*K, UBE*2*N*), while C5 was enriched for RNA splicing and processing machinery (*MALAT* 1, *SON, SRSF* 2, *HNRNPH*1, *TRA*2*B*; Supplementary Table). These programs reflect the systemic collapse of protein and RNA homeostatic networks characteristic of the ageing brain (Supplementary Table).

To test whether these genetically defined programs reflect transcriptional perturbations beyond the discovery cohort without relying on single-gene selection, we analyzed an independent bulk microarray dataset comprising 46 AD cases and 32 controls^16^. Genes were ranked by the limma moderated *t*-statistic^17^, followed by an GSEA analysis using the top 200 genes from each program^18^(Methods; Fig. 3g).

Four of the six modules reached statistical significance (FDR < 0.05). The two cell-subtype-specific programs C1 and C3 shifted in biologically opposite and expected directions. Specifically, the C1 program (Neuron/Oligo) was coordinately downregulated (NES = −1.53, FDR = 0.008), consistent with the synaptic loss central to AD^19^. Conversely, the glial C3 program was significantly upregulated (NES = +1.37, FDR = 0.030, Figure 3f). The leading edge genes of C3 included *GFAP, SLC1A3, CLU*, and the cholesterol regulators *SREBF2* and *NPC1*, reproducing a canonical reactive-astrogliosis profile^13,20^. Programs C4 and C5 accounted for the remaining enrichments, reflecting broadly expressed translational and metabolic machinery that is consistent with the expression-weighted gene score (see Methods). This coupled axis of synaptic decline and glial activation represents a canonical AD transcriptomic signature^21^, recovered here anchored to genetically defined modules rather than post-hoc differential expression.By contrast, the cell-type-agnostic C6 program showed no transcriptional enrichment (FDR = 0.22). This lack of enrichment is expected for a small, primarily genetically driven module that evades detection in bulk tissue expression, given that only 19 of its 27 genes mapped.

The defining feature of the Alzheimer’s architecture is therefore its heterogeneity: risk that reads as broadly glial at the lineage level resolves into mechanistically distinct programs—neuro-axonal and astrocytic-homeostatic—alongside a cell-type-agnostic lipid core, a structure a single lineage-level enrichment would collapse into one glial signal.

### THREAD resolves shared and disease-specific genetic programs across schizophrenia and bipolar disorder

Unlike AD, with its focal pathology, psychiatric disorders lack discrete lesions and arise from highly polygenic risk distributed across the cortex. Schizophrenia (SCZ) and bipolar disorder (BIP) share a high degree of common-variant heritability^22^ yet are clinically distinct, and separating the cellular programs common to both from those specific to each remains open—a setting that tests whether decomposing risk into multiple programs can resolve shared from disorder-specific biology. We applied THREAD to each disorder using GWAS summary statistics^22,23^ and a large single-nucleus atlas of the human prefrontal cortex (GSE254569; 21 cell subtypes across seven major lineages) spanning control individuals and multiple psychiatric diagnoses, including SCZ, BIP, major depressive disorder and schizoaffective disorder, fitting the model independently for each (Methods).

The decomposition resolved both disorders into a neuronal core shared between them and an extension specific to schizophrenia. The model converged to five modules in SCZ (C1–C5; 8,013/5,994/2,235/372/23 genes) and four in BIP (C1–C4; 8,861/5,143/2,156/392 genes), prioritizing eight cell subtypes in SCZ and five in BIP (Figure 4a). In contrast to the glia-dominated architecture of AD, both converged on a neuronal core—upper-layer and deep-layer excitatory neurons (Exc-L2-3, Exc-L4-6-1), parvalbumin and VIP interneurons (In-PVALB-Ba, In-VIP) and protoplasmic astrocytes (Astro-PP)—recapitulating the known enrichment of psychiatric heritability in cortical excitatory neurons and interneurons^22,24,25^, including the vulnerability of prefrontal parvalbumin interneurons^26^, and confirming that the prioritized subtypes are disease-appropriate rather than a fixed signature. The two disorders exhibited substantial overlap while retaining disorder-specific components: all five BIP subtypes were also prioritized in SCZ, which additionally recruited RELN- and LAMP5-expressing interneurons (In-RELN, In-LAMP5-1^27^) and mid-layer excitatory neurons (Exc-L3-5). Because each module carried a distinct combination of these subtypes, the shared core and the SCZ-specific extension emerge as separable programs rather than one aggregate neuronal signal.

**Figure 4.**
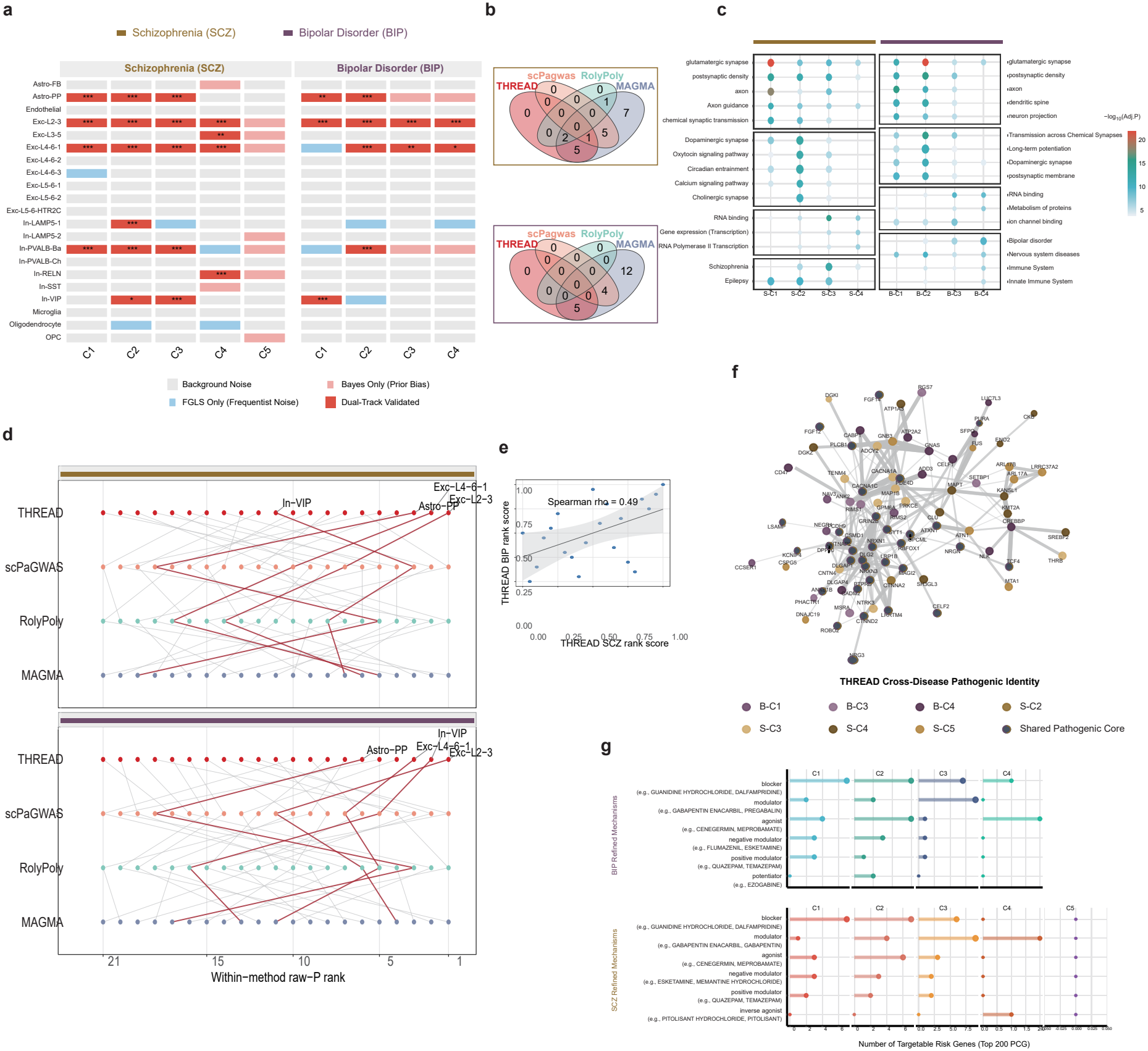
Comparative dissection of schizophrenia (SCZ) and bipolar disorder (BIP). **a**, Cell-subtype association heatmaps across 21 prefrontal cortex states for SCZ (C1–C5, left) and BIP (C1–C4, right), categorized into background noise, FGLS-only, Bayes-only, and dual-track validated calls (^*^*q* < 0.05, ^**^*q* < 0.01, ^***^*q* < 0.001). **b**, Venn diagrams of prioritized cortical cell states across methods in SCZ (top) and BIP (bottom). **c**, Pathway enrichment profiles (−log_10_(adjusted *P*)) for SCZ (S-C1 to S-C4) and BIP (B-C1 to B-C4) modules. **d**, Within-method raw *P*-value rank trajectories across methods in SCZ (top) and BIP (bottom). Red lines highlight key shared cortical subtypes. **e**, Cross-disorder correlation of THREAD cell-state rank scores between SCZ and BIP (Spearman’s *ρ* = 0.49; shaded band, 95% CI). **f**, Protein–protein interaction network of top module driver genes, highlighting the Shared Pathogenic Core (dark grey) and module-specific subnetworks. **g**, Mechanism-of-action stratification of targetable risk genes (top 200 coding genes per module) for BIP (top) and SCZ (bottom).

This pattern of overlap and divergence is what distinguishes THREAD from existing methods on identical inputs (Figure 4b): MAGMA returned widespread enrichment across nearly all subtypes in both disorders—biologically informative but too diffuse to separate shared from specific components; RolyPoly, the single-component model THREAD generalizes, was more conservative and called no subtype significant at the same threshold; and scPagwas returned intermediate sets (six in SCZ, four in BIP). Only THREAD returned focused subtype sets that retained the canonical excitatory and interneuron signals while exposing the shared-versus-specific structure. Quantifying this overlap directly, the relative prioritization of cell subtypes was positively correlated between SCZ and BIP (Spearman *ρ* = 0.49; Figure 4e), supporting a common neuronal liability, yet several subtypes departed from the diagonal—Astro-PP and deep-layer excitatory neurons (Exc-L4-6-1) weighted toward SCZ, upper-layer excitatory neurons (Exc-L2-3) and VIP interneurons (In-VIP) toward BIP —marking where the two architectures diverge.

The same pattern of overlap and divergence was evident at the molecular level (Figure 4c–f). Across both disorders, module genes were enriched for synaptic and axonal programs (glutamatergic synapse, postsynaptic density, dendritic spine, axon guidance) and for neurotransmission pathways (dopaminergic synapse, calcium signaling, long-term potentiation), consistent with disrupted excitatory–inhibitory balance^24^, while disease-ontology terms directly recovered both disorders. A protein–protein interaction network of the top module genes (STRING, combined score > 0.4)^28^ placed cross-disorder risk genes—the calcium-channel gene *CACNA1C* ^29^, the NMDA-receptor subunit *GRIN2B*, and the synaptic genes *NRXN1* and *TCF4* —at its centre, with disorder-biased genes in the periphery (Figure 4f), mirroring the cellular structure at the molecular scale. Finally, a fraction of the prioritized risk genes were targets of clinically used neuropsychiatric drugs spanning dopaminergic, glutamatergic (NMDA-receptor) and calcium-channel– modulating compounds (DGIdb; Figure 4g); because these targets were distributed across modules rather than concentrated in one, the decomposition offers a basis for module-stratified therapeutic repurposing.

For schizophrenia and bipolar disorder, the contribution is organizational: the decomposition resolves two genetically correlated disorders into a neuronal liability they share and an interneuron extension specific to schizophrenia, locating their overlap and their divergence in particular cell populations rather than in a single genetic-correlation coefficient.

### THREAD decomposes colorectal-cancer genetic risk into tumor-intrinsic and microenvironmental programs

Unlike psychiatric disorders, where risk is distributed across populations within one tissue, solid tumors are organized around a division between malignant compartment and the surrounding tumor microenvironment (TME), and whether inherited susceptibility acts through tumor-intrinsic mechanisms, the microenvironment, or both is a central question in cancer genetics. Colorectal cancer (CRC) is well suited to address it: its epithelium and TME are extensively characterized, and its tumors are stratified by the consensus molecular subtype (CMS) classification—a transcriptome-based taxonomy separating canonical epithelial (CMS2) from metabolic (CMS3), immune/microsatellite-unstable (CMS1) and mesenchymal/stromal (CMS4) tumors^30^. We applied THREAD to CRC GWAS summary statistics^31^ with a single-cell atlas of colorectal tumors and normal colon (GSE132465)^32^, in which malignant cells are labeled by CMS class alongside non-malignant epithelial, immune and stromal populations (Figure 5a); we focused on the tumor compartment, analyzing normal colon in parallel.

**Figure 5.**
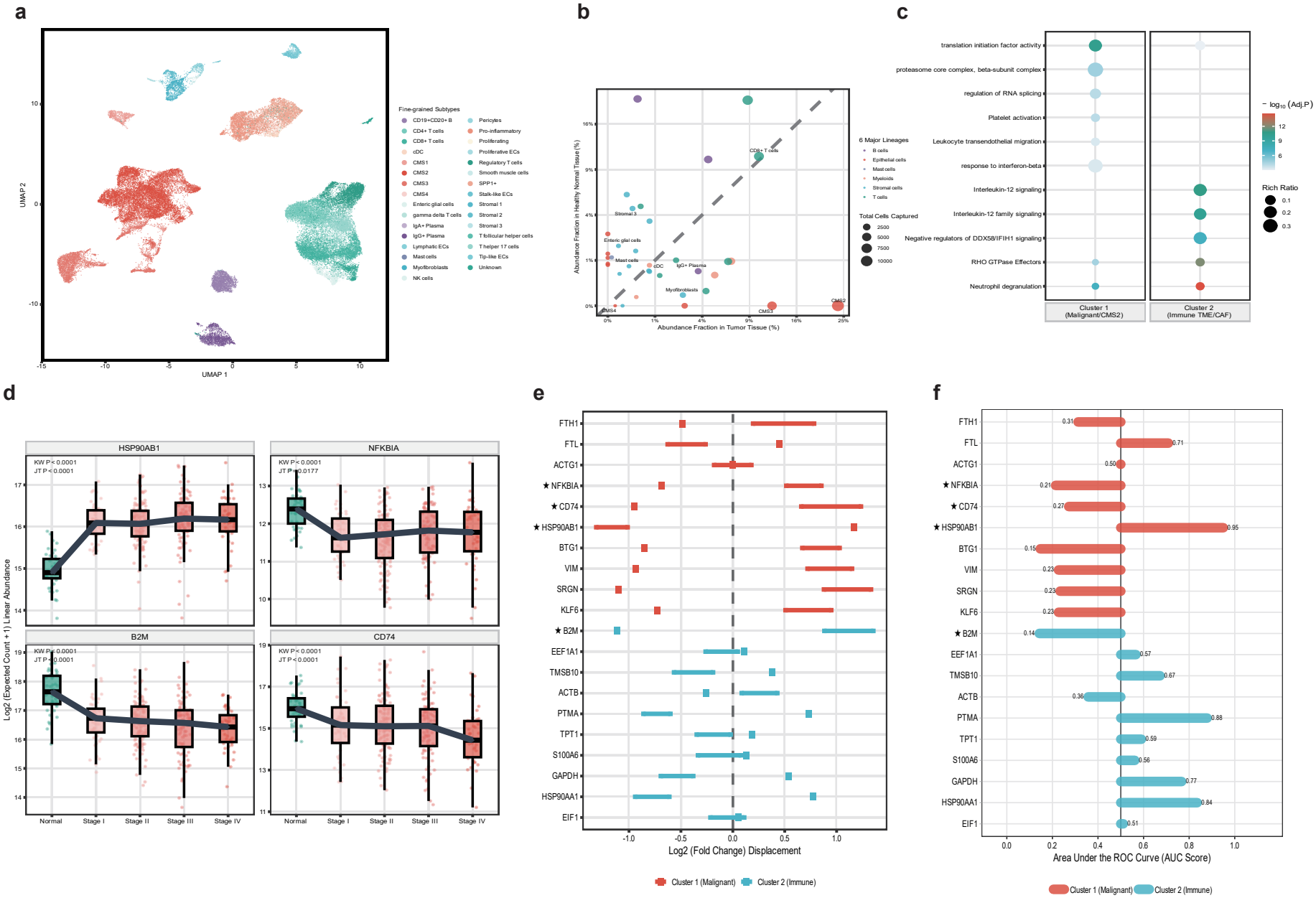
Deconvolution of colorectal cancer (CRC) heritability. **a**, UMAP of CRC single-cell transcriptomes (GSE132465) across 31 subtypes. **b**, Cellular abundance (%) in normal colon mucosa versus CRC tumour tissue across six major lineages. Dot size indicates total captured cells. **c**, Pathway enrichment comparison between Cluster 1 (Malignant/CMS2) and Cluster 2 (Immune TME/CAF). **d**, Stage-stratified expression (log_2_(expected count+1)) in TCGA-COAD across Normal and Stages I–IV for *HSP90AB1, NFKBIA, B2M*, and *CD74. P* values from Kruskal–Wallis (KW) and two-sided Jonckheere– Terpstra (JT) tests. Center lines, medians; boxes, IQR; whiskers, 1.5 × IQR. **e**, Tumour-versus-normal log_2_(fold change) displacement in TCGA-COAD for the top 10 genes in Cluster 1 (red) and Cluster 2 (blue). Stars (*) denote genes shown in **d. f**, Receiver operating characteristic AUC scores for discriminating tumour from normal tissue in TCGA-COAD across the top 20 module genes.

The decomposition separated CRC heritability into two programs along exactly this axis. The model converged to two modules: C1 was associated exclusively with the canonical epithelial subtype CMS2, whereas C2 was associated with a diverse immune and stromal collection—CD8^+^ T cells, conventional dendritic cells, IgG^+^ plasma cells, mast cells, myofibroblasts, Stromal-3 and enteric glia—together with the CMS3 and CMS4 malignant subtypes, the mesenchymal CMS4 grouping with fibroblast-rich stroma as expected from its stromal infiltration^30^. Rather than a single enriched population, these two modules represent a tumor-cell-intrinsic epithelial program (C1) and a microenvironmental immune–stromal program (C2).

Both the cellular composition and the biology of these programs supported the division. Relative to normal colon, tumors expanded their malignant epithelium—most strongly CMS2—alongside stromal and myeloid populations, and the populations with the largest abundance changes (CMS2 and multiple stromal populations) were among the prioritized subtypes (Figure 5b), so the modules track the cellular reorganization of malignant transformation rather than transcriptional variation alone. Functionally (Figure 5c), the epithelial program was dominated by biosynthetic and proteostatic processes—translation-initiation-factor activity (*q* = 4.7 × 10^−11^), the proteasome core, and RNA-splicing regulation—matching the protein-synthesis and quality-control demands of WNT/MYC-driven CMS2 tumors^30^, whereas the microenvironmental program was enriched for innate-immune, cytokine and cytoskeletal processes: neutrophil degranulation (*q* = 2.3 × 10^−15^), a feature of tumor-associated neutrophils in CRC^33^; interleukin-12 signaling (*q* = 1.5 × 10^−11^), linking its dendritic-cell and T-cell populations; and RHO-GTPase effector signaling (*q* = 1.1 × 10^−12^), consistent with the myofibroblast and mesenchymal (CMS4) populations^34^. The two programs are therefore polarized between proliferative-epithelial and immune–stromal-remodeling biology.

To ask whether these programs reflect clinically meaningful progression, we tracked their top-ranked genes across pathological stage in an independent TCGA-COAD cohort (Jonckheere–Terpstra trend test; Figure 5d–f). Thirteen of the 20 highest-ranked genes showed significant monotonic stage trends (*P* < 0.05), even though their ranking made no use of stage. Because the within-module score is expression-weighted, broadly expressed housekeeping genes also rank among the top (Methods); we therefore treat this ranking as a permissive candidate set and draw the conclusions below from genes with specific, directionally coherent changes rather than from the highest-ranked gene alone. These trends were directionally coherent: *HSP90AB1* separated tumor from normal tissue (AUC= 0.95) and rose with stage (*P* < 0.0001), matching the pro-proliferative, prognostically adverse role of HSP90 in CRC^35,36^, while antigen-presentation components fell—*B2M*, whose loss is prognostic for nodal metastasis^37^, and *CD74* decreased with stage (*P* < 0.0001), as did the NF-*κ*B inhibitor *NFKBIA* (*P* = 0.0177) Progression was thus marked by rising proliferative capacity and eroding antigen presentation, a recognized route to immune evasion^38^, linking the programs inferred from inherited risk to observable tumor evolution.

The colorectal result places inherited susceptibility within a compartment structure: rather than acting through malignant cells alone, risk divides between a tumor-intrinsic epithelial program and the immune–stromal microenvironment—a partition that does not stay static, with proliferation rising and antigen presentation eroding across pathological stage as the disease advances.

### THREAD identifies two multicellular breast-cancer risk programs that recapitulate subtype structure and show time-dependent prognostic associations

Breast cancer comprises molecularly and cellularly heterogeneous tumors whose clinical classification is commonly summarized by bulk tumor subtypes, although these labels do not directly resolve the multicellular malignant, immune and stromal programs from which they emerge^39^. We therefore asked whether inherited breast-cancer risk could be mapped onto such multicellular programs and whether the resulting programs were reproducible at the patient level. THREAD was applied to breast-cancer GWAS summary statistics^40^ together with a single-cell atlas of ER^+^, HER2^+^ and triple-negative breast cancers (GSE176078; ~98,000 cells across nine major lineages)^39^ (Figure 6a–c).

**Figure 6.**
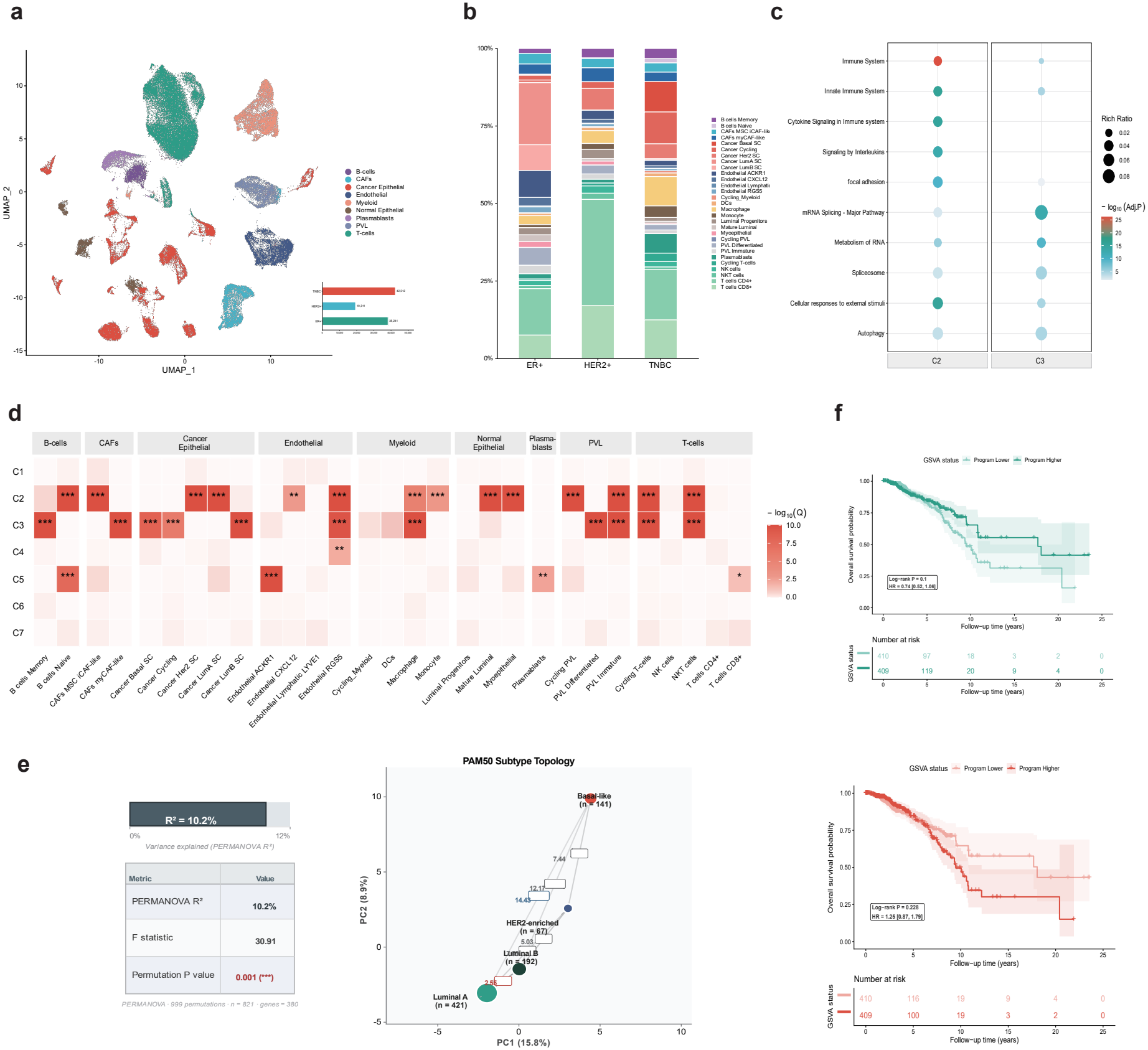
Multi-compartment resolution of breast cancer (BC) heritability. **a**, UMAP of the BC single-cell atlas across nine major lineages, with inset bar plot showing cell counts by receptor status (ER^+^, HER2^+^, TNBC). **b**, Cellular composition of 29 fine-grained cell states across ER^+^, HER2^+^, and TNBC tumours. **c**, Pathway enrichment profiles for BC modules C2 and C3. **d**, Dual-track association heatmap (-log_10_(*q*)) across seven BC modules (C1–C7) and 29 cell states (^*^*q* < 0.05, ^**^*q* < 0.01, ^***^*q* < 0.001). **e**, Projection of bulk breast tumours onto the DPM latent space (PC1 vs. PC2) across PAM50 subtypes: Basal-like (*n* = 159), HER2-enriched (*n* = 76), Luminal A (*n* = 490), and Luminal B (*n* = 213). Boxed values show pairwise centroid distances; left table shows PERMANOVA results (999 permutations). **f**, Kaplan–Meier overall survival curves and risk tables stratified by median expression of module C2 (top) and C3 (bottom). Log-rank *P* values, Cox hazard ratios (HR), and 95% CIs (shaded bands) are shown.

Heritable risk was concentrated in two cross-cell-type programs, C2 and C3, that differed in both their malignant-cell composition and functional annotation. Neither program was restricted to a single cellular compartment: both included immune populations (including macrophages, NKT cells and cycling T cells), stromal populations (CAFs, endothelial and perivascular-like cells) and malignant populations. Their malignant components nevertheless differed, with C2 including luminal-associated malignant states (LumA-SC and HER2-SC) and C3 including basal and cycling malignant states (basal-SC and cycling cancer cells)^39,41^. Functional enrichment further distinguished the two programs (Figure 6c). C2 was enriched for innate immune, cytokine and interleukin signaling together with focal-adhesion pathways, defining an immune–inflammatory program, whereas C3 was enriched for RNA processing and pre-mRNA splicing together with autophagy-related processes, consistent with the dependence of basal-like tumors on RNA-processing machinery and cellular stress-adaptation pathways^42,43^.

We next tested whether these programs reproduced in an independent patient cohort using TCGA-BRCA primary tumors. Projection of the C2/C3 signatures into TCGA yielded a 380-gene patient-level signature space in 821 tumors. The first two principal components captured 24.7% of the variance (PC1, 15.8%; PC2, 8.9%) and revealed an ordered PAM50 topology, with Basal-like and Luminal A tumors showing the greatest separation and Luminal A and Luminal B remaining the closest pair (Figure 6d). PAM50 subtype was associated with 10.2% of the multivariate variation in this signature space (PERMANOVA *R*^2^ = 0.102, *F* = 30.91, permutation *P* = 0.001). Pairwise effects were strongest between Basal-like and Luminal A tumors (*R*^2^ = 0.114) and weakest between Luminal A and Luminal B tumors (*R*^2^ = 0.029), indicating that the dominant structure reflected broad differences between molecular subtype classes rather than complete separation of adjacent subtypes. Because within-subtype dispersion also differed significantly (PERMDISP *F* = 55.52, *P* = 0.001), the PERMANOVA result was interpreted as evidence of subtype-associated multivariate structure rather than a pure centroid-separation effect.

Program-level scoring provided a complementary view of this structure. GSVA scores calculated from the mapped C2 and C3 signatures differed strongly across PAM50 subtypes^44,45^. C2 activity was higher in Basal-like and HER2-enriched tumors and lower in Luminal A and Luminal B tumors (Kruskal–Wallis *H* = 72.12, BH *q* = 3.0 × 10^−15^), whereas C3 showed the reciprocal pattern, with higher activity in Luminal A and Luminal B tumors (Kruskal–Wallis *H* = 66.94, BH *q* = 1.93 × 10^−14^). The corresponding rank-based effect sizes were *ϵ*^2^ = 0.085 for C2 and *ϵ*^2^ = 0.078 for C3. Pairwise comparisons indicated that these differences were driven predominantly by separation between the broader luminal and non-luminal subtype groups rather than by strong differences between Basal-like and HER2-enriched tumors or between Luminal A and Luminal B tumors. Thus, the patient-level activities of C2 and C3 reproduced a reciprocal axis of breast-cancer subtype heterogeneity without simply recapitulating the malignant-cell labels used to characterize the discovery cohort.

We next examined whether these continuous program activities were associated with patient outcome independently of PAM50 subtype. In 819 patients with available overall-survival information (120 deaths), PAM50-adjusted Cox models showed opposite average associations for the two programs: higher C2 activity was associated with a lower mortality hazard (HR per SD = 0.78, 95% CI 0.65–0.95, BH *q* = 0.0128), whereas higher C3 activity was associated with a higher mortality hazard (HR per SD = 1.34, 95% CI 1.10–1.62, BH *q* = 0.0058). Proportional-hazards diagnostics, however, indicated that the score effects were not constant over follow-up. Explicit time-varying coefficient models confirmed significant score-by-time interactions for both C2 and C3 (BH *q* = 0.029 for each; Figure 6e). During the first 1–2 years, the estimated effects were close to the null with wide confidence intervals. Thereafter, the programs progressively diverged: at 5 years, the estimated HR per SD was 0.73 (95% CI 0.59–0.89) for C2 and 1.44 (95% CI 1.18–1.77) for C3. The same directions persisted into later follow-up, although estimates at the longest follow-up times were supported by progressively smaller risk sets. We therefore interpret the conventional Cox estimates as follow-up-averaged associations and the time-varying analysis as evidence that the prognostic separation associated with C2 and C3 becomes more apparent during intermediate-to-late follow-up.

Together, these analyses map inherited breast-cancer risk onto two multicellular programs with distinct functional profiles that reproduce as patient-level transcriptomic structure in an independent cohort. C2 and C3 intersect with, but are not reducible to, the PAM50 taxonomy: they form reciprocal subtype-associated activity gradients and retain opposite associations with outcome after adjustment for PAM50. Rather than defining fixed high- and low-risk patient classes, the results support continuous and time-dependent relationships between these multicellular programs and breast-cancer outcome, linking the cell-state organization of inherited risk to both molecular heterogeneity and longer-term disease course.

## Methods

### GWAS summary statistics and SNP-to-gene mapping

We obtained GWAS summary statistics for the analyzed complex traits and applied stringent quality control. SNPs located on sex chromosomes or with a minor allele frequency (MAF) < 0.01 were excluded. Furthermore, variants within the Major Histocompatibility Complex (MHC) region were strictly removed to prevent its extensive, long-range linkage disequilibrium (LD) from confounding polygenic parameter inference.

To systematically construct gene-level genetic associations, we mapped the retained SNPs to target genes based on genomic proximity. Following established evidence that the majority of eQTLs consistently reside near promoters, we adopted a conservative 10-kb window centered on the TSS of each gene. While incorporating distal functional variants via chromatin conformation maps is biologically plausible, doing so inherently biases the analysis toward well-characterized lineages; thus, strictly proximity-based assignments were utilized. SNPs mapping to the overlapping windows of multiple genes were duplicated and treated as independent SNP-gene pairs. Given the scale of highly polygenic models, this standard procedure does not significantly bias parameter inference.

The resulting gene-level heritability estimates were strongly right-skewed, with a small number of genes exhibiting substantially larger values than the remainder. To reduce skewness and limit the influence of extreme observations, the estimates were log-transformed prior to model fitting. Because numerical estimation occasionally produced values extremely close to zero, a small constant (10^−8^) was added before transformation when necessary. The transformed response was subsequently standardized, yielding the response variable used throughout the analysis.

All analyses used genome build hg19/GRCh37. Gene coordinates were taken from the Ensembl GRCh37 gene annotation (release 87), and pairwise LD between variants was computed from a PLINK-processed 1000 Genomes Phase 3 European-ancestry reference panel.

### Single-cell expression reference and cell-subtype profiles

Quality-controlled single-cell or single-nucleus RNA-seq count matrices with author-provided cell-type and subtype annotations were used as expression references. For datasets distributed as raw count matrices, cells were first processed in Seurat^46^ using standard quality-control procedures, including filtering low-quality cells on the basis of detected genes and mitochondrial transcript content, followed by library-size normalization with the LogNormalize procedure. Datasets distributed with processed expression values were used as provided by the original authors.

To obtain stable cell-subtype-level expression profiles, normalized expression values were aggregated across cells sharing the same author-provided subtype annotation, generating pseudobulk mean-expression profiles for each subtype. The resulting gene-by-subtype matrix was log-transformed and standardized prior to model fitting, yielding an *N* x *P* expression-specificity matrix **X**, where *N* denotes genes and *P* denotes cell subtypes. The row **x**_*j*_ E ℝ^*P*^ represents the expression-specificity profile of gene *j* across all annotated cell subtypes.

THREAD relates gene-level genetic variance to the expression specificity of the cell populations defined in the reference atlas. Although all analyses here were performed and grouped into cell-subtype resolution for interpretation, the framework is agnostic to annotation depth and applies equally to any cellular resolution.

### THREAD model

THREAD decomposes phenotype-associated polygenic risk into interpretable multicellular programs by integrating GWAS summary statistics with single-cell expression profiles within a heteroscedastic Dirichlet process mixture framework. The construction starts from the polygenic model across *p* SNPs:

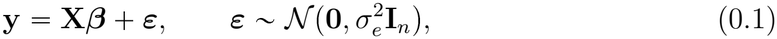

where **y** ∈ ℝ^*n*^ is the phenotype vector, **X** ∈ ℝ^*n×p*^ is the standardized genotype matrix across *p* SNPs, ***β*** is the vector of true SNP effects, and *n* is the GWAS sample size.

GWAS summary statistics are obtained from one-SNP-at-a-time regressions. Let **X**_*i*_ ∈ ℝ^*n*^ be the standardized genotype vector for SNP *i*, the marginal effect estimate is 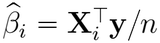. Collecting the estimates for all *p* SNPs gives 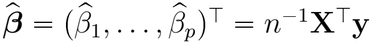, where **X** = (**X**_1_, …, **X**_*p*_).

Substituting the polygenic model (0.1) into the marginal estimators yields

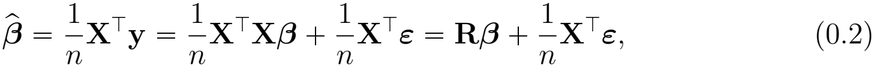

where **R** = *n*^−1^**X**^T^**X** is the LD matrix.

For the set of SNPs *S*_*j*_ assigned to gene *j*, Let 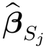 and 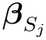 denote the vectors of marginal effect estimates and true GWAS effect sizes for the SNPs assigned to gene *j* respectively, and let **R**_*j*_ denote the local LD correlation matrix. We assign a random-effect prior to 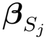:

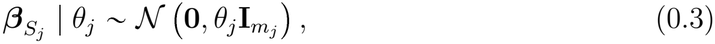

where *m*_*j*_ = /*S*_*j*_/ and *θ*_*j*_ is the per-SNP genetic variance attributable to gene *j*. Marginalizing over 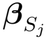 yields the gene-level distribution of GWAS marginal effect estimates for gene *j*:

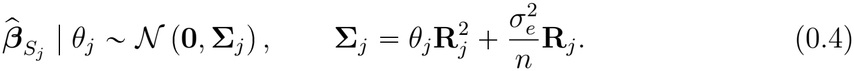

To connect GWAS summary statistics to single-cell expression while accommodating functional heterogeneity across genes, we allow the relationship between single-cell expression profiles and polygenic risk *θ*_*j*_ to vary across latent gene programs. Specifically, Let **s**_*j*_ = (*s*_*j*1_, …, *s*_*jP*_)^T^ ∈ ℝ^*P*^ denote the cell-type expression profile of gene *j*, and let *c*_*j*_ denote its latent program label. Conditional on *c*_*j*_ = *k*, we link the per-SNP effect-size variance of gene *j* to its expression profile through a program-specific linear model:

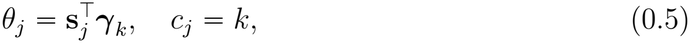

where ***γ***_*k*_ = (*γ*_*k*1_, …, *γ*_*kP*_)^T^ quantifies the contribution of each cell type to program *k*. Thus, genes assigned to the same program share a common mapping from cell-type expression to polygenic effect-size variance, whereas genes in different programs may be associated with distinct cell types.

To estimate the program-specific effects ***γ***_*k*_ from GWAS summary statistics, we transform the marginal effect estimates 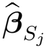 into a scalar gene-level response variable:

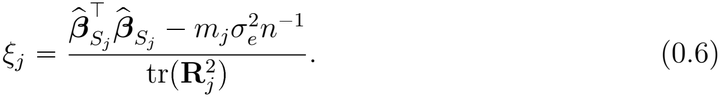

Notice that 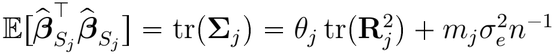, therefore *ξ*_*j*_ is consequently an unbiased estimator of the gene-level per-SNP effect-size variance *θ*_*j*_. We approximate the distribution of *ξ*_*j*_ by a Gaussian working likelihood matched to its exact first two moments:

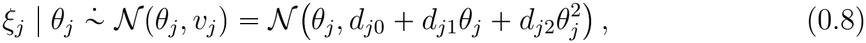

where 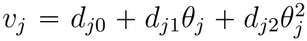 is the variance of *ξ*_*j*_, and the coefficients *d*_*j*0_, *d*_*j*1_, *d*_*j*2_ are functions of the local LD matrix **R**_*j*_, the GWAS sample size *n* and the residual variance 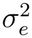. The detailed derivation of *v*_*j*_ are provided in the Supplementary Notes.

Combining (0.5) and (0.8) gives the program-specific heteroscedastic regression model

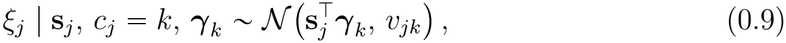

where 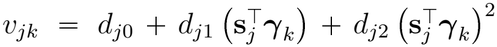. This construction preserves the gene-specific sampling uncertainty inherited from the GWAS summary statistics while allowing different groups of genes to exhibit different cell-type architectures.

To automatically infer the number of latent gene programs and estimate the underlying cellular effects ***γ***_*k*_ without predefining the cluster count, THREAD formulates this heteroscedastic regression as a Dirichlet process mixture model over genes.

Let 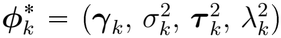 denote the coefficient vector of program *k* together with its associated parameters, The hierarchical Bayesian formulation is defined as:

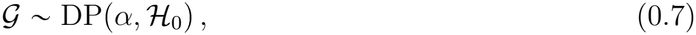

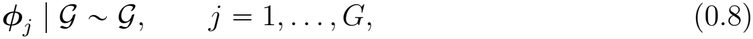

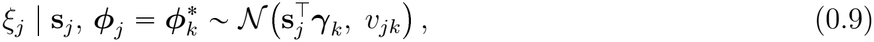

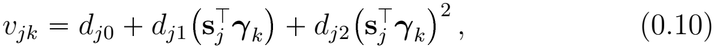

where the base distribution *H*_0_ assigns ***γ***_*k*_ a hierarchical Laplace prior in normal–exponential scale-mixture form:

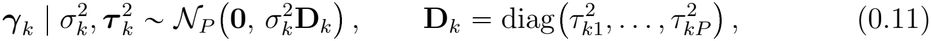

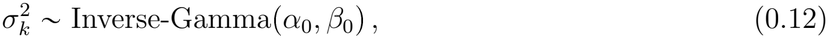

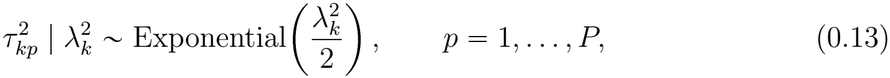

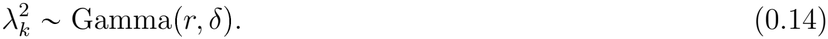

The clustering structure follows from the discreteness of *G*. Under the stick-breaking construction,

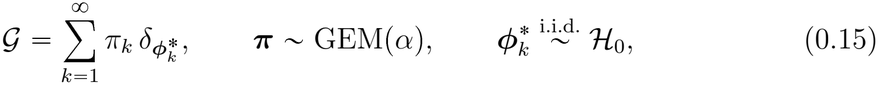

where ***π*** represents the infinite-dimensional mixture proportions drawn from a Griffiths–Engen–McClosk (GEM) distribution with concentration parameter *α*. Therefore introducing the latent label *c*_*j*_ / ***π*** ~ Categorical(*π*_1_, *π*_2_, …) gives 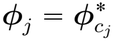, and equation (0.9) becomes

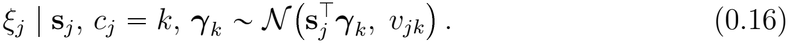

Genes sharing the same label are governed by a common coefficient vector ***γ***_*k*_ and constitute one latent module, whereas genes in different modules may be associated with distinct cell types. Posterior inference is performed by MCMC, the full sampling procedures are provided in the Supplementary Notes. The complete hyperprior hierarchy and all hyperparameter values are given in Supplementary Note and Supplementary Table.

### Prior-guided initialisation

Although THREAD are highly flexible, randomly initialised MCMC can suffer from long burn-in periods and local optima if initialized randomly. To accelerate chain convergence and provide a biologically informed starting partition, trait-associated genes were embedded in a pre-trained 200-dimensional Gene2vec co-expression space and clustered by *k*-means to generate initial module labels. For genes lacking Gene2vec representations, we imputed their initial labels using a KNN voting scheme based on their scaled single-cell expression profiles. This initialisation affects only the starting state and not the stationary distribution of the sampler.

### Posterior inference by MCMC

We approximate the posterior with a Markov chain Monte Carlo sampler that updates conjugate parameters by exact Gibbs sampling and the remaining parameters by tailored non-conjugate moves; the full conditional distributions and the complete sampling scheme are given in Supplementary Note 1. Module labels are updated with Neal’s Algorithm 8, which represents currently empty modules by a small number of auxiliary components proposed from the prior. The coefficient vectors ***γ***_*k*_ are non-conjugate—because the heteroscedastic variance *v*_*jk*_ depends on ***γ***_*k*_ through both the mean and the variance of the Gaussian kernel—and are sampled by stepping-out slice sampling, whereas the Bayesian-Lasso scale parameters retain closed-form conjugate updates. We ran *N*_iter_ iterations per trait, discarded the first *N*_burn_ as burn-in, and thinned the remainder; convergence was monitored from the stability of the sampled module count together with standard diagnostics (Supplementary Table).

### Point estimate of the module partition

Because the mixture likelihood is invariant to relabelling of the modules, individual MCMC samples of the labels are not directly comparable; we therefore summarise the partition through pairwise co-assignment, which is permutation-invariant. From the post-burn-in samples we formed a posterior similarity matrix (PSM) whose (*i, j*) entry is the posterior probability that genes *i* and *j* occupy the same module. We fixed the number of modules at the mode of the sampled module count along the chain, discarding transient micro-clusters (10 genes) as sampling noise, and then obtained the point-estimate partition by partitioning-around-medoids on the dissimilarity matrix **1** - PSM, selecting the final number of modules by the maximum average silhouette width. This partition defines the modules used in all subsequent analyses.

### Dual significance criterion for cell-subtype prioritisation

A module–cell-subtype association was reported only when it satisfied two independent criteria—one Bayesian, one frequentist—evaluated on the point-estimate partition. The first is a posterior criterion based on lFSR: aligning the coefficient samples to the point-estimate modules, we computed lFSR_*kp*_ = Pr(*γ*_*kp*_ 0 / data) for cell subtype *p* in module *k* and required lFSR_*kp*_ < 0.05. The second is a frequentist criterion based on the FDR: each module was refitted by feasible generalised least squares (FGLS)—iteratively reweighting genes by 1/*v*_*jk*_ until convergence and using heteroscedasticity-consistent (sandwich) standard errors—after which a one-sided test for positive enrichment (*H*_1_ : *γ*_*kp*_ > 0) was corrected across all module x cell-subtype comparisons by the Benjamini– Hochberg procedure. A cell subtype was called *dual-significant* for a module when both lFSR_*kp*_ < 0.05 and FDR_*kp*_ < 0.05. Because the Bayesian posterior mean is shrunk toward zero by the Lasso prior and is therefore biased for quantification, the FGLS estimate 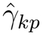—not the posterior mean—was reported as the effect size. cell subtypes that are jointly dual-significant within a module constitute a synergistic, multicellular program.

### Prioritisation of genes within modules

To nominate candidate molecular effectors of each program, we scored every gene by an expression-weighted combination of its module’s coefficients,

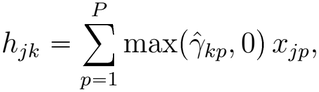

restricting the sum to positive coefficients so that the score reflects only positive contributions to genetic variance, and ranked genes within each module in descending order of *h*_*jk*_. This score is an expression-weighted prioritisation rather than a direct measure of disease relevance: because it scales with expression magnitude, broadly and highly expressed genes—including housekeeping genes that are abundant even within a disease-relevant module—tend to receive high ranks. We therefore treated the top-ranked genes (top 200 unless otherwise stated) as a permissive candidate set for set-level analysis rather than as individually validated targets, and drew no biological conclusion from the single highest-ranked gene. Downstream analyses were correspondingly set-based—pathway over-representation and, in the disease applications, <monospace>fgsea</monospace> over the module signature in an externally ranked cohort—so that no single housekeeping gene can drive a result; where individual genes were examined directly (e.g. the colorectal-cancer stage trends), the directionally coherent, stage-tracking signal was carried by specific disease-relevant genes rather than by the broadly expressed genes that the expression-weighted score also ranks highly.

### Integrated cell-subtype score

THREAD estimates, for every module *k*, a coefficient 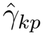 for each cell subtype *p* that quantifies its contribution to that module’s genetic variance. To summarise the overall importance of a cell subtype across the entire model—rather than its role within any single module—we defined an integrated cell-subtype score as the total absolute coefficient of that cell subtype across all *K* modules,

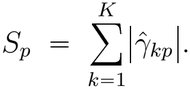

Absolute values are used because, unlike the within-module gene score—where negative coefficients are set to zero so that the score reflects only positive contributions to genetic variance—a negative coefficient still encodes a genuine association between a cell subtype and genetic variance and therefore contributes to that cell subtype’s overall effect. The score depends only on the fitted coefficients and deliberately not on a cell subtype’s average expression level, so that cell subtypes are ranked by the genetic weight the model assigns them rather than by transcriptional abundance; weighting by mean expression would otherwise inflate broadly and highly expressed populations irrespective of disease relevance, the cell-subtype analogue of the housekeeping-gene effect noted for the within-module gene score. (Dividing by *K* gives the mean absolute coefficient and yields an identical ranking.) cell subtypes were ordered by *S*_*p*_ and displayed as a lollipop plot (Figure 3c).

The score is a magnitude summary only; statistical prioritisation is carried entirely by the dual significance criterion. Accordingly, a cell subtype was marked as significant in this display solely when it was dual-significant—satisfying both the lFSR and the FDR criterion—in at least one module, with the marker graded by its strongest evidence across modules (the minimum FGLS FDR *q*-value, shown as one, two or three symbols for *q* < 0.05, *q* < 0.01 and *q* < 0.001), and cell subtypes that were not dual-significant in any module were de-emphasised regardless of their score.

### Functional annotation of prioritised programs

The top-ranked genes of each module (its *signature*) were annotated for biological function; full gene lists and outputs are provided in the Supplementary Information. Across all diseases, functional over-representation was assessed with KOBAS-i^47^ against KEGG, Reactome, KEGG DISEASE, Gene Ontology and the NHGRI-EBI GWAS Catalog, using the hypergeometric (Fisher’s exact) test with Benjamini–Hochberg control of the false discovery rate at 0.05. Disease-specific analyses were added as follows.

#### Upstream regulators (Alzheimer’s disease)

For each module, upstream transcription factors were inferred with ChEA3^48^; the eight top-ranked factors by integrated mean rank are reported, with complete rankings in Supplementary Table.

#### Protein interactions and druggability (schizophrenia and bipolar disorder)

The 15 highest-scoring protein-coding genes per module were pooled as input to a STRING^28^ protein–protein interaction network (combined-score threshold 0.4, coloured by module of origin), and druggable targets were annotated with DGIdb^49^ from the top 200 genes per module.

### Cell-subtype abundance comparison

For colorectal cancer, we related the prioritized programs to the cellular reorganization of the tissue by comparing cell-subtype composition between the normal and tumour compartments of the single-cell atlas. Within each compartment, every annotated subtype was expressed as its fraction of all captured cells (subtypes annotated as unknown were excluded), and the two compartments were compared subtype by subtype against the line of equal abundance, with the total number of cells captured per subtype indicating its sampling depth. This descriptive comparison was used to assess whether the subtypes prioritized from genetic risk coincided with those whose abundance shifted most between normal and malignant tissue (Figure 5b).

### Independent cohort validation

Prioritised programs were validated in independent cohorts using the modality appropriate to each disease; cohort accessions, sample sizes and quality control are listed in Supplementary Table, and exhaustive per-gene results in the Supplementary Information. Not every modality was applied to every disease.

#### Independent transcriptomic validation (Alzheimer’s disease)

To test whether the genetically defined modules are reproducibly perturbed beyond the discovery cohort, and to avoid any single-gene selection bias, we used gene set enrichment analysis in an independent bulk cohort that differs in brain region, modality and platform. All genes were ranked by the limma^17^ moderated *t*-statistic (case versus control), and the top 200 protein-coding signature genes of each module were tested for coordinated enrichment with fgsea^18^, reporting the normalised enrichment score, *P* and FDR per module; module-level enrichment curves are shown for the two core modules.

#### Clinical progression and diagnostic performance (colorectal cancer)

Top-ranked genes of the two leading modules were examined in the TCGA-COAD cohort. Stage-dependent expression was tested across pathological stages (Normal through Stage IV) with the non-parametric Jonckheere–Terpstra trend test, with the Kruskal–Wallis test reported alongside as an unordered control. Diagnostic separation of tumour from normal tissue was quantified for each gene by the area under the ROC curve derived from the Mann–Whitney statistic, and effect sizes were summarised as log_2_ fold changes with 95% confidence intervals from two-sample *t*-tests. The two genes per module shown in the main figure were selected for interpretability; the full set of twenty genes is reported in the Supplementary Information.

#### Molecular-subtype projection and survival (breast cancer)

The two prognostic programs were projected onto the external TCGA-BRCA cohort by scoring every tumour for each module signature with GSVA^44^ over the top-ranked module genes (Gaussian kernel for continuous bulk data). The variance in PAM50 intrinsic-subtype identity explained by the two signatures was quantified by PERMANOVA^50^. For survival analysis, patients were dichotomised at the median score of each signature separately and overall survival was compared by the log-rank test, with hazard ratios and 95% confidence intervals from univariable Cox proportional-hazards models.

### Simulation design

We designed simulations to evaluate the regimes in which decomposing polygenic risk into program-specific cell-type effects accurately recovers latent multicellular programs and heterogeneous risk architectures, compared with conventional approaches RolyPoly and MAGMA that model all trait-associated genes under a single uniform cellular architecture. Each simulated dataset contained *G* genes (*G* ∈ {500, 1000, 3000}) and *P* = 5 cell types. For gene *j* (*j* = 1, …, *G*), the cell-type expression vector **x**_*j*_ = (*x*_*j*1_, …, *x*_*jP*_)^T^ ∈ ℝ^5^ was constructed by drawing each element independently as *x*_*jp*_ = /*ϵ*_*jp*_/, where *ϵ*_*jp*_ ~ *N* (1, 0.5^2^). This element-wise independent sampling ensures that the gene expression matrix **X** ∈ ℝ^*G×P*^ carries no intrinsic program or modular structure; instead, all latent cellular architecture is introduced strictly through program-specific coefficient vectors ***γ***_*k*_ that map cell-type expression profile **x**_*j*_ to the gene-level effect-size variance *θ*_*j*_, defined as the per-SNP effect variance of variants assigned to gene *j*.

Each gene was allocated to one of *K*_0_ latent programs by an independent multinomial draw, where *c*_*j*_ ∈ {1, …, *K*_0_} records the program to which gene *j* belongs. The fixed mixing weights 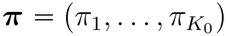, with *π*_*k*_ = Pr(*c*_*j*_ = *k*) and Σ_*k*_ *π*_*k*_ = 1, set the expected proportion of genes contributed by each program. Every program *k* was characterized by a sparse coefficient vector ***γ***_*k*_ ∈ ℝ^*P*^ whose non-zero entries designate the causal cell types of that program. Stacking these vectors row-wise gives the generating coefficient 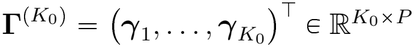, where the entry *γ* quantifies the contribution of cell type *p* within program *k*.

We evaluated three architectures. For *K*_0_ = 1, with ***π*** = (1), all genes shared the same coefficient vector,

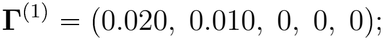

for *K*_0_ = 3, with ***π*** = (0.35, 0.35, 0.30),

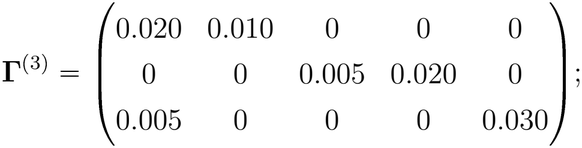

and for *K*_0_ = 5, with ***π*** = (0.35, 0.35, 0.10, 0.10, 0.10),

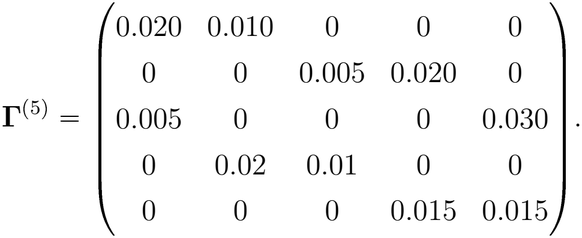

Every coefficient matrix was additionally rescaled by a factor *s* ∈ {1, 10}, giving two per-cell-type signal strengths that differ by an order of magnitude. Given the allocation *c*_*j*_, the gene-level effect-size variance was set as 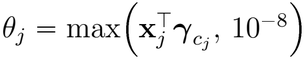, where the lower bound ensuring a positive-definite covariance in the sampling step.

For each gene *j*, we simulated *M* = 500 SNPs. The LD matrix **R** ∈ ℝ^*M×M*^ across these SNPs was generated under a first-order autoregressive structure with entries *R*_*ij*_ = *ρ*^l*i*-*j*l^ (*ρ* = 0.3). Assuming a GWAS sample size of *n* = 10,000 and a residual variance of 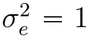, the SNP-level marginal effect estimates 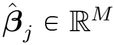 were drawn directly from the multivariate normal distribution:

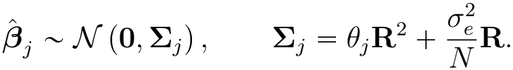

#### Benchmark grid and targeted evaluation regimes

The main benchmark evaluated 18 primary settings spanning all factorial combinations of program counts *K*_0_ ∈ {1, 3, 5}, total gene counts *G* ∈ {500, 1000, 3000}, and coefficient scale factors *s* ∈ {1, 10}, with 50 independent Monte Carlo replicates per setting (900 datasets in total). To evaluate model performance under realistic biological complexities beyond the balanced grid, we further constructed three targeted evaluation regimes:

i. **Rare-program regime**: Fixed at *G* = 3000 and scale factor *s* = 1, this setting introduced a highly imbalanced mixing proportion of (0.80, 0.10, 0.10) under the three-program matrix **Γ**^(3)^. It specifically evaluates whether low-frequency minor programs (each accounting for only 10% of genes) remain recoverable when their signal is heavily diluted by a dominant major program comprising 80% of all genes.
ii. **Null-background regime**: Fixed at *G* = 3000 and *s* = 1, this setting incorporated three active signal programs alongside an explicit non-informative null program ***γ***_null_ = **0**, whose genes carry no cell-type-linked genetic signal and generate GWAS summary statistics purely from LD structure and sampling noise. The background fraction of null genes *f*_null_ was systematically varied across {0.25, 0.50, 0.75, 0.875, 0.9375}, with the remaining (1 − *f*_null_)*G* genes distributed equally among the three active signal programs defined by the generating coefficient matrix **Γ**^(3)^. This regime assesses false-positive control and compares the operating characteristics under increasingly noisy gene backgrounds.
iii. **Collinearity regime**: Designed to test false-positive control under correlated cell-type expression, expression profiles were modified such that a non-causal cell type shared a strong linear correlation (*r* > 0.8) with a true causal cell type. This regime evaluates whether THREAD’s sparsity-inducing prior effectively suppresses false discoveries on correlated bystander cell types while maintaining high sensitivity for true causal drivers.

### Benchmarking against existing methods

On identical inputs, THREAD was compared with established cell-type prioritisation methods: MAGMA gene-property analysis and RolyPoly in the simulations, and MAGMA, RolyPoly and scPagwas in the psychiatric application. All comparators were run with default parameters. For the cross-method comparison, each method’s per-cell-subtype significance was expressed as −log_10_ *P* and ranked within method; agreement between methods, and between disorders, was quantified by the Spearman rank correlation of these scores (for example, schizophrenia versus bipolar disorder for THREAD, and RolyPoly versus THREAD within schizophrenia). On real data, where the data-generating coefficients are unknown, no oracle reference was available and methods were compared on the cell subtypes they prioritised.

## Discussion

We set out to ask not merely which cell subtypes are enriched for the heritability of a complex disease, but how that heritability is organized across them. THREAD answers this with a representation that sits deliberately between two prevailing views of disease genetics: a single coefficient vector linking cell-type expression to genetic risk, shared across all genes, and methods that assign that risk at single-cell resolution.

By placing a Dirichlet process mixture over genes, THREAD partitions trait-associated genes into a data-driven number of latent modules, each governed by its own vector of cell-subtype coefficients, and interprets the cell subtypes jointly prioritized within a module as a coordinated, multicellular program. Across neurodegeneration, psychiatric disease and two cancers, this decomposition recovered cellular programs that were biologically coherent, reproducible in independent cohorts, and aligned with clinically meaningful axes of disease.

This resolution follows from two design choices. Because the mixture is nonparametric, THREAD learns the number of programs and their gene membership jointly with the coefficients, separating genes driven by different cell subtypes rather than averaging them into one enrichment score; our simulations show that a single-coefficient model does not merely lose power on heterogeneous risk but misattributes it, propagating each detected cell subtype across programs in which it plays no part. THREAD thus generalizes the single-component polygenic regression of which RolyPoly is a special case, while avoiding the dropout sensitivity incurred by scoring risk at single-cell resolution. The likelihood, moreover, is heteroscedastic by derivation—the sampling variance of each gene’s estimate is a quadratic function of its mean with coefficients fixed by local linkage disequilibrium— so that genes are weighted by their own precision and effect sizes are recovered without bias, and a dual criterion intersecting Bayesian directional certainty with a frequentist false discovery rate converts the inferred coefficients into high-confidence associations.

Applied across diseases of increasing cellular complexity, THREAD produced decompositions specific to each disorder rather than generic. In Alzheimer’s disease, risk resolved into a glia-dominated architecture spanning a neuron–oligodendrocyte program, an astrocytic homeostatic program, and—separately—a compact, cell-type-agnostic core carrying the *APOE*-linked lipid-transport biology^15^, a cross-lineage signal that filtering heritability through cell-type expression alone would miss. In schizophrenia and bipolar disorder, it separated a shared excitatory–inhibitory neuronal core from a schizophrenia-specific interneuron extension as distinct programs rather than one aggregate signal, with druggable targets distributed across them. In the two cancers, the same machinery recovered the organizing axis of solid tumours—a tumour-intrinsic epithelial program set against a microenvironmental immune–stromal program in colorectal cancer, and two multicellular ecosystem programs anchored on distinct malignant compartments in breast cancer— each tracking an independent clinical axis, from stage progression to molecular subtype and survival. The recurring feature is that the prioritized unit was multicellular: a coordinated set of cell subtypes and pathways rather than a single enriched population. This reframes cell-type mapping from “which cell subtype” to “which coordinated cellular program”, a description better matched to how complex disease biology is now understood.

Several limitations bound these conclusions. THREAD views genetic risk through a single-cell expression reference, so the resolution at which lineages can be prioritized is bounded by the granularity and composition of that atlas; in the entorhinal-cortex reference used for Alzheimer’s disease, microglia constitute only a small fraction of nuclei, limiting the power to resolve a microglial program despite its established relevance, and a different reference could shift the inferred decomposition. The within-module gene score is expression-weighted and is not a direct index of disease relevance—broadly expressed genes rank highly—so we treat it only as a permissive, set-level candidate list and draw conclusions from set-based analyses rather than from individual genes. The framework operates on GWAS summary statistics through an LD-corrected response and a derived variance model, inheriting the assumptions of that reduction, and the cohorts analysed are predominantly of European ancestry, leaving cross-ancestry generalizability to be established. Finally, the inferred programs are associational decompositions of inherited risk: they nominate coordinated cellular mechanisms and align them with independent phenotypes, but establishing causality will require direct experimental perturbation.

These limitations point to natural extensions. Matched, higher-resolution single-cell and spatial references—and multi-ancestry GWAS—would sharpen and broaden the decompositions, while integrating regulatory layers such as eQTL and chromatin-accessibility data could move the inferred programs from correlation toward mechanism; modelling disease-stage or temporal structure, and validating prioritized programs by perturbation, are clear next steps. Because THREAD requires only GWAS summary statistics and a single-cell reference, it is readily applicable across the growing catalogue of well-powered GWAS and cell atlases. By resolving polygenic risk into interpretable, multicellular programs, THREAD offers a general and reproducible route from a macroscopic genetic association to the coordinated cellular biology through which it acts.

## Data availability

All datasets analysed in this study are publicly available from the indicated repositories. Alzheimer’s disease GWAS summary statistics were obtained from IEU OpenGWAS (dataset <monospace>ieu-b-2</monospace>; https://opengwas.io/datasets/ieu-b-2). Schizophrenia GWAS summary statistics were obtained from the Psychiatric Genomics Consortium 2014 study (PGC SCZ 2014; https://figshare.com/articles/dataset/scz2014/14672163?file=28570554), and bipolar disorder GWAS summary statistics were obtained from the PGC3 study (https://figshare.com/articles/dataset/PGC3_bipolar_disorder_GWAS_summary_statistics/14102594). Colorectal-cancer GWAS summary statistics were obtained from FinnGen through IEU OpenGWAS (dataset finn-b-C3 COLORECTAL EXALLC; https://opengwas.io/datasets/finn-b-C3_COLORECTAL_EXALLC). Breast-cancer GWAS summary statistics were obtained from the Breast Cancer Association Consortium (BCAC; https://www.ccge.medschl.cam.ac.uk/breast-cancer-association-consortium-bcac/data-data-access/summary-results/gwas-summary-results).

The transcriptomic datasets used in the disease-specific analyses are available from the NCBI Gene Expression Omnibus (GEO). The Alzheimer’s disease analyses used GSE138852 (https://www.ncbi.nlm.nih.gov/geo/query/acc.cgi?acc=GSE138852) and the independent validation dataset GSE109887 (https://www.ncbi.nlm.nih.gov/geo/query/acc.cgi?acc=GSE109887). The schizophrenia and bipolar disorder analyses used GSE254569 (https://www.ncbi.nlm.nih.gov/geo/query/acc.cgi?acc=GSE254569); the colorectal-cancer analysis used GSE132465 (https://www.ncbi.nlm.nih.gov/geo/query/acc.cgi?acc=GSE132465); and the breast-cancer analysis used GSE176078 (https://www.ncbi.nlm.nih.gov/geo/query/acc.cgi?acc=GSE176078).

Bulk transcriptomic and clinical data from TCGA-COAD and TCGA-BRCA were accessed through the UCSC Xena public data resource (https://xenabrowser.net/datapages/). The corresponding TCGA projects are also available through the NCI Genomic Data Commons under project identifiers TCGA-COAD (https://portal.gdc.cancer.gov/projects/TCGA-COAD) and TCGA-BRCA (https://portal.gdc.cancer.gov/projects/TCGA-BRCA), respectively.

The 1000 Genomes Project Phase 3 European-ancestry reference panel on GRCh37/hg19, used to derive linkage-disequilibrium information, was obtained as PLINK-format reference files from Zenodo (record 7768714; https://zenodo.org/records/7768714; DOI: https://doi.org/10.5281/zenodo.7768714). Source data underlying the main and Supplementary figures are provided with this paper.

## code availability

The source code for THREAD, an R package implementing the proposed Bayesian nonparametric framework, is publicly available at https://github.com/Jiacheng-Lou/THREAD. The repository provides installation instructions, package documentation, and usage examples.

## Acknowledgments

We thank all study participants, researchers, and consortia who contributed to the public genome-wide association study (GWAS) summary statistics, single-cell transcriptomic atlases, and clinical cohorts analysed in this work. In particular, we acknowledge the Psychiatric Genomics Consortium (PGC), the International Genomics of Alzheimer’s

Project (IGAP), and other respective GWAS consortia for making summary-level genetic data publicly available. We also thank the investigators who generated and shared the single-cell and bulk transcriptomic datasets deposited in the Gene Expression Omnibus and the Single Cell Portal. The results shown here are in part based upon data generated by The Cancer Genome Atlas (TCGA) Research Network (https://www.cancer.gov/tcga). We also thank the members of our research groups for valuable discussions and suggestions.

