## Supplementary Notes, Tables and Figures for "THREAD: a Bayesian nonparametric framework for resolving polygenic risk into multicellular programs"

The Supplementary Notes provide the methodological derivations, simulation protocols, and external validation workflows supporting the main text. Section 1 presents the complete mathematical specification for the framework of THREAD, derives the linkage disequilibrium (LD)-determined variance coefficients that define its heteroscedastic working likelihood, and details the hybrid Markov chain Monte Carlo (MCMC) scheme used for posterior inference and module discovery. Section 2 describes the ground-truth module alignment criteria and evaluation metrics used in the simulation benchmarks. Section 3 details the data processing and statistical procedures for independent transcriptomic and clinical validations in external Alzheimer's disease (GSE109887) and colorectal cancer (TCGA-COAD) cohorts.

### 1 Methodological details of THREAD

#### 1.1 Exact Derivation of Heteroscedastic Sampling Variance

For gene $j$ ($j = 1, \dots, G$), let $S_j$ denote the set of $m_j = |S_j|$ SNPs mapping to the gene locus, and let $R_j \in \mathbf{R}^{m_j \times m_j}$ denote the corresponding local linkage disequilibrium (LD) correlation matrix. Following the polygenic model of the main text, the vector of marginal GWAS effect estimates satisfies $\hat{\boldsymbol{\beta}}_{S_j} = \mathbf{R}_j \boldsymbol{\beta}_{S_j} + n^{-1}\mathbf{X}^\top \boldsymbol{\epsilon}$ with $\boldsymbol{\varepsilon} \sim \mathcal{N}(\mathbf{0}, \sigma_e^2 \mathbf{I}_n)$, where $n$ is the GWAS sample size and $\sigma_e^2$ is the phenotypic residual variance. Assigning the random-effect prior $\boldsymbol{\beta}_{S_j} \mid \theta_j \sim \mathcal{N}(0, \theta_j I_{m_j})$, and marginalizing over $\boldsymbol{\beta}_{S_j}$ yields the distribution of the estimated effect vector $\hat{\boldsymbol{\beta}}_{S_j}$:

$$\hat{\boldsymbol{\beta}}_{S_j} \mid \theta_j \sim \mathcal{N}(0, \boldsymbol{\Sigma}_j), \quad \boldsymbol{\Sigma}_j = \theta_j \mathbf{R}_j^2 + \frac{\sigma_e^2}{n} \mathbf{R}_j, \tag{1}$$

where $\theta_j \geq 0$ is the latent per-SNP genetic variance.

To extract the latent genetic variance $\theta_j$ that connects GWAS summary statistics to the cell-type expression profile $\mathbf{s}_j \in \mathbb{R}^P$, we consider the quadratic summary statistic $\hat{\boldsymbol{\beta}}_{S_j}^\top \hat{\boldsymbol{\beta}}_{S_j}$. By the properties of quadratic forms of centered Gaussian vectors, its expectation satisfies:

$$\mathbb{E}\left[\hat{\boldsymbol{\beta}}_{S_j}^\top \hat{\boldsymbol{\beta}}_{S_j}\right] = \mathrm{tr}(\boldsymbol{\Sigma}_j) = \theta_j \,\mathrm{tr}(\mathbf{R}_j^2) + \frac{\sigma_e^2}{n} \mathrm{tr}(\mathbf{R}_j) = \theta_j \,\mathrm{tr}(\mathbf{R}_j^2) + \frac{m_j \sigma_e^2}{n},$$

utilizing the identity $\mathrm{tr}(\mathbf{R}_j) = m_j$ for standardized genotypes. Subtracting the contribution of sampling noise, $m_j \sigma_e^2 / n$, and normalizing by $\mathrm{tr}(\mathbf{R}_j^2)$ defines the gene-level response variable:

$$\xi_j = \frac{\hat{\boldsymbol{\beta}}_{S_j}^\top \hat{\boldsymbol{\beta}}_{S_j} - m_j \sigma_e^2 n^{-1}}{\mathrm{tr}(\mathbf{R}_j^2)},$$

which is an exact unbiased estimator of $\theta_j$, i.e., $\mathbb{E}[\xi_j] = \theta_j$.

Since the sampling variance of $\hat{\boldsymbol{\beta}}_{S_j}^\top \hat{\boldsymbol{\beta}}_{S_j}$ is $2\,\mathrm{tr}(\boldsymbol{\Sigma}_j^2)$, $\xi_j$ is an affine transformation of $\hat{\boldsymbol{\beta}}_{S_j}^\top \hat{\boldsymbol{\beta}}_{S_j}$, its variance is:

$$\mathrm{Var}(\xi_j) = \frac{2\,\mathrm{tr}(\boldsymbol{\Sigma}_j^2)}{\left[\mathrm{tr}(\mathbf{R}_j^2)\right]^2}. \tag{2}$$

Expanding $\boldsymbol{\Sigma}_j^2$ algebraically as a polynomial in $\theta_j$:

$$\boldsymbol{\Sigma}_j^2 = \left(\theta_j \mathbf{R}_j^2 + \frac{\sigma_e^2}{n} \mathbf{R}_j\right)^2 = \theta_j^2 \mathbf{R}_j^4 + 2\theta_j \frac{\sigma_e^2}{n} \mathbf{R}_j^3 + \frac{\sigma_e^4}{n^2} \mathbf{R}_j^2.$$

Taking the trace and substituting into Eq. (2) yields the exact heteroscedastic sampling variance:

$$v_j \equiv \mathrm{Var}(\xi_j) = d_{j0} + d_{j1}\theta_j + d_{j2}\theta_j^2,$$

where the variance coefficients $d_{j0}$, $d_{j1}$ and $d_{j2}$ are deterministic scalars determined entirely by $n$, $\sigma_e^2$, and the LD matrix traces:

$$d_{j0} = \frac{2\sigma_e^4}{n^2 \operatorname{tr}(\mathbf{R}_j^2)}, \quad d_{j1} = \frac{4\sigma_e^2 \operatorname{tr}(\mathbf{R}_j^3)}{n \left[\operatorname{tr}(\mathbf{R}_j^2)\right]^2}, \quad d_{j2} = \frac{2 \operatorname{tr}(\mathbf{R}_j^4)}{\left[\operatorname{tr}(\mathbf{R}_j^2)\right]^2}.$$

#### 1.2 Hierarchical Model

To elucidate how cell-type-specific transcriptional profiles mediate complex trait polygenic risk while accommodating functional heterogeneity across the genome, we formulate a non-parametric Bayesian regression framework governed by a Dirichlet process mixture (DPM). Standard integrative methods typically assume a uniform, genome-wide mapping from cellular expression to genetic architecture, which overlooks the fact that distinct biological pathways often operate through distinct cell types.

To overcome this limitation, THREAD clusters genes $j = 1, \dots, G$ into latent functional modules. Within module $k$, genes share a common coefficient vector $\boldsymbol{\gamma}_k$, which describes how cell-type expression contributes to genetic risk, whereas the coefficient vectors are allowed to differ across modules. This construction accommodates functional heterogeneity across genes without requiring the number of modules to be specified in advance.

For gene $j$, let $\boldsymbol{s}_j = (s_{j1}, \dots, s_{jP})^\top \in \mathbb{R}^P$ denote its cell-type expression-specificity profile and let $c_j$ denote its latent module label. The expression profile is linked to the gene-level per-SNP genetic variance through

$$\theta_j = \boldsymbol{s}_j^\top \boldsymbol{\gamma}_k \mid c_j = k.$$

Thus, $\theta_j$ provides the bridge between single-cell expression and GWAS association signals.

Because the true genetic effect-size variance $\theta_j$ of gene $j$ is not directly observed, we construct an LD-corrected gene-level response $\xi_j = (\widehat{\boldsymbol{\beta}}_{S_j}^\top \widehat{\boldsymbol{\beta}}_{S_j} - m_j \sigma_e^2 n^{-1}) / \operatorname{tr}(\boldsymbol{R}_j^2)$ from the GWAS summary statistics. As shown in 1.1, so that $\xi_j$ serves as an unbiased gene-level proxy for the latent genetic variance. Its moment-matched Gaussian working likelihood forms the observation model of THREAD.

**Observation layer.** Based on the derivation in section 1.1, we can specify a Gaussian working likelihood for the LD-corrected gene-level response $\xi_j$:

$$\xi_j \mid \theta_j \sim \mathcal{N}(\theta_j, v_j), \qquad v_j = d_{j0} + d_{j1}\theta_j + d_{j2}\theta_j^2.$$

Conditional on module assignment $c_j = k$, we link $\theta_j$ to its cell-type expression profile $\boldsymbol{s}_j$ through $\theta_j = \boldsymbol{s}_j^\top \boldsymbol{\gamma}_k$. Substituting this relationship into the working likelihood gives the observation model

$$\xi_j \mid \boldsymbol{s}_j, \boldsymbol{\gamma}_k, c_j = k \sim \mathcal{N}\left(\boldsymbol{s}_j^\top \boldsymbol{\gamma}_k, v_{jk}\right),$$
$$v_{jk} = d_{j0} + d_{j1}\left(\boldsymbol{s}_j^\top \boldsymbol{\gamma}_k\right) + d_{j2}\left(\boldsymbol{s}_j^\top \boldsymbol{\gamma}_k\right)^2.$$

**Dirichlet process mixing layer.** To accommodate an unknown and flexible number of latent modules without rigid model selection, we place a Dirichlet process (DP) prior on the module-specific parameters. Let $\boldsymbol{\phi}_k^* = (\boldsymbol{\gamma}_k, \sigma_k^2, \boldsymbol{\tau}_k^2, \lambda_k^2)$ collect the module-specific cell-type coefficient vector $\boldsymbol{\gamma}_k$ along with its associated adaptive shrinkage scale parameters, where $\boldsymbol{\tau}_k^2 = (\tau_{k1}^2, \dots, \tau_{kP}^2)^\top$. For each gene $j \in \{1, \dots, G\}$, let $\boldsymbol{\phi}_j$ denote its gene-specific parameter vector, which takes the value $\boldsymbol{\phi}_{c_j}^*$ under the latent module assignment $c_j$. We assign the Dirichlet process prior:

$$\mathcal{G} \sim \mathrm{DP}(\alpha, \mathcal{H}_0),$$
$$\boldsymbol{\phi}_j \mid \mathcal{G} \overset{\text{i.i.d.}}{\sim} \mathcal{G}, \qquad j = 1, \dots, G,$$

where $\alpha > 0$ is the concentration parameter and $\mathcal{H}_0$ is the base probability measure.

Equivalently, under the stick-breaking representation,

$$\mathcal{G} = \sum_{k=1}^{\infty} \pi_k \delta_{\boldsymbol{\phi}_k^*}, \qquad \boldsymbol{\pi} \sim \mathrm{GEM}(\alpha), \qquad \boldsymbol{\phi}_k^* \overset{\text{i.i.d.}}{\sim} \mathcal{H}_0.$$

Introducing the latent module indicators gives

$$c_j \mid \boldsymbol{\pi} \sim \mathrm{Categorical}(\boldsymbol{\pi}), \qquad \boldsymbol{\phi}_j = \boldsymbol{\phi}_{c_j}^*.$$

Analytically integrating out the infinite mixing proportions $\boldsymbol{\pi}$ yields the Chinese Restaurant Process (CRP) predictive allocation probabilities for the module assignments $\mathbf{c} = (c_1, \ldots, c_G)^\top$:

$$\Pr(c_j = k \mid \mathbf{c}_{-j}, \alpha) = \begin{cases} \dfrac{n_{-j,k}}{G - 1 + \alpha}, & k \text{ is an occupied module,} \\ \dfrac{\alpha}{G - 1 + \alpha}, & k \text{ is a newly generated module,} \end{cases}$$

where $\mathbf{c}_{-j}$ denotes the module assignments of all genes excluding gene $j$, and $n_{-j,k}$ is the number of genes currently allocated to module $k$. Consequently, the number of active, occupied modules $K$ is inferred dynamically from the data.

**Base measure and within-module shrinkage.** The base measure $\mathcal{H}_0$ assigns each module a hierarchical Laplace prior in its normal–exponential scale-mixture representation:

$$\boldsymbol{\gamma}_k \mid \sigma_k^2, \boldsymbol{\tau}_k^2 \sim \mathcal{N}_P\left(\mathbf{0}, \sigma_k^2 \mathbf{D}_k\right), \qquad \mathbf{D}_k = \mathrm{diag}(\tau_{k1}^2, \ldots, \tau_{kP}^2), \tag{3}$$

$$\sigma_k^2 \sim \text{Inverse-Gamma}(\alpha_0, \beta_0), \tag{4}$$

$$\tau_{kp}^2 \mid \lambda_k^2 \sim \mathrm{Exponential}(\lambda_k^2/2), \qquad p = 1, \ldots, P, \tag{5}$$

$$\lambda_k^2 \sim \mathrm{Gamma}(r, \delta). \tag{6}$$

Here, $\sigma_k^2$ controls the overall scale of the coefficients within module $k$, the local scales $\tau_{kp}^2$ provide coefficient-specific adaptation, and $\lambda_k^2$ controls the module-level degree of shrinkage. Marginalizing over $\tau_{kp}^2$ gives

$$\gamma_{kp} \mid \sigma_k^2, \lambda_k \sim \mathrm{Laplace}\left(0, \frac{\sigma_k}{\lambda_k}\right).$$

This prior shrinks coefficients of uninformative cell types toward zero while allowing cell types carrying substantial signals to retain nonzero effects. The prior specification and empirical-Bayes calibration of $(\alpha, \alpha_0, \beta_0, r, \delta)$ are summarized in Supplementary Table 1.

**Gaussian working likelihood.** The statistic $\hat{\boldsymbol{\beta}}_{S_j}^\top \hat{\boldsymbol{\beta}}_{S_j}$ is a sum of $m_j$ weakly dependent squared terms; for $m_j$ not too small we approximate the distribution of $\xi_j$ by a Gaussian matched to its exact first two moments, $\xi_j \mid \theta_j \dot{\sim} \mathcal{N}(\theta_j, v_j)$. Substituting the module-specific linear predictor $\theta_j = \mathbf{s}_j^\top \gamma_k$ for $c_j = k$ gives the heteroscedastic likelihood (**??**)–(**??**), in which the variance $v_{jk}$ depends on module $k$ through $\gamma_k$ and on gene $j$ through $(d_{j0}, d_{j1}, d_{j2})$.

#### 1.3 Joint posterior

Let $\boldsymbol{\xi} = (\xi_1, \ldots, \xi_G)^\top$, $\boldsymbol{S} = (\boldsymbol{s}_1, \ldots, \boldsymbol{s}_G)^\top \in \mathbb{R}^{G \times P}$, and let $\mathcal{C}_k = \{j : c_j = k\}$ and $n_k = |\mathcal{C}_k|$ $(k = 1, \ldots, K)$ denote the set and number of genes currently assigned to occupied module $k$, where $K$ is the current number of occupied modules. For module $k$, as defined before, let $\boldsymbol{\phi}_k^* = \left(\boldsymbol{\gamma}_k, \sigma_k^2, \boldsymbol{\tau}_k^2, \lambda_k^2\right)$ denote the module-specific parameter tuple.

Conditional on $c_j = k$, define $\mu_{jk} = \boldsymbol{s}_j^\top \boldsymbol{\gamma}_k$, $v_{jk} = d_{j0} + d_{j1}\mu_{jk} + d_{j2}\mu_{jk}^2$, the contribution of gene $j$ to the likelihood under module $k$ is therefore $\mathcal{N}\left(\xi_j \mid \mu_{jk}, v_{jk}\right)$.

After integrating out the random mixing measure $\mathcal{G}$, the Dirichlet process induces the Chinese restaurant process partition probability

$$p(\boldsymbol{c} \mid \alpha) = \frac{\alpha^K \Gamma(\alpha)}{\Gamma(\alpha + G)} \prod_{k=1}^{K} \Gamma(n_k),$$

where $\boldsymbol{c} = (c_1, \ldots, c_G)^\top$.

Let $h_0(\boldsymbol{\phi}_k^*)$ denote the density associated with the base probability measure $\mathcal{H}_0$. Under the hierarchical normal–exponential scale-mixture specification,

$$h_0(\boldsymbol{\phi}_k^*) = p\left(\boldsymbol{\gamma}_k \mid \sigma_k^2, \boldsymbol{\tau}_k^2\right) p(\sigma_k^2) \left\{ \prod_{p=1}^{P} p(\tau_{kp}^2 \mid \lambda_k^2) \right\} p(\lambda_k^2),$$

where

$$\begin{aligned}
p(\boldsymbol{\gamma}_k \mid \sigma_k^2, \boldsymbol{\tau}_k^2) &= \frac{1}{(2\pi)^{P/2} (\sigma_k^2)^{P/2} |\boldsymbol{D}_k|^{1/2}} \exp\left\{ -\frac{1}{2\sigma_k^2} \boldsymbol{\gamma}_k^\top \boldsymbol{D}_k^{-1} \boldsymbol{\gamma}_k \right\}, \\
p(\sigma_k^2) &= \frac{\beta_0^{\alpha_0}}{\Gamma(\alpha_0)} (\sigma_k^2)^{-(\alpha_0+1)} \exp\left( -\frac{\beta_0}{\sigma_k^2} \right), \\
p(\tau_{kp}^2 \mid \lambda_k^2) &= \frac{\lambda_k^2}{2} \exp\left( -\frac{\lambda_k^2 \tau_{kp}^2}{2} \right), \quad p = 1, \ldots, P, \\
p(\lambda_k^2) &= \frac{\delta^r}{\Gamma(r)} (\lambda_k^2)^{r-1} \exp(-\delta \lambda_k^2).
\end{aligned}$$

Conditioning on the fixed concentration parameter $\alpha$, the fixed hyperparameters $(\alpha_0, \beta_0, r, \delta)$, and the precomputed LD-derived coefficients $\boldsymbol{d} = \{\boldsymbol{d}_j = (d_{j0}, d_{j1}, d_{j2})^\top : j = 1, \ldots, G\}$, the joint posterior density of the partition and the parameters of the occupied modules is

$$p\left(\boldsymbol{c}, \{\boldsymbol{\phi}_k^*\}_{k=1}^K \mid \boldsymbol{\xi}, \boldsymbol{S}, \boldsymbol{d}, \alpha, \alpha_0, \beta_0, r, \delta\right) \propto p(\boldsymbol{c} \mid \alpha) \prod_{k=1}^{K} \left[ h_0(\boldsymbol{\phi}_k^*) \prod_{j \in \mathcal{C}_k} \mathcal{N}\left(\xi_j \mid \boldsymbol{s}_j^\top \boldsymbol{\gamma}_k, v_{jk}\right) \right].$$

Equivalently, expanding the base density $h_0(\boldsymbol{\phi}_k^*)$ gives

$$\begin{aligned}
p\left(\boldsymbol{c}, \{\boldsymbol{\gamma}_k, \sigma_k^2, \boldsymbol{\tau}_k^2, \lambda_k^2\}_{k=1}^K \mid \boldsymbol{\xi}, \boldsymbol{S}, \boldsymbol{d}\right) &\propto p(\boldsymbol{c} \mid \alpha) \prod_{k=1}^{K} \left[ \prod_{j \in \mathcal{C}_k} \mathcal{N}\left(\xi_j \mid \boldsymbol{s}_j^\top \boldsymbol{\gamma}_k, v_{jk}\right) \right] \\
\times \prod_{k=1}^{K} \left[ \mathcal{N}_P\left(\boldsymbol{\gamma}_k \mid \boldsymbol{0}, \sigma_k^2 \boldsymbol{D}_k\right) p(\sigma_k^2 \mid \alpha_0, \beta_0) p(\lambda_k^2 \mid r, \delta) \times \prod_{p=1}^{P} p\left(\tau_{kp}^2 \mid \lambda_k^2\right) \right]. \quad (7)
\end{aligned}$$

Posterior computation alternates between updates of the allocation variables and the parameters of the occupied modules. The module allocations are updated using Neal's auxiliary-variable Algorithm 8, the non-conjugate coefficient vectors are updated numerically, and the remaining shrinkage parameters are updated by exact Gibbs sampling.

Posterior computation proceeds by cycling through the full conditional distributions derived below; conjugate parameters are updated by exact Gibbs draws and the non-conjugate coefficient vectors by slice sampling.

##### 1.4 Full conditional distributions and updates

Conditioning on the observed gene-level responses $\boldsymbol{\xi} = (\xi_1, \ldots, \xi_G)^\top$, the cell-type expression-specificity matrix $\boldsymbol{S} = (\boldsymbol{s}_1, \ldots, \boldsymbol{s}_G)^\top \in \mathbb{R}^{G \times P}$, the precomputed LD-derived variance coefficients $\boldsymbol{d} = \{\boldsymbol{d}_j = (d_{j0}, d_{j1}, d_{j2})^\top : j = 1, \ldots, G\}$, the concentration parameter $\alpha$, and the hyperparameters $(\alpha_0, \beta_0, r, \delta)$, posterior inference proceeds via a hybrid Markov chain Monte Carlo (MCMC) sampler. To initialize the chain, initial module assignments $\boldsymbol{c}^{(0)}$ are obtained by $k$-means clustering of trait-associated genes in a pre-trained Gene2vec co-expression embedding [Du et al., 2019], with genes lacking an embedding assigned via $k$-nearest-neighbor voting on their scaled expression profiles; this initialization affects only the starting state $\boldsymbol{c}^{(0)}$ and not the stationary posterior distribution. In each iteration, the sampler cycles through the full conditional distributions of the latent module allocations $\boldsymbol{c} = (c_1, \ldots, c_G)^\top$ and the module-specific parameter tuples $\{\boldsymbol{\phi}_k^*\}_{k=1}^K$, where $\boldsymbol{\phi}_k^* = (\boldsymbol{\gamma}_k, \sigma_k^2, \boldsymbol{\tau}_k^2, \lambda_k^2)$. The MCMC sampling procedure is executed for $T$ iterations, discarding the first $B$ iterations as burn-in and thinning the remaining draws at interval $L$ to form the retained posterior ensemble $\mathcal{T}$.

###### 1.4.1 Update of latent module assignments $c_j$

We update the latent module assignments $c_j$ ($j = 1, \ldots, G$) using the auxiliary-variable Gibbs sampler [Neal, 2000]. Specifically, when updating gene $j$, let $\boldsymbol{c}_{-j}$ denote the module assignments of all genes excluding $j$, let $K^-$ denote the number of distinct occupied modules in $\boldsymbol{c}_{-j}$, and let $n_{-j,k} = |\{j' \neq j : c_{j'} = k\}|$ ($k = 1, \ldots, K^-$) denote the number of genes currently assigned to occupied module $k$ after removing gene $j$. If gene $j$ was not a singleton prior to removal, we introduce $m$ temporary auxiliary components indexed by $k = K^- + 1, \ldots, K^- + m$, whose parameter tuples $\boldsymbol{\phi}_k^* = (\boldsymbol{\gamma}_k, \sigma_k^2, \boldsymbol{\tau}_k^2, \lambda_k^2)$ are drawn independently from the base density $h_0(\boldsymbol{\phi}_k^*)$ by sequentially sampling:

$$\lambda_k^2 \sim \text{Gamma}(r, \delta), \tag{8}$$

$$\tau_{kp}^2 \mid \lambda_k^2 \overset{\text{i.i.d.}}{\sim} \text{Exponential}\left(\frac{\lambda_k^2}{2}\right), \qquad p = 1, \ldots, P, \tag{9}$$

$$\sigma_k^2 \sim \text{Inverse-Gamma}(\alpha_0, \beta_0), \tag{10}$$

$$\boldsymbol{\gamma}_k \mid \sigma_k^2, \boldsymbol{\tau}_k^2 \sim \mathcal{N}_P(\boldsymbol{0}, \sigma_k^2 \boldsymbol{D}_k), \qquad \boldsymbol{D}_k = \text{diag}(\tau_{k1}^2, \ldots, \tau_{kP}^2). \tag{11}$$

If gene $j$ was a singleton in $\boldsymbol{c}$, its existing parameter tuple is retained as one of the $m$ auxiliary components, and the remaining $m - 1$ auxiliary tuples are drawn i.i.d. from $h_0(\boldsymbol{\phi}_k^*)$. For each candidate module $k \in \{1, \ldots, K^- + m\}$, the linear predictor $\mu_{jk} = \boldsymbol{s}_j^\top \boldsymbol{\gamma}_k$ and the variance $v_{jk} = d_{j0} + d_{j1}\mu_{jk} + d_{j2}\mu_{jk}^2$ are evaluated deterministically.

The full conditional allocation probability for gene $j$ over the $K^- + m$ candidate modules is given by

$$\Pr\Big(c_j = k \,\Big|\, \boldsymbol{c}_{-j}, \xi_j, \boldsymbol{s}_j, \boldsymbol{d}_j, \{\boldsymbol{\phi}_\ell^*\}_{\ell=1}^{K^-+m}, \alpha\Big)$$
$$\propto \begin{cases} \dfrac{n_{-j,k}}{G - 1 + \alpha} \mathcal{N}\Big(\xi_j \mid \boldsymbol{s}_j^\top \boldsymbol{\gamma}_k, v_{jk}\Big), & k = 1, \ldots, K^-, \\[2ex] \dfrac{\alpha/m}{G - 1 + \alpha} \mathcal{N}\Big(\xi_j \mid \boldsymbol{s}_j^\top \boldsymbol{\gamma}_k, v_{jk}\Big), & k = K^- + 1, \ldots, K^- + m. \end{cases}$$

If an auxiliary component $k > K^-$ is selected, it is instantiated as a newly occupied module retaining its proposed tuple $\boldsymbol{\phi}_k^*$; all unselected auxiliary components are discarded, empty modules are removed, and the active occupied modules are relabeled as $1, \ldots, K$.

###### 1.4.2 Update of regression coefficients $\boldsymbol{\gamma}_k$

**Full conditional posterior.** For each occupied module $k \in \{1, \ldots, K\}$, let $\mathcal{C}_k = \{j : c_j = k\}$ denote the index set of genes assigned to module $k$, with $n_k = |\mathcal{C}_k|$. Isolating the terms in the joint posterior density that depend on $\boldsymbol{\gamma}_k$, the full conditional density satisfies

$$p\Big(\boldsymbol{\gamma}_k \;\Big|\; \{\xi_j, \boldsymbol{s}_j\}_{j\in\mathcal{C}_k}, \sigma_k^2, \boldsymbol{\tau}_k^2\Big) \propto p\Big(\boldsymbol{\gamma}_k \mid \sigma_k^2, \boldsymbol{\tau}_k^2\Big) \prod_{j\in\mathcal{C}_k} \mathcal{N}\Big(\xi_j \mid \boldsymbol{s}_j^\top \boldsymbol{\gamma}_k, v_{jk}\Big).$$

Taking the logarithm, we have:

$$\log p\Big(\boldsymbol{\gamma}_k \;\Big|\; \{\xi_j, \boldsymbol{s}_j\}_{j\in\mathcal{C}_k}, \sigma_k^2, \boldsymbol{\tau}_k^2\Big) = \underbrace{-\frac{1}{2}\sum_{j\in\mathcal{C}_k} \log v_{jk}}_{\text{(A)}} \underbrace{-\frac{1}{2}\sum_{j\in\mathcal{C}_k} \frac{(\xi_j - \boldsymbol{s}_j^\top \boldsymbol{\gamma}_k)^2}{v_{jk}}}_{\text{(B)}} \underbrace{-\frac{1}{2\sigma_k^2}\boldsymbol{\gamma}_k^\top \boldsymbol{D}_k^{-1}\boldsymbol{\gamma}_k}_{\text{(C)}} + \text{const}, \tag{12}$$

where the constant denotes additive terms that do not depend on $\boldsymbol{\gamma}_k$, and $v_{jk} = d_{j0} + d_{j1}(\boldsymbol{s}_j^\top \boldsymbol{\gamma}_k) + d_{j2}(\boldsymbol{s}_j^\top \boldsymbol{\gamma}_k)^2$. In a conventional homoscedastic regression model where $v_{jk}$ is independent of $\boldsymbol{\gamma}_k$, term (A) is constant with respect to $\boldsymbol{\gamma}_k$, and the combination of (B) and (C) forms a standard quadratic function in $\boldsymbol{\gamma}_k$, yielding a conjugate multivariate Gaussian distribution. Under our LD-informed heteroscedastic model, $\boldsymbol{\gamma}_k$ enters term (A) through the logarithm of a quadratic polynomial and term (B) through both the numerator and the denominator of a rational function. Consequently, the full conditional density in Eq. (12) does not belong to any standard closed-form distribution family. Therefore, we adopt stepping-out slice sampling to update $\boldsymbol{\gamma}_k$.

**Component-wise stepping-out slice sampling.** To sample efficiently from Eq. (12) without manual step-size tuning or gradient-based proposal rejections, we update the $P$ coordinates of $\boldsymbol{\gamma}_k = (\gamma_{k1}, \ldots, \gamma_{kP})^\top$ sequentially using the univariate stepping-out slice sampler [Neal, 2003]. For each coordinate $p \in \{1, \ldots, P\}$, let $f_p(\gamma_{kp})$ denote the log full conditional density in Eq. (12) viewed as a univariate function of $\gamma_{kp}$, with the remaining components $\boldsymbol{\gamma}_{k,-p}$ held fixed. At iteration $t$, slice sampling updates $\gamma_{kp}^{(t)}$ by first drawing a vertical density level $y \sim \text{Uniform}\big(0, \exp\{f_p(\gamma_{kp}^{(t)})\}\big)$ and then sampling $\gamma_{kp}^{(t+1)}$ uniformly from the resulting horizontal slice above $y$. Given a bracket width $w > 0$, one coordinate update proceeds in three steps:

1. *Define the horizontal slice:* Draw an auxiliary variable $e \sim \text{Exponential}(1)$ and define the vertical log-slice threshold

$$z = f_p\Big(\gamma_{kp}^{(t)}\Big) - e \quad \big(\text{so that } y = e^z \sim \text{Uniform}\big(0, \exp\{f_p(\gamma_{kp}^{(t)})\}\big)\big),$$

which defines the target horizontal slice $S_z = \{\gamma_{kp} \in \mathbb{R} : f_p(\gamma_{kp}) \geq z\}$.

2. *Bracket the slice $S_z$ via stepping-out:* Because the exact boundaries of $S_z$ are usually unavailable in closed form, we construct an enclosing interval $[L, R]$. Draw $u \sim \text{Uniform}(0, 1)$ and place an initial horizontal bracket $[L, R]$ of width $w$ randomly around $\gamma_{kp}^{(t)}$ by setting

$$L = \gamma_{kp}^{(t)} - wu, \qquad R = L + w.$$

Then, repeatedly expand the left endpoint via $L \leftarrow L - w$ while $f_p(L) > z$, and expand the right endpoint via $R \leftarrow R + w$ while $f_p(R) > z$, until both endpoints lie below the slice threshold $z$.

3. *Shrinkage procedure:* Draw a candidate value $\gamma_{kp}^* \sim \text{Uniform}(L, R)$. If $\gamma_{kp}^* \in S_z$ (i.e., $f_p(\gamma_{kp}^*) \geq z$), accept the candidate by setting $\gamma_{kp}^{(t+1)} = \gamma_{kp}^*$ and terminate the loop. Otherwise, shrink the bracket toward the current state $\gamma_{kp}^{(t)}$ by setting $L \leftarrow \gamma_{kp}^*$ if $\gamma_{kp}^* < \gamma_{kp}^{(t)}$ or $R \leftarrow \gamma_{kp}^*$ if $\gamma_{kp}^* > \gamma_{kp}^{(t)}$, and draw a new candidate uniformly from the reduced interval $[L, R]$ until a candidate inside $S_z$ is drawn.

##### 1.4.3 Update of the global prior scale $\sigma_k^2$

In the model parameterization of THREAD, the full conditional of $\sigma_k^2$ depends only on the inverse-gamma prior and the normal prior for $\boldsymbol{\gamma}_k$, therefore we have:

$$p(\sigma_k^2 \mid \boldsymbol{\gamma}_k, \boldsymbol{\tau}_k^2) \propto (\sigma_k^2)^{-P/2} \exp\left[-\frac{\boldsymbol{\gamma}_k^\top \boldsymbol{D}_k^{-1} \boldsymbol{\gamma}_k}{2\sigma_k^2}\right] (\sigma_k^2)^{-\alpha_0-1} \exp\left(-\frac{\beta_0}{\sigma_k^2}\right). \tag{13}$$

Consequently,

$$\sigma_k^2 \mid \boldsymbol{\gamma}_k, \boldsymbol{\tau}_k^2 \sim \text{Inverse-Gamma}\left(\alpha_0 + \frac{P}{2}, \beta_0 + \frac{1}{2}\boldsymbol{\gamma}_k^\top \boldsymbol{D}_k^{-1} \boldsymbol{\gamma}_k\right), \tag{14}$$

which permits an exact Gibbs update.

##### 1.4.4 Update of $\tau_{kp}^2$ and $\lambda_k^2$

The shrinkage hyperparameters $\tau_{kp}^2$ interact with the data only through $\boldsymbol{\gamma}_k$, so its full conditional distributions follow directly from the standard Bayesian Lasso [Park and Casella, 2008]. Combining the normal prior in (3) with the exponential prior in (5), the full conditional density for each local scale $\tau_{kp}^2$ $(p = 1, \ldots, P)$ is

$$p\left(\tau_{kp}^2 \,\middle|\, \gamma_{kp}, \sigma_k^2, \lambda_k^2\right) \propto \left(\tau_{kp}^2\right)^{-1/2} \exp\left\{-\frac{\gamma_{kp}^2}{2\sigma_k^2 \tau_{kp}^2} - \frac{\lambda_k^2}{2}\tau_{kp}^2\right\}.$$

Applying the change of variables $\eta_{kp} = 1/\tau_{kp}^2$ yields the inverse-Gaussian kernel

$$p(\eta_{kp} \mid \gamma_{kp}, \sigma_k^2, \lambda_k^2) \propto \eta_{kp}^{-3/2} \exp\{-\frac{\gamma_{kp}^2}{2\sigma_k^2}\eta_{kp} - \frac{\lambda_k^2}{2\eta_{kp}}\},$$

so that

$$\eta_{kp} = \frac{1}{\tau_{kp}^2} \,\middle|\, \gamma_{kp}, \sigma_k^2, \lambda_k^2 \;\sim\; \text{Inverse-Gaussian}\left(\sqrt{\frac{\lambda_k^2 \sigma_k^2}{\gamma_{kp}^2}},\; \lambda_k^2\right), \qquad p = 1, \ldots, P, \tag{15}$$

For the penalty parameter $\lambda_k^2$, combining the $P$ independent exponential priors in (3) with the conjugate Gamma prior in (6) yields

$$\lambda_k^2 \mid \boldsymbol{\tau}_k^2 \;\sim\; \text{Gamma}\left(P + r,\; \delta + \frac{1}{2}\sum_{p=1}^{P} \tau_{kp}^2\right). \tag{16}$$

Thus, the updates for both $\tau_{kp}^2$ and $\lambda_k^2$ are exact Gibbs steps requiring no tuning.

#### 1.5 Posterior Summarization, Label Switching, and Convergence

**Label-invariant posterior similarity matrix and point partitioning.** Because the joint posterior density in (7) is invariant to permutations of the module indices $k \in \{1, \ldots, K\}$, the numerical labels assigned to modules within individual MCMC sweeps are exchangeable and not directly comparable across iterations. Moreover, the number of occupied modules may vary between draws. Rather than imposing artificial ordering constraints on the parameter space, we base partition inference on label-invariant pairwise co-clustering probabilities. Let $\mathcal{T}$ denote the collection of $|\mathcal{T}|$ retained post-burn-in iterations. From $\mathcal{T}$, we construct the empirical posterior similarity matrix

$\boldsymbol{Q} \in [0,1]^{G\times G}$, whose $(i,j)$ entry estimates the marginal posterior probability that genes $i$ and $j$ belong to the same latent functional module:

$$\boldsymbol{Q}_{ij} = \frac{1}{|\mathcal{T}|}\sum_{t\in\mathcal{T}} \mathbb{I}\Big(c_i^{(t)} = c_j^{(t)}\Big), \qquad i,j \in \{1,\dots,G\}, \tag{17}$$

where $\mathbb{I}(\cdot)$ denotes the indicator function. It is apparent that $\boldsymbol{Q}_{ij}$ is invariant to permutations of the labels within any draw.

To extract a representative point partition $\hat{\boldsymbol{c}} = (\hat{c}_1,\dots,\hat{c}_G)^\top$ from the posterior ensemble, we convert the posterior similarity matrix $\boldsymbol{Q}$ into a pairwise dissimilarity matrix $\boldsymbol{\Delta} = \mathbf{1}_G\mathbf{1}_G^\top - \boldsymbol{Q}$ and apply partitioning around medoids (PAM). When summarizing the posterior distribution of the active module count, because the Dirichlet process naturally spawns transient singleton or micro-clusters during MCMC exploration, occupied modules containing at most 10 genes in a given posterior draw $t \in \mathcal{T}$ are treated as transient micro-clusters. We then define $K_{\text{mode}}$ as the posterior mode of the number of non-micro modules across $\mathcal{T}$. The final number of modules $\hat{K}$ is selected by maximizing the average silhouette width over candidate module counts within a local window around $K_{\text{mode}}$. The corresponding $\hat{K}$-medoids partition of $\boldsymbol{\Delta}$ directly yields the gene-level module assignments $\hat{\boldsymbol{c}} = (\hat{c}_1,\dots,\hat{c}_G)^\top$ and final gene modules $\hat{\mathcal{C}}_k = \{j \in \{1,\dots,G\} : \hat{c}_j = k\}$ for $k = 1,\dots,\hat{K}$. This two-stage procedure decouples gene-module recovery from iteration-specific cluster labeling.

**Module alignment and dual-significance calling for module–subtype associations.** Once the representative partition $\hat{\boldsymbol{c}}$ and the resulting $\hat{K}$ gene modules $\hat{\mathcal{C}}_k = \{j : \hat{c}_j = k\}$ $(k = 1,\dots,\hat{K})$ are obtained from the optimal $\hat{K}$-medoids clustering of $\boldsymbol{\Delta}$, we resolve iteration-specific label switching by aligning the posterior draws in $\mathcal{T}$ to these fixed modules. Specifically, let $\mathcal{C}_m^{(t)} = \{j : c_j^{(t)} = m\}$ denote the gene set of active module $m \in \{1,\dots,K^{(t)}\}$ in posterior draw $t \in \mathcal{T}$, with associated cell-type coefficient vector $\boldsymbol{\gamma}_m^{(t)} = (\gamma_{m1}^{(t)},\dots,\gamma_{mP}^{(t)})^\top$. For each final module $k$ and draw $t$, we identify the best-matching sampled module $m_k^{(t)}$ via Jaccard index:

$$m_k^{(t)} = \underset{m\in\{1,\dots,K^{(t)}\}}{\arg\max} \frac{|\hat{\mathcal{C}}_k \cap \mathcal{C}_m^{(t)}|}{|\hat{\mathcal{C}}_k \cup \mathcal{C}_m^{(t)}|}, \tag{18}$$

and define the aligned posterior coefficient as $\tilde{\gamma}_{kp}^{(t)} = \gamma^{(t)}_{m_k^{(t)},p}$ for each cell type $p \in \{1,\dots,P\}$. The Bayesian evidence for a positive cell-type contribution to genetic risk in module $k$ is then quantified by the local false sign rate (lFSR), computed from the aligned coefficient $\{\tilde{\gamma}_{kp}^{(t)}\}_{t\in\mathcal{T}}$ as

$$\text{lFSR}_{kp} = \widehat{\Pr}(\tilde{\gamma}_{kp} \le 0 \mid \boldsymbol{\xi}, \boldsymbol{S}, \boldsymbol{d}) = \frac{1}{|\mathcal{T}|}\sum_{t\in\mathcal{T}} \mathbb{I}\Big(\tilde{\gamma}_{kp}^{(t)} \le 0\Big). \tag{19}$$

Since the hierarchical Laplace prior intentionally shrinks posterior coefficient estimates toward zero to stabilize high-dimensional module discovery, we complement the Bayesian sign probabilities $\text{lFSR}_{kp}$ with an unpenalized frequentist refit within each discovered module $\hat{\mathcal{C}}_k$ using feasible generalized least squares (FGLS). Specifically, restricting the regression model to genes $j \in \hat{\mathcal{C}}_k$, we estimate the module-specific coefficient vector $\boldsymbol{\gamma}_k = (\gamma_{k1},\dots,\gamma_{kP})^\top$ via iteratively reweighted least squares (IRLS), using observation weights $\hat{v}_{jk}^{-1}$ defined by

$$\hat{v}_{jk} = \max\Big(d_{j0} + d_{j1}\hat{\mu}_{jk} + d_{j2}\hat{\mu}_{jk}^2,\, v_{\min}\Big), \qquad \hat{\mu}_{jk} = \boldsymbol{s}_j^\top \hat{\boldsymbol{\gamma}}_k. \tag{20}$$

Here, $v_{\min} = 10^{-6}$ is set as a variance lower bound to ensure numerical stability. The variance weights and coefficient estimates are updated iteratively until convergence, yielding the unpenalized estimate

$\hat{\gamma}_k^{\text{FGLS}}$ and its heteroscedasticity-consistent standard errors $\text{SE}(\hat{\gamma}_{kp}^{\text{FGLS}})$. We then perform a one-sided Wald test of $H_0 : \gamma_{kp} \leq 0$ against $H_1 : \gamma_{kp} > 0$ using the test statistic $z_{kp} = \hat{\gamma}_{kp}^{\text{FGLS}}/\text{SE}(\hat{\gamma}_{kp}^{\text{FGLS}})$. To control the false discovery rate (FDR) across all $\hat{K} \times P$ module–cell-type pairs, we apply the Benjamini–Hochberg procedure to the $p$-values to obtain FDR-adjusted $q$-values $q_{kp}$. Under our default dual-significance criterion, a module–cell-type pair $(k, p)$ is declared significant if both $\text{lFSR}_{kp} < 0.05$ and $q_{kp} < 0.05$, with the unshrunk FGLS estimate $\hat{\gamma}_{kp}^{\text{FGLS}}$ reported as the final effect size.

**Convergence diagnostics.** Because individual component labels can switch across MCMC iterations, convergence and partition stability were assessed through label-invariant summaries of the sampled partitions (Supplementary Figs. 1 and 2): (i) the MCMC trace of the number of occupied gene modules $K^{(t)}$ over the recorded iterations, together with its empirical posterior distribution across retained draws $\mathcal{T}$ (Supplementary Fig. Xa,b); (ii) the average silhouette width profile across candidate module counts used to verify the optimality of the selected module count $\hat{K}$; and (iii) the empirical posterior co-clustering matrix $\boldsymbol{Q}$, reordered according to the selected point partition $\hat{\boldsymbol{c}}$ to examine the block-diagonal coherence and stability of the inferred module structure . All posterior summaries were computed exclusively from the post-burn-in retained draws $\mathcal{T}$.

The complete hybrid MCMC sampling scheme and post-processing inference pipeline for THREAD are detailed in Algorithm 1.

**Algorithm 1:** Hybrid MCMC sampler and posterior inference for heteroscedastic Dirichlet-process polygenic regression (DPR)

**Input:** LD-corrected gene responses $\boldsymbol{\xi} = (\xi_1, \ldots, \xi_G)^\top$; cell-type expression $\boldsymbol{S} = (\boldsymbol{s}_1, \ldots, \boldsymbol{s}_G)^\top$; precomputed LD variance coefficients $\boldsymbol{d} = \{\boldsymbol{d}_j = (d_{j0}, d_{j1}, d_{j2})^\top\}_{j=1}^G$; variance floor $v_{\min}$; initial labels $\boldsymbol{c}^{(0)}$; DP concentration $\alpha$ and auxiliary count $m$; hyperparameters $(\alpha_0, \beta_0, r, \delta)$; slice bracket width $w$; iterations $T$, burn-in $B$, thinning interval $L$.

**Output:** Retained post-burn-in draws $\mathcal{T}$; representative point partition $\hat{\boldsymbol{c}}$ and modules $\{\hat{\mathcal{C}}_k\}_{k=1}^{\hat{K}}$; dual-significant module–cell-type associations $(k, p)$ with effect sizes $\hat{\gamma}_{kp}^{\text{FGLS}}$.

Initialize $\boldsymbol{c} \leftarrow \boldsymbol{c}^{(0)}$, occupied module count $K$, and parameter tuples $\{\boldsymbol{\phi}_k^* = (\boldsymbol{\gamma}_k, \sigma_k^2, \boldsymbol{\tau}_k^2, \lambda_k^2)\}_{k=1}^K$; set $\mathcal{T} \leftarrow \varnothing$

**for** $t = 1, \ldots, T$ **do**
- **for** $j = 1, \ldots, G$ **do**
  - Let $K^-$ be the number of occupied modules in $\boldsymbol{c}_{-j}$, relabeled as $\{1, \ldots, K^-\}$ with gene counts $n_{-j,k}$
  - **if** *gene* $j$ *was a singleton in* $\boldsymbol{c}$ **then**
    - Retain its tuple $\boldsymbol{\phi}_{K^-+1}^* \leftarrow \boldsymbol{\phi}_{c_j}^*$ and draw $\boldsymbol{\phi}_{K^-+2}^*, \ldots, \boldsymbol{\phi}_{K^-+m}^* \overset{\text{i.i.d.}}{\sim} h_0(\boldsymbol{\phi}^*)$ via Eqs. (8)–(11)
  - **else**
    - Draw $m$ auxiliary tuples $\boldsymbol{\phi}_{K^-+1}^*, \ldots, \boldsymbol{\phi}_{K^-+m}^* \overset{\text{i.i.d.}}{\sim} h_0(\boldsymbol{\phi}^*)$ via Eqs. (8)–(11)
  - **end**
  - **for** $k = 1, \ldots, K^- + m$ **do**
    - $\mu_{jk} \leftarrow \boldsymbol{s}_j^\top \boldsymbol{\gamma}_k; \quad v_{jk} \leftarrow \max\big(d_{j0} + d_{j1}\mu_{jk} + d_{j2}\mu_{jk}^2,\ v_{\min}\big)$
    - $\ell_k \leftarrow \log \mathcal{N}(\xi_j \mid \mu_{jk}, v_{jk}) + \mathbb{I}(k \le K^-) \log n_{-j,k} + \mathbb{I}(k > K^-) \log(\alpha/m)$
  - **end**
  - Sample $c_j \sim \text{Categorical}\{\text{softmax}(\ell_1, \ldots, \ell_{K^-+m})\}$
  - **if** $c_j > K^-$ **then** instantiate a new occupied module retaining the selected auxiliary tuple $\boldsymbol{\phi}_{c_j}^*$
  - Discard unselected auxiliary tuples and relabel active occupied modules as $1, \ldots, K$
- **end**
- **for** $k = 1, \ldots, K$ **do**
  - **for** $p = 1, \ldots, P$ **do**
    - Update $\gamma_{kp} \mid \boldsymbol{\gamma}_{k,-p}, \sigma_k^2, \boldsymbol{\tau}_k^2$ via univariate stepping-out slice sampling (width $w$) on the log full conditional $f_p(\gamma_{kp})$ in Eq. (12)
  - **end**
  - Sample $\sigma_k^2 \mid \boldsymbol{\gamma}_k, \boldsymbol{\tau}_k^2 \sim \text{Inverse-Gamma}\big(\alpha_0 + \frac{P}{2},\ \beta_0 + \frac{1}{2}\boldsymbol{\gamma}_k^\top \boldsymbol{D}_k^{-1} \boldsymbol{\gamma}_k\big)$ via Eq. (14)
  - **for** $p = 1, \ldots, P$ **do**
    - Sample $\eta_{kp} \mid \gamma_{kp}, \sigma_k^2, \lambda_k^2 \sim \text{Inverse-Gaussian}\Big(\sqrt{\lambda_k^2 \sigma_k^2 / \gamma_{kp}^2},\ \lambda_k^2\Big)$ via Eq. (15), and set $\tau_{kp}^2 \leftarrow 1/\eta_{kp}$
  - **end**
  - Sample $\lambda_k^2 \mid \boldsymbol{\tau}_k^2 \sim \text{Gamma}\Big(P + r,\ \delta + \frac{1}{2}\sum_{p=1}^P \tau_{kp}^2\Big)$ via Eq. (16)
- **end**
- **if** $t > B$ ***and*** $(t - B) \bmod L = 0$ **then** append draw $t$ with state $\big(\boldsymbol{c}^{(t)}, \{\boldsymbol{\phi}_m^{*(t)}\}_{m=1}^{K^{(t)}}, K^{(t)}\big)$ to $\mathcal{T}$

**end**

Compute posterior similarity $Q_{ij} \leftarrow |\mathcal{T}|^{-1} \sum_{t \in \mathcal{T}} \mathbb{I}\big(c_i^{(t)} = c_j^{(t)}\big)$ and dissimilarity matrix $\boldsymbol{\Delta} \leftarrow \boldsymbol{1}_G \boldsymbol{1}_G^\top - \boldsymbol{Q}$

Obtain point partition $\hat{\boldsymbol{c}}$ and modules $\hat{\mathcal{C}}_k = \{j : \hat{c}_j = k\}$ $(k = 1, \ldots, \hat{K})$ by PAM on $\boldsymbol{\Delta}$

**for** $k = 1, \ldots, \hat{K}$ **do**
- For each draw $t \in \mathcal{T}$, match $\hat{\mathcal{C}}_k$ to active cluster $m_k^{(t)}$ via Eq. (18) and set aligned draws $\tilde{\gamma}_{kp}^{(t)} \leftarrow \gamma_{m_k^{(t)},p}^{(t)}$; compute $\text{lFSR}_{kp}$ via Eq. (**??**)
- Refit module $\hat{\mathcal{C}}_k$ via FGLS until convergence to obtain $\hat{\boldsymbol{\gamma}}_k^{\text{FGLS}}$ and $\text{SE}(\hat{\gamma}_{kp}^{\text{FGLS}})$

**end**

Compute one-sided Wald statistics $z_{kp} = \hat{\gamma}_{kp}^{\text{FGLS}} / \text{SE}(\hat{\gamma}_{kp}^{\text{FGLS}})$ and FDR-adjusted $q$-values $q_{kp}$

Report $(k, p)$ as dual-significant if $\text{lFSR}_{kp} < 0.05$ and $q_{kp} < 0.05$, with final effect size $\hat{\gamma}_{kp}^{\text{FGLS}}$

#### 2 Simulation Implementation, Benchmark Protocols, and Evaluation Metrics

The data-generating process, true program coefficient matrices $\mathbf{\Gamma}^{(K_0)}$ ($K_0 \in \{1, 3, 5\}$), mixing proportions $\boldsymbol{\pi}$, total gene counts $G \in \{500, 1000, 3000\}$, coefficient scale factors $s \in \{1, 10\}$, and benchmark settings are specified in the Methods section of the main text. Here, we provide the implementation and scoring details used to obtain the simulation results. All simulated datasets were analyzed using the same THREAD inference pipeline (Algorithm 1), running the MCMC sampler for $T = 5,000$ iterations with the first $B = 4,000$ iterations discarded as burn-in and every post-burn-in draw retained without thinning ($L = 1$, yielding $|\mathcal{T}| = 1,000$ posterior draws).

##### 2.1 Ground-Truth Module Alignment and Significance Criteria

Because mixture labels are exchangeable, the $\hat{K}$ inferred THREAD modules $\hat{\mathcal{C}}_{\hat{k}} = \{j : \hat{c}_j = \hat{k}\}$ ($\hat{k} \in \{1, \ldots, \hat{K}\}$) were aligned to the $K_0$ ground-truth generating programs $\mathcal{C}^*_k = \{j : c_j = k\}$ ($k \in \{1, \ldots, K_0\}$) prior to program-level evaluation by maximizing pairwise gene-set overlap (Jaccard index):

$$\hat{k}(k) = \underset{\hat{k} \in \{1, \ldots, \hat{K}\}}{\arg\max} \frac{|\mathcal{C}^*_k \cap \hat{\mathcal{C}}_{\hat{k}}|}{|\mathcal{C}^*_k \cup \hat{\mathcal{C}}_{\hat{k}}|}, \qquad k = 1, \ldots, K_0. \tag{21}$$

Program-level cell-type discovery was then evaluated across all $K_0 \times P$ ground-truth program–subtype pairs $(k, p)$, where a pair $(k, p)$ is defined as truly causal if $\gamma^*_{kp} > 0$ and non-causal if $\gamma^*_{kp} = 0$.

Within each aligned module, THREAD calls were determined by our default **dual-significance** criterion, which requires both a local false sign rate $\mathrm{lFSR}_{kp} < 0.05$ and a BH-adjusted $q_{kp} < 0.05$ from the within-module FGLS refit. To dissect the contribution of each component, we also separately evaluated two single-criterion variants: **Bayesian-only** ($\mathrm{lFSR}_{kp} < 0.05$) and **Frequentist-only** ($q_{kp} < 0.05$). Because MAGMA and RolyPoly return only a single genome-wide decision vector $\boldsymbol{v} = (v_1, \ldots, v_P)^\top \in \{0, 1\}^P$ (at BH-adjusted FDR $< 0.05$) rather than program-specific calls, we replicated $\boldsymbol{v}$ across all $K_0$ programs for evaluation: if cell type $p$ was identified as globally significant ($v_p = 1$), it was treated as a positive call in every program $k \in \{1, \ldots, K_0\}$.

##### 2.2 Evaluation Metrics and Summary Reporting

We defined an idealized **oracle FGLS estimator** for simulation benchmarking by fixing the gene-to-program indicators to their ground-truth labels $\boldsymbol{c} = (c_1, \ldots, c_G)^\top$, rather than inferring them via the Dirichlet process mixture. Given the true gene partition, it directly fits a separate FGLS model within each true program via IRLS, serving as a theoretical upper bound for coefficient recovery.

Model performance was quantified across 50 independent Monte Carlo replicates per setting along three aspects:

1. **Program-level subtype discovery:** For each true program $k \in \{1, \ldots, K_0\}$, $\mathrm{Precision}_k$, $\mathrm{Recall}_k$, and the harmonic mean $\mathrm{F1}_k$ were computed by comparing predicted positive cell types against the true causal cell type set $\{p : \gamma^*_{kp} > 0\}$. The primary endpoint was the macro-averaged F1 score across all true programs:

$$\text{Macro-F1} = \frac{1}{K_0} \sum_{k=1}^{K_0} \mathrm{F1}_k, \qquad \mathrm{F1}_k = \frac{2\,\mathrm{Precision}_k \cdot \mathrm{Recall}_k}{\mathrm{Precision}_k + \mathrm{Recall}_k}. \tag{22}$$

   In addition, overall Sensitivity (Recall), Specificity, and empirical FDR were computed across the pooled set of $K_0 \times P$ program–subtype pairs.

2. **Coefficient recovery:** For each aligned program–cell-type pair $(k, p)$, estimation accuracy was evaluated using the FGLS refit estimate $\hat{\gamma}_{kp}^{\mathrm{FGLS}}$ via signed bias and absolute error:

$$\mathrm{Bias}_{kp} = \hat{\gamma}_{kp}^{\mathrm{FGLS}} - \gamma_{kp}^{*}, \qquad \mathrm{AbsBias}_{kp} = \left|\hat{\gamma}_{kp}^{\mathrm{FGLS}} - \gamma_{kp}^{*}\right|. \tag{23}$$

3. **Partition quality:** Gene-program clustering accuracy was quantified by the Adjusted Rand Index (ARI) between the point partition $\hat{\boldsymbol{c}}$ and the true program assignment vector $\boldsymbol{c}$, alongside the posterior mode $K_{\mathrm{mode}}$ of non-micro modules and the final selected module count $\hat{K}$.

### 3 Independent validation in external cohorts

#### 3.1 Independent validation of AD-associated gene modules in GSE109887

Independent validation of the AD-associated gene modules was performed using the GSE109887 middle temporal gyrus expression dataset. Sample titles and phenotypic annotations were extracted from the corresponding GEO series-matrix file. Diagnostic labels were identified from the sample-characteristic fields by matching terms referring to Alzheimer disease or AD and terms referring to control or normal samples. Expression-matrix columns were reconciled with the GEO sample annotations using the numeric sample identifiers embedded in the expression-column names and sample titles. Samples for which a diagnosis could not be assigned unambiguously were excluded. The final analysis included 46 AD samples and 32 control samples, with control samples specified as the reference group.

Gene symbols were obtained from the first annotation field of the expression table. When multiple gene symbols were separated by commas, the first symbol was retained. Entries with missing, empty, or `Unknown` gene symbols were removed. Expression features mapping to the same gene symbol were collapsed using `limma::avereps`. The resulting expression matrix was transformed as $\log_2(x+1)$ and normalized across samples by quantile normalization using `limma::normalizeBetweenArrays`.

Differential-expression analysis was performed using the `limma` framework. A linear model containing diagnosis as the explanatory variable was fitted to each gene, with control samples used as the reference level. Gene-wise variance estimates were moderated by empirical Bayes shrinkage using `limma::eBayes`. The coefficient corresponding to AD relative to control was extracted, and genes were ranked in decreasing order of the moderated $t$-statistic. Positive ranking statistics therefore indicate higher expression in AD, whereas negative statistics indicate lower expression in AD.

Module signatures were constructed from the THREAD gene rankings obtained in the primary AD analysis. Within each of the six inferred modules, genes were ordered by their THREAD rank, duplicate gene symbols were removed, and the top 200 unique genes were retained as the module signature. Each signature was intersected with the gene universe represented in the GSE109887 limma-ranked list. The number and proportion of signature genes successfully mapped to the external dataset were recorded for each module. Genes absent from the external expression dataset were retained in the signature-overlap report but were not included in the enrichment calculation.

Gene-set enrichment analysis was performed using the `fgsea` package, with the limma moderated $t$-statistics used as the ranked input. The six THREAD module signatures were tested using a minimum gene-set size of 10, a maximum gene-set size of 500, and `eps = 0`. The random-number seed was fixed at 42. Nominal enrichment $P$ values were adjusted across the six module signatures using the Benjamini–Hochberg procedure, and modules with an adjusted $P < 0.05$ were considered significantly enriched. Positive normalized enrichment scores indicate enrichment toward genes upregulated in AD, whereas negative scores indicate enrichment toward genes downregulated in AD.

Leading-edge genes were extracted directly from the `fgsea` result for each module. Each leading-edge gene was linked to its THREAD signature rank and score and to its limma rank, log-fold change, moderated $t$-statistic, nominal $P$ value, and gene-level Benjamini–Hochberg-adjusted $P$ value. Module-level fgsea adjusted $P$ values and gene-level limma adjusted $P$ values were retained as distinct statistical quantities. Complete module-level enrichment statistics, signature-mapping results, sample metadata, and leading-edge gene annotations are provided in Supplementary Data Sx. Enrichment curves for modules C2, C4, C5, and C6 are shown in Supplementary Fig. 4, whereas the C1 and C3 enrichment profiles are presented in the main figure.

##### 3.2 Clinical validation of CRC module genes in TCGA-COAD

Clinical validation of the CRC tumor-associated gene modules was performed using TCGA-COAD clinical annotations and matched gene-expression profiles. The validation set consisted of the top 10 protein-coding genes selected from each of the two CRC tumor-associated THREAD modules, yielding a prespecified panel of 20 genes. The selected genes were reconciled with the expression matrix using gene symbols and Ensembl identifiers. The expression values in the source matrix were already represented on a $\log_2$ scale and were therefore used without an additional logarithmic transformation. The THREAD module assignment, within-module rank, THREAD score, Ensembl identifier, and expression-mapping status of each selected gene were retained in the supplementary source data.

Normal samples were identified from the TCGA sample-type annotation or from the normal-tissue barcode suffix. For the stage-stratified analysis, pathological stage annotations were harmonized by combining substages IA and IB into Stage I, substages IIA–IIC into Stage II, substages IIIA–IIIC into Stage III, and substages IVA and IVB into Stage IV. Samples with missing or unmappable pathological-stage annotations were excluded from the stage-stratified analysis. The ordered groups were defined as Normal, Stage I, Stage II, Stage III, and Stage IV. For the binary tumor–normal analysis, matched samples not classified as normal were assigned to the tumor group.

For each gene, expression distributions across Normal and stages I–IV were summarized using the sample count, mean, standard deviation, median, interquartile range, minimum, and maximum. An omnibus Kruskal–Wallis test was used to assess whether expression differed among the five ordered groups. A two-sided Jonckheere–Terpstra test was then used to assess an ordered trend from Normal through Stage IV. Because an association across Normal and tumor stages can be driven primarily by the transition from normal tissue to tumor, a second two-sided Jonckheere–Terpstra test was performed after restricting the data to Stages I–IV. Spearman correlations between the ordered stage code and gene expression were reported to indicate the direction of each trend. Kruskal–Wallis, Normal-to-Stage IV Jonckheere–Terpstra, and Stage I-to-Stage IV Jonckheere–Terpstra $P$ values were adjusted separately across the 20 tested genes using the Benjamini–Hochberg procedure.

Tumor–normal expression differences were evaluated on the existing $\log_2$ expression scale. For each gene, the tumor-minus-normal difference in mean expression was reported as a $\log_2$ fold-change estimate together with its 95% confidence interval from a two-sample Welch $t$-test. A Wilcoxon rank-sum test was additionally used to provide a distribution-free comparison between tumor and normal samples. Welch-test and Wilcoxon-test $P$ values were adjusted separately across the 20 genes using the Benjamini–Hochberg procedure.

A directional area under the receiver-operating-characteristic curve was calculated from the Mann–Whitney rank statistic. The directional AUC was defined as the probability that a randomly selected tumor sample had higher expression than a randomly selected normal sample. Consequently, values greater than 0.5 indicate higher expression in tumor, whereas values below 0.5 indicate lower expression in tumor and do not necessarily indicate weak discrimination. To summarize direction-independent separation, a discrimination AUC was defined as

$$\mathrm{AUC}_{\mathrm{discrimination}} = \max\left(\mathrm{AUC}_{\mathrm{directional}}, 1 - \mathrm{AUC}_{\mathrm{directional}}\right).$$

Confidence intervals for both AUC measures were estimated using 2,000 stratified bootstrap resamples, with normal and tumor samples resampled separately and the random-number seed fixed for reproducibility.

Four representative genes were displayed individually in the main stage-stratified figure, whereas expression profiles for the complete prespecified 20-gene panel are shown in Supplementary Fig. 6. Complete sample-level expression data, stage-specific summaries, stage-trend statistics, tumor–normal effect estimates, and directional and discrimination AUC results are provided in Supplementary Data Sx. The complete stage-trend and tumor–normal results are additionally summarized in Supplementary Tables 4 and 5.

**Supplementary Table 1: Prior specification and empirical-Bayes hyperparameter calibration in THREAD.**

| **Parameter** | **Prior component** | **Default specification** |
|---|---|---|
| $\alpha$ | Dirichlet process concentration | 0.1 |
| $\alpha_0$ | Inverse-Gamma shape | 2 |
| $\beta_0$ | Inverse-Gamma scale | $\max\{\widehat{v}_\gamma/2,\ 10^{-16}\}$ |
| $r$ | Gamma shape | 1 |
| $\delta$ | Gamma rate | 1 |

*Note.* The scale hyperparameter $\beta_0$ is estimated from the input data before MCMC using a weighted ridge approximation. Here, $\widehat{v}_\gamma$ denotes the estimated coefficient variance, obtained by averaging valid initial-cluster estimates or, when unavailable, from a global weighted ridge fit. The remaining hyperparameters are assigned the listed default values.

**Supplementary Table 2a: Main-grid simulation performance across 18 settings.**

| $K_0$ | $G$ | $s$ | THREAD macro-F1 | Bayes-only macro-F1 | FGLS-only macro-F1 | MAGMA macro-F1 | RolyPoly macro-F1 | THREAD ARI |
|---|---|---|---|---|---|---|---|---|
| 1 | 500 | 1 | 0.981 (0.066) | 0.933 (0.100) | 0.981 (0.066) | 0.992 (0.040) | 0.940 (0.093) | 1.000 (0.000) |
| 1 | 500 | 10 | 0.981 (0.066) | 0.941 (0.097) | 0.981 (0.066) | 0.992 (0.040) | 0.944 (0.091) | 1.000 (0.000) |
| 1 | 1000 | 1 | 0.976 (0.066) | 0.948 (0.103) | 0.976 (0.066) | 0.992 (0.040) | 0.940 (0.093) | 1.000 (0.000) |
| 1 | 1000 | 10 | 0.976 (0.066) | 0.945 (0.108) | 0.976 (0.066) | 0.992 (0.040) | 0.916 (0.100) | 1.000 (0.000) |
| 1 | 3000 | 1 | 0.983 (0.062) | 0.940 (0.105) | 0.983 (0.062) | 0.984 (0.055) | 0.944 (0.091) | 0.940 (0.240) |
| 1 | 3000 | 10 | 0.987 (0.056) | 0.930 (0.103) | 0.987 (0.056) | 0.984 (0.055) | 0.936 (0.094) | 0.960 (0.198) |
| 3 | 500 | 1 | 0.944 (0.063) | 0.917 (0.081) | 0.925 (0.072) | 0.565 (0.008) | 0.567 (0.007) | 0.590 (0.038) |
| 3 | 500 | 10 | 0.944 (0.063) | 0.922 (0.076) | 0.920 (0.076) | 0.565 (0.008) | 0.567 (0.007) | 0.591 (0.039) |
| 3 | 1000 | 1 | 0.958 (0.053) | 0.928 (0.065) | 0.932 (0.065) | 0.570 (0.004) | 0.571 (0.002) | 0.586 (0.029) |
| 3 | 1000 | 10 | 0.958 (0.050) | 0.925 (0.065) | 0.932 (0.071) | 0.570 (0.004) | 0.571 (0.002) | 0.587 (0.027) |
| 3 | 3000 | 1 | 0.955 (0.068) | 0.917 (0.085) | 0.939 (0.079) | 0.571 (0.000) | 0.571 (0.000) | 0.589 (0.017) |
| 3 | 3000 | 10 | 0.953 (0.070) | 0.913 (0.090) | 0.936 (0.080) | 0.571 (0.000) | 0.571 (0.000) | 0.590 (0.016) |
| 5 | 500 | 1 | 0.945 (0.059) | 0.928 (0.058) | 0.872 (0.070) | 0.571 (0.005) | 0.571 (0.000) | 0.512 (0.033) |
| 5 | 500 | 10 | 0.935 (0.059) | 0.913 (0.067) | 0.866 (0.069) | 0.571 (0.005) | 0.571 (0.000) | 0.513 (0.034) |
| 5 | 1000 | 1 | 0.948 (0.050) | 0.921 (0.057) | 0.878 (0.075) | 0.571 (0.000) | 0.571 (0.000) | 0.523 (0.027) |
| 5 | 1000 | 10 | 0.950 (0.044) | 0.923 (0.050) | 0.888 (0.077) | 0.571 (0.000) | 0.571 (0.005) | 0.525 (0.027) |
| 5 | 3000 | 1 | 0.960 (0.040) | 0.935 (0.050) | 0.919 (0.062) | 0.571 (0.000) | 0.571 (0.000) | 0.526 (0.016) |
| 5 | 3000 | 10 | 0.954 (0.036) | 0.933 (0.047) | 0.920 (0.059) | 0.571 (0.000) | 0.571 (0.000) | 0.527 (0.017) |

| $K_0$ | $G$ | $s$ | THREAD macro-F1 | Bayes-only macro-F1 | FGLS-only macro-F1 | MAGMA macro-F1 | RolyPoly macro-F1 | THREAD ARI |
|---|---|---|---|---|---|---|---|---|

*Note.* Values are mean (SD) across 50 replicates. $K_0$ denotes the true number of latent programs, $G$ denotes the number of genes, and $s$ denotes the coefficient scale factor. THREAD denotes the final dual-significance rule requiring both Bayesian and frequentist evidence.

#### Supplementary Table 2b1: Rare-program and null-background stress-test classification performance.

| Scenario | Setting | THREAD macro-F1 | Bayes-only macro-F1 | FGLS-only macro-F1 | MAGMA macro-F1 | RolyPoly macro-F1 | THREAD ARI |
|---|---|---|---|---|---|---|---|
| Rare program | 80/10/10 programs | 0.964 (0.049) | 0.946 (0.064) | 0.946 (0.062) | 0.566 (0.007) | 0.568 (0.007) | 0.758 (0.014) |
| Null background | null = 0.25 | 0.984 (0.035) | 0.340 (0.129) | 0.842 (0.133) | 0.566 (0.010) | 0.571 (0.000) | 0.486 (0.061) |
| Null background | null = 0.50 | 0.980 (0.040) | 0.525 (0.168) | 0.836 (0.100) | 0.539 (0.023) | 0.569 (0.006) | 0.277 (0.184) |
| Null background | null = 0.75 | 0.972 (0.050) | 0.636 (0.229) | 0.559 (0.282) | 0.350 (0.222) | 0.560 (0.011) | 0.235 (0.373) |
| Null background | null = 0.875 | 0.987 (0.031) | – | – | 0.127 (0.199) | 0.537 (0.034) | 0.363 (0.475) |
| Null background | null = 0.9375 | 0.985 (0.032) | – | – | 0.070 (0.146) | 0.441 (0.138) | 0.658 (0.475) |

*Note.* Values are mean (SD) across 50 replicates. The rare-program setting used an imbalanced three-program architecture with program proportions 80%, 10% and 10%. Null-background settings used three signal programs plus an all-zero null program. ARI was computed for THREAD module recovery. Dashes indicate metrics not available in the corresponding summary file.

#### Supplementary Table 2b2: False-positive burden under null-background settings.

| Null fraction | THREAD mean FP | Bayes-only mean FP | FGLS-only mean FP | MAGMA mean FP | RolyPoly mean FP |
|---|---|---|---|---|---|
| 0.25 | 1.6 | 25.9 | 2.7 | 8.4 | 9.0 |
| 0.50 | 0.6 | 12.5 | 2.5 | 5.7 | 8.7 |
| 0.75 | 2.8 | 6.9 | 11.6 | 2.6 | 7.6 |
| 0.875 | 3.8 | 5.0 | 27.6 | 0.7 | 6.0 |
| 0.9375 | 0.8 | 1.3 | 13.7 | 0.5 | 3.6 |

*Note.* FP denotes the mean number of false-positive program–subtype calls across 50 replicates. THREAD denotes the final dual-significance rule. Bayes-only uses the Bayesian lFSR criterion alone, whereas FGLS-only uses the frequentist FDR criterion alone.

==================================================================

**Supplementary Table 2c: Representative feature-level coefficient recovery in the three-program setting.**

| $s$ | True coefficient | FGLS estimate mean (SD) | Mean bias | RMSE |
|---|---|---|---|---|
| 1 | 0.000 | 0.0000 (0.0002) | 0.0000 | 0.0002 |
| 1 | 0.005 | 0.0050 (0.0002) | 0.0000 | 0.0002 |
| 1 | 0.010 | 0.0100 (0.0003) | 0.0000 | 0.0003 |
| 1 | 0.020 | 0.0200 (0.0002) | 0.0000 | 0.0002 |
| 1 | 0.030 | 0.0302 (0.0003) | 0.0002 | 0.0004 |
| 10 | 0.000 | -0.0002 (0.0020) | -0.0002 | 0.0020 |
| 10 | 0.050 | 0.0500 (0.0019) | 0.0000 | 0.0019 |
| 10 | 0.100 | 0.0997 (0.0027) | -0.0003 | 0.0026 |
| 10 | 0.200 | 0.2004 (0.0021) | 0.0004 | 0.0021 |
| 10 | 0.300 | 0.3021 (0.0031) | 0.0021 | 0.0037 |

*Note.* The table summarizes representative $K_0 = 3$, $G = 1000$ settings at coefficient scale factors $s = 1$ and $s = 10$. Rows are grouped by true coefficient values across program–subtype pairs and 50 replicates. Bias was calculated as the estimated coefficient minus the true coefficient. RMSE denotes root mean squared error.

**Supplementary Table 3: Independent validation of AD-associated gene modules in GSE109887.**

| Module | Mapped genes | NES | Nominal $P$ | FDR | Leading-edge size |
|---|---|---|---|---|---|
| C1 | 182/200 (91.0%) | -1.53 | 0.003 | 0.008 | 67 |
| C2 | 170/200 (85.0%) | 1.14 | 0.167 | 0.201 | 54 |
| C3 | 159/200 (79.5%) | 1.37 | 0.020 | 0.030 | 48 |
| C4 | 139/185 (75.1%) | -1.55 | 0.005 | 0.010 | 48 |
| C5 | 169/200 (84.5%) | -1.58 | 0.002 | 0.008 | 66 |
| C6 | 19/27 (70.4%) | -1.20 | 0.221 | 0.221 | 6 |

**Supplementary Table 4: Stage-dependent expression trends of THREAD-prioritized genes in TCGA-COAD.**

| Gene | Module | Rank | KW $q$ | JT $q$ (N–IV) | $\rho$ (N–IV) | JT $q$ (I–IV) | $\rho$ (I–IV) |
|---|---|---|---|---|---|---|---|
| FTH1 | C1 | 1 | $2.96 \times 10^{-4}$ | 0.056 | -0.13 | 0.995 | 0.02 |
| FTL | C1 | 2 | $6.65 \times 10^{-5}$ | 0.001 | 0.20 | 0.446 | 0.07 |
| ACTG1 | C1 | 3 | 0.991 | 0.993 | -0.00 | 0.995 | -0.00 |
| NFKBIA | C1 | 4 | $5.34 \times 10^{-7}$ | 0.032 | -0.13 | 0.346 | 0.09 |
| CD74 | C1 | 5 | $1.16 \times 10^{-5}$ | $4.13 \times 10^{-5}$ | -0.25 | 0.175 | -0.12 |
| HSP90AB1 | C1 | 6 | $5.20 \times 10^{-17}$ | $1.51 \times 10^{-10}$ | 0.37 | 0.305 | 0.10 |
| BTG1 | C1 | 7 | $2.74 \times 10^{-10}$ | $1.13 \times 10^{-4}$ | -0.23 | 0.995 | 0.01 |
| VIM | C1 | 8 | $2.92 \times 10^{-7}$ | 0.181 | -0.08 | 0.175 | 0.14 |
| SRGN | C1 | 9 | $1.21 \times 10^{-5}$ | 0.002 | -0.19 | 0.995 | -0.02 |
| KLF6 | C1 | 10 | $2.64 \times 10^{-6}$ | 0.002 | -0.19 | 0.995 | 0.00 |
| B2M | C2 | 1 | $5.16 \times 10^{-11}$ | $1.55 \times 10^{-8}$ | -0.33 | 0.175 | -0.12 |
| EEF1A1 | C2 | 2 | 0.288 | 0.868 | 0.02 | 0.754 | -0.04 |
| TMSB10 | C2 | 3 | 0.009 | 0.102 | 0.10 | 0.995 | -0.02 |
| ACTB | C2 | 4 | 0.013 | 0.952 | -0.01 | 0.175 | 0.12 |
| PTMA | C2 | 5 | $8.89 \times 10^{-12}$ | $2.82 \times 10^{-5}$ | 0.25 | 0.995 | -0.01 |
| TPT1 | C2 | 6 | 0.053 | 0.007 | 0.16 | 0.175 | 0.13 |
| S100A6 | C2 | 7 | 0.822 | 0.550 | 0.04 | 0.995 | 0.00 |
| GAPDH | C2 | 8 | $2.46 \times 10^{-6}$ | 0.061 | 0.11 | 0.305 | -0.09 |
| HSP90AA1 | C2 | 9 | $2.53 \times 10^{-9}$ | 0.002 | 0.19 | 0.603 | -0.06 |
| EIF1 | C2 | 10 | 0.073 | 0.150 | 0.09 | 0.266 | 0.11 |

*Note.* KW, Kruskal–Wallis test across Normal and stages I–IV; JT (N–IV), two-sided Jonckheere–Terpstra test across Normal and stages I–IV; JT (I–IV), the corresponding test restricted to tumor stages I–IV. $\rho$ denotes the Spearman correlation between the ordered stage code and $\log_2$ expression. Positive values indicate increasing expression and negative values indicate decreasing expression. $P$ values were adjusted separately across the 20 genes within each test family using the Benjamini–Hochberg method.

**Supplementary Table 5: Tumor–normal expression differences and discrimination metrics for THREAD-prioritized genes in TCGA-COAD.**

| Gene | Module | Rank | $\log_2$FC [95% CI] | Welch $t$ $q$ | Directional AUC | Discrimination AUC | Wilcoxon $q$ |
|---|---|---|---|---|---|---|---|
| FTH1 | C1 | 1 | -0.49 [-0.78, -0.20] | 0.002 | 0.313 | 0.687 | $2.32 \times 10^{-5}$ |
| FTL | C1 | 2 | 0.45 [0.26, 0.63] | $1.63 \times 10^{-5}$ | 0.709 | 0.709 | $2.32 \times 10^{-5}$ |
| ACTG1 | C1 | 3 | 0.00 [-0.18, 0.18] | 0.993 | 0.499 | 0.501 | 0.985 |
| NFKBIA | C1 | 4 | -0.68 [-0.85, -0.52] | $3.65 \times 10^{-11}$ | 0.214 | 0.786 | $1.05 \times 10^{-8}$ |
| CD74 | C1 | 5 | -0.95 [-1.23, -0.66] | $9.21 \times 10^{-9}$ | 0.272 | 0.728 | $4.12 \times 10^{-6}$ |
| HSP90AB1 | C1 | 6 | 1.16 [1.01, 1.32] | $2.69 \times 10^{-21}$ | 0.950 | 0.950 | $2.32 \times 10^{-19}$ |
| BTG1 | C1 | 7 | -0.85 [-1.03, -0.67] | $6.65 \times 10^{-13}$ | 0.147 | 0.853 | $1.26 \times 10^{-12}$ |
| VIM | C1 | 8 | -0.93 [-1.15, -0.72] | $4.99 \times 10^{-13}$ | 0.227 | 0.773 | $3.80 \times 10^{-8}$ |
| SRGN | C1 | 9 | -1.10 [-1.33, -0.88] | $2.29 \times 10^{-16}$ | 0.234 | 0.766 | $6.86 \times 10^{-8}$ |
| KLF6 | C1 | 10 | -0.73 [-0.95, -0.51] | $2.71 \times 10^{-8}$ | 0.227 | 0.773 | $3.80 \times 10^{-8}$ |
| B2M | C2 | 1 | -1.12 [-1.35, -0.88] | $6.65 \times 10^{-13}$ | 0.143 | 0.857 | $8.67 \times 10^{-13}$ |
| EEF1A1 | C2 | 2 | 0.11 [-0.05, 0.26] | 0.203 | 0.566 | 0.566 | 0.200 |
| TMSB10 | C2 | 3 | 0.38 [0.19, 0.57] | $2.58 \times 10^{-4}$ | 0.672 | 0.672 | $5.13 \times 10^{-4}$ |
| ACTB | C2 | 4 | -0.26 [-0.43, -0.09] | 0.005 | 0.357 | 0.643 | 0.004 |
| PTMA | C2 | 5 | 0.73 [0.60, 0.86] | $1.44 \times 10^{-16}$ | 0.879 | 0.879 | $4.03 \times 10^{-14}$ |
| TPT1 | C2 | 6 | 0.18 [0.02, 0.35] | 0.034 | 0.590 | 0.590 | 0.077 |
| S100A6 | C2 | 7 | 0.13 [-0.08, 0.34] | 0.240 | 0.561 | 0.561 | 0.226 |
| GAPDH | C2 | 8 | 0.54 [0.38, 0.69] | $3.90 \times 10^{-9}$ | 0.767 | 0.767 | $6.86 \times 10^{-8}$ |
| HSP90AA1 | C2 | 9 | 0.78 [0.61, 0.94] | $5.00 \times 10^{-13}$ | 0.835 | 0.835 | $1.47 \times 10^{-11}$ |
| EIF1 | C2 | 10 | 0.05 [-0.11, 0.22] | 0.545 | 0.511 | 0.511 | 0.867 |

*Note.* $\log_2$FC denotes the difference in mean $\log_2$ expression between tumor and normal samples. Directional AUC is the probability that a randomly selected tumor sample has higher expression than a randomly selected normal sample; values below 0.5 therefore indicate lower expression in tumor rather than absence of discrimination. Discrimination AUC was defined as $\max(\mathrm{AUC}, 1 - \mathrm{AUC})$. Welch and Wilcoxon $P$ values were adjusted separately across the 20 genes using the Benjamini–Hochberg method.

**Supplementary Table 6: Cellular composition of the GSE176078 breast-cancer discovery cohort.**

| Major lineage | ER+ | HER2+ | TNBC | Total |
|---|---|---|---|---|
| B-cells | 606 (1.6%) | 624 (3.2%) | 1,976 (4.6%) | 3,206 (3.2%) |
| CAFs | 2,573 (6.7%) | 1,449 (7.5%) | 2,551 (6.0%) | 6,573 (6.6%) |
| Cancer Epithelial | 11,878 (31.1%) | 1,775 (9.2%) | 10,836 (25.5%) | 24,489 (24.5%) |
| Endothelial | 5,206 (13.6%) | 1,016 (5.3%) | 1,383 (3.3%) | 7,605 (7.6%) |
| Myeloid | 1,831 (4.8%) | 1,422 (7.4%) | 6,422 (15.1%) | 9,675 (9.7%) |
| Normal Epithelial | 2,430 (6.4%) | 968 (5.0%) | 957 (2.3%) | 4,355 (4.4%) |
| Plasmablasts | 631 (1.7%) | 226 (1.2%) | 2,667 (6.3%) | 3,524 (3.5%) |
| PVL | 3,265 (8.5%) | 894 (4.6%) | 1,264 (3.0%) | 5,423 (5.4%) |
| T-cells | 9,821 (25.7%) | 10,937 (56.6%) | 14,456 (34.0%) | 35,214 (35.2%) |

*Note.* Values are numbers of retained cells, with percentages calculated relative to the total number of cells within the corresponding clinical subtype. Percentages in the Total column are relative to all retained cells. ER+, estrogen-receptor-positive breast cancer; HER2+, human epidermal growth factor receptor 2-positive breast cancer; TNBC, triple-negative breast cancer; CAF, cancer-associated fibroblast; PVL, perivascular-like cells.

**Supplementary Table 7: Adjusted-significant cell types identified by three baseline methods across the real-data benchmark datasets.**

| Dataset | MAGMA | RolyPoly | scPagwas |
|---|---|---|---|
| AD | 0/31 | 5/31 | 6/31 |
| BIP | 21/21 | 0/21 | 4/21 |
| SCZ | 21/21 | 3/21 | 6/21 |
| CRC normal | 0/31 | 0/31 | 2/31 |
| CRC tumor | 0/31 | 0/31 | 6/30* |
| BRCA tumor | 29/29 | 0/29 | 0/29 |

*Note.* Entries indicate the number of cell types significant after Benjamini–Hochberg correction ($Q_{\mathrm{BH}} < 0.05$) divided by the number of cell types tested by each method. *In CRC tumor, Mast cells were not tested by scPagwas because only three Mast cells were available, below the method's minimum cell-count threshold of five.

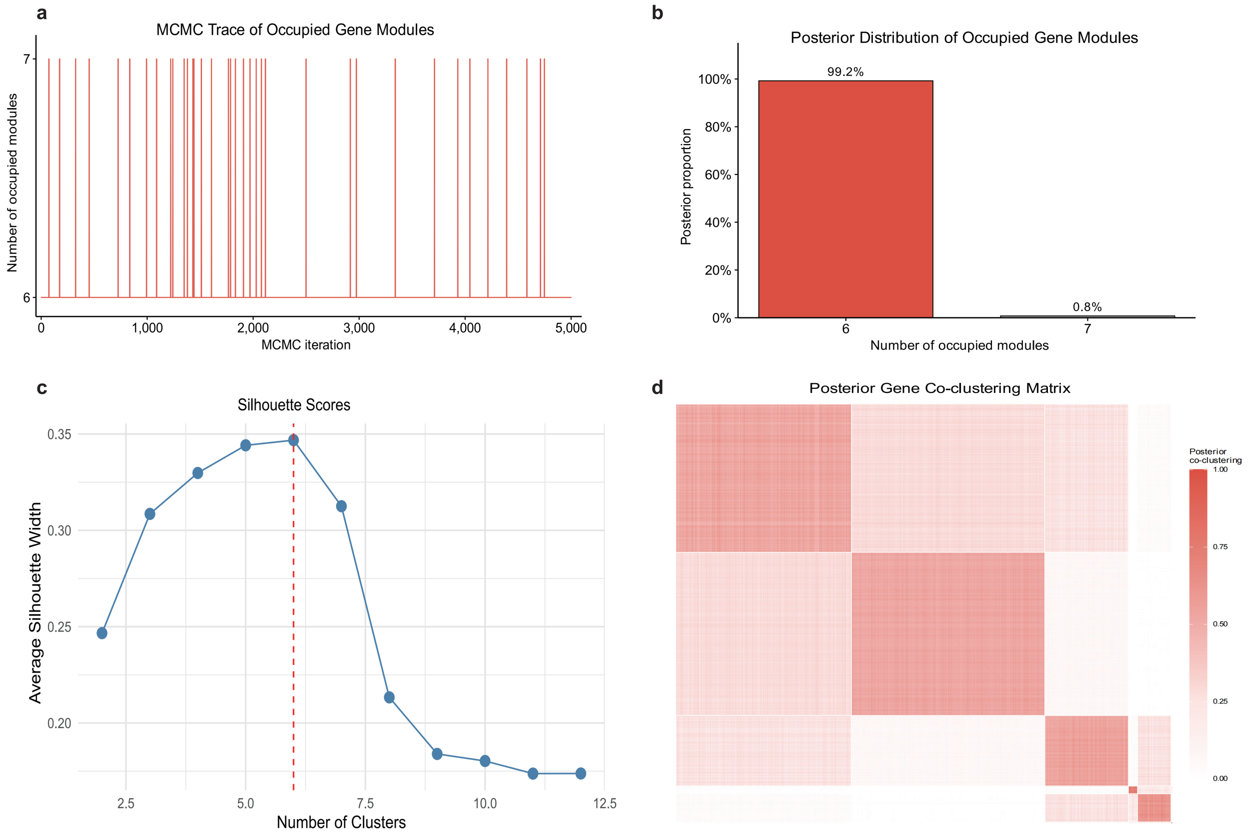

**Supplementary Figure 1: MCMC convergence and posterior partition diagnostics for the Alzheimer's disease (AD) analysis.** **a**, Trace plot of the number of active, occupied non-micro gene modules ($K^{(t)}$) across 5,000 MCMC iterations. **b**, Empirical posterior distribution of the number of occupied gene modules across retained post-burn-in draws, showing strong posterior concentration at the mode $K_{\mathrm{mode}} = 6$ (99.2% of retained draws). **c**, Average silhouette width profile evaluated across candidate cluster counts ($K = 2, \ldots, 12$) via partitioning around medoids (PAM) on the posterior dissimilarity matrix $\boldsymbol{\Delta} = \mathbf{1}\mathbf{1}^\top - \boldsymbol{Q}$. The vertical red dashed line indicates the optimal selected module count ($\hat{K} = 6$, maximum average silhouette width $= 0.347$). **d**, Empirical posterior gene co-clustering similarity matrix $\boldsymbol{Q} \in [0, 1]^{G\times G}$, with rows and columns ordered by the final $\hat{K} = 6$ point partition, demonstrating sharp block-diagonal coherence and high partition stability.

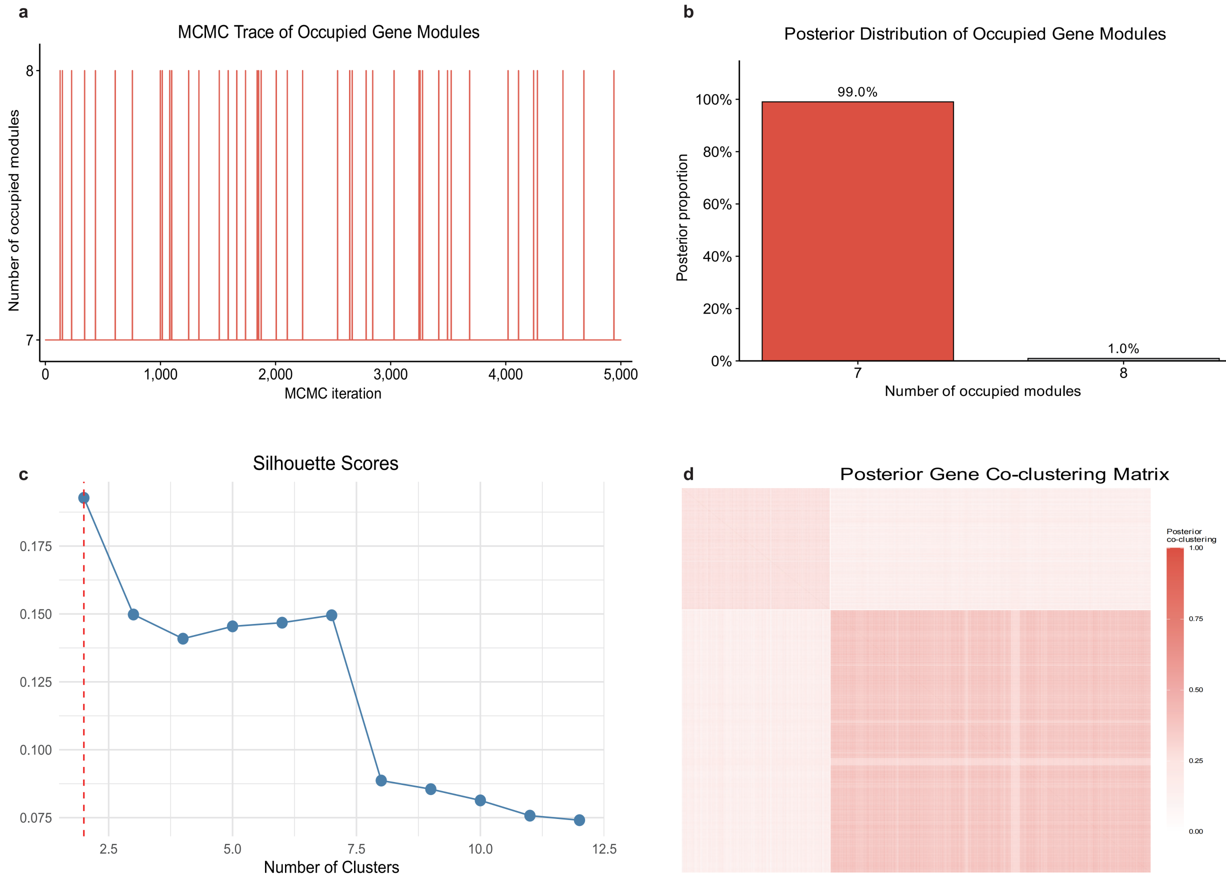

**Supplementary Figure 2: MCMC convergence and posterior partition diagnostics for the colorectal cancer (CRC) analysis.** **a**, Trace plot of the number of occupied non-micro gene modules ($K^{(t)}$) across 5,000 MCMC iterations. **b**, Empirical posterior distribution of the number of occupied modules across retained post-burn-in draws, showing 99.0% posterior mass at seven raw modules prior to silhouette-guided PAM consolidation. **c**, Average silhouette width profile across candidate cluster counts ($K = 2, \ldots, 12$) on the posterior dissimilarity matrix $\boldsymbol{\Delta}$, with the red dashed line marking the global maximum at $\hat{K} = 2$ modules for the tumour compartment, and a secondary local plateau at $K = 7$ corresponding to the finer-grained normal-colon partition. **d**, Posterior gene co-clustering matrix $\boldsymbol{Q}$ ordered by the final point partition, illustrating the dominant two-compartment block structure separating the tumour-intrinsic epithelial program (C1) from the microenvironmental immune–stromal program (C2).

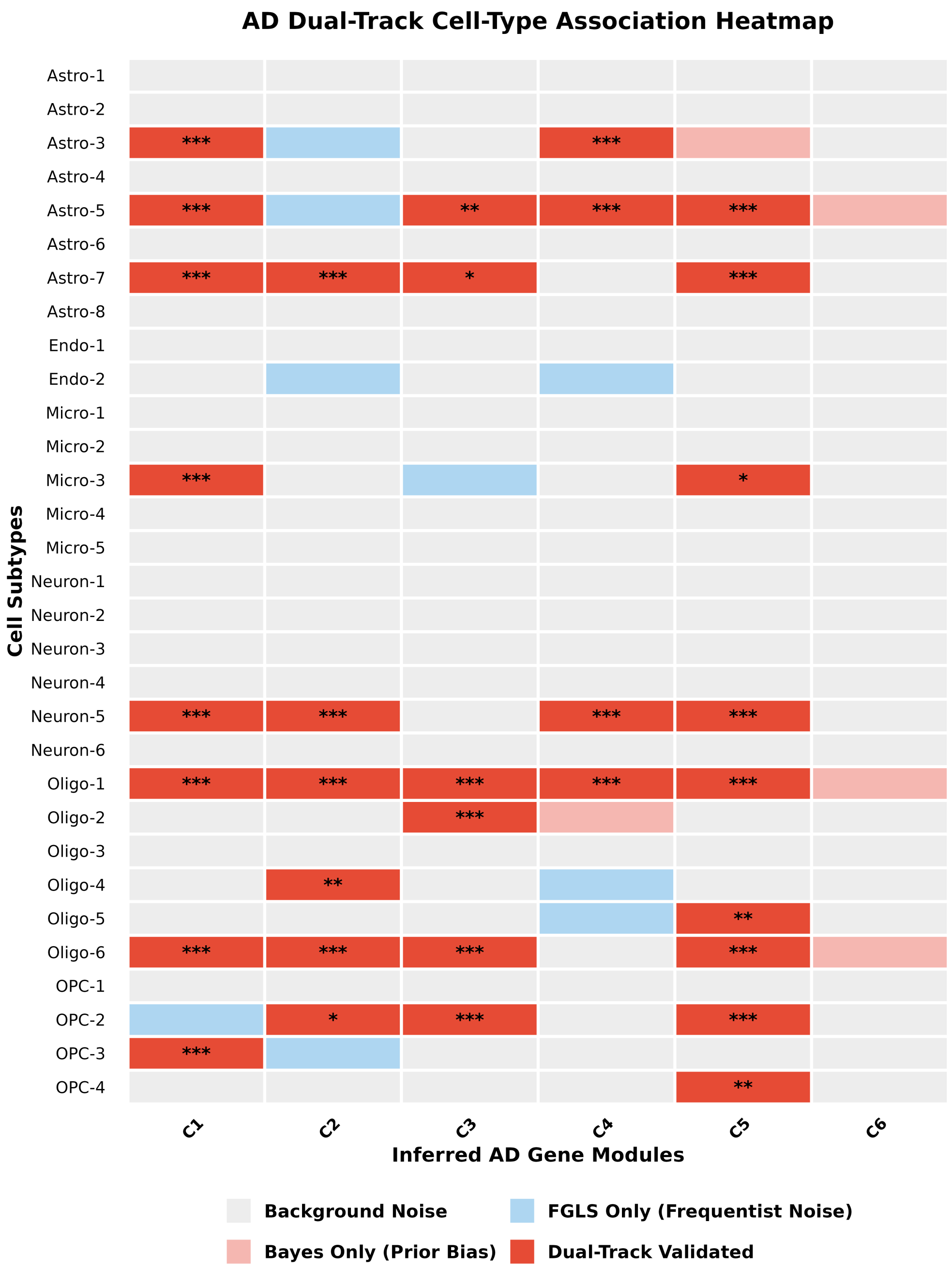

**Supplementary Figure 3: Module-resolved cell-subtype association landscape under the dual-significance criterion in Alzheimer's disease.** Heatmap displaying the statistical support for associations between the 31 entorhinal-cortex cell subtypes (rows, spanning astrocyte, endothelial, microglial, neuronal, oligodendrocyte and OPC lineages from GSE138852) and the six inferred AD gene modules C1–C6 (columns). Each module–subtype pair is classified into one of four categories: background noise (grey; neither criterion met), frequentist-only calls (light blue; FGLS BH-adjusted $q < 0.05$ but Bayesian lFSR $\geq 0.05$), Bayesian-only calls (light pink; lFSR $< 0.05$ but FGLS $q \geq 0.05$), and dual-significant associations (dark red; satisfying both lFSR $< 0.05$ and FGLS $q < 0.05$). Asterisks within dual-significant cells denote the frequentist FGLS FDR significance level ($^{*}q < 0.05$, $^{**}q < 0.01$, $^{***}q < 0.001$). Module C6 (Lipid·APOE) exhibits no dual-significant cell-subtype specificity, consistent with a cell-type-agnostic genetic risk core.

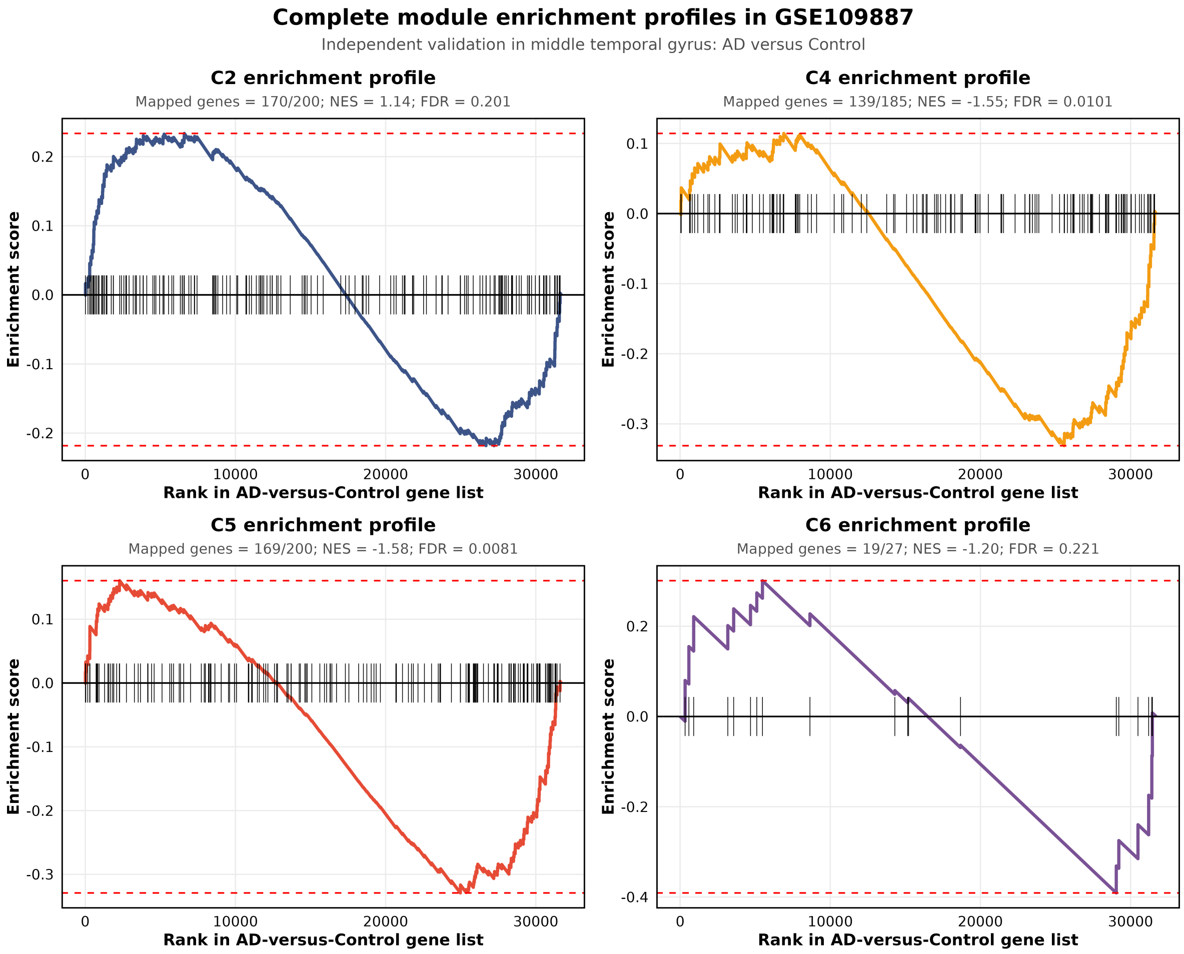

**Supplementary Figure 4: Independent transcriptomic validation of AD gene modules C2, C4, C5 and C6 in the external GSE109887 middle temporal gyrus cohort.** Gene-set enrichment analysis (`fgsea`) running-sum profiles for the top-ranked module gene signatures evaluated against the full transcriptome ranked by `limma` moderated $t$-statistics comparing AD cases ($n = 46$) with non-demented controls ($n = 32$; positive ranks correspond to upregulation in AD and negative ranks to downregulation in AD). Vertical black ticks mark the positions of mapped module signature genes along the ranked gene list, and horizontal red dashed lines indicate the maximum deviation from zero defining the enrichment score. Shown are the synaptic/chromatin module C2 (top left; 170/200 mapped genes, NES = 1.14, BH-adjusted FDR = 0.201), the proteostasis module C4 (top right; 139/185 mapped genes, NES = −1.55, FDR = 0.0101), the RNA/proteostasis module C5 (bottom left; 169/200 mapped genes, NES = −1.58, FDR = 0.0081), and the compact cell-type-agnostic Lipid·APOE module C6 (bottom right; 19/27 mapped genes, NES = −1.20, FDR = 0.221). Enrichment profiles for modules C1 and C3 are shown in Fig. 3F of the main text.

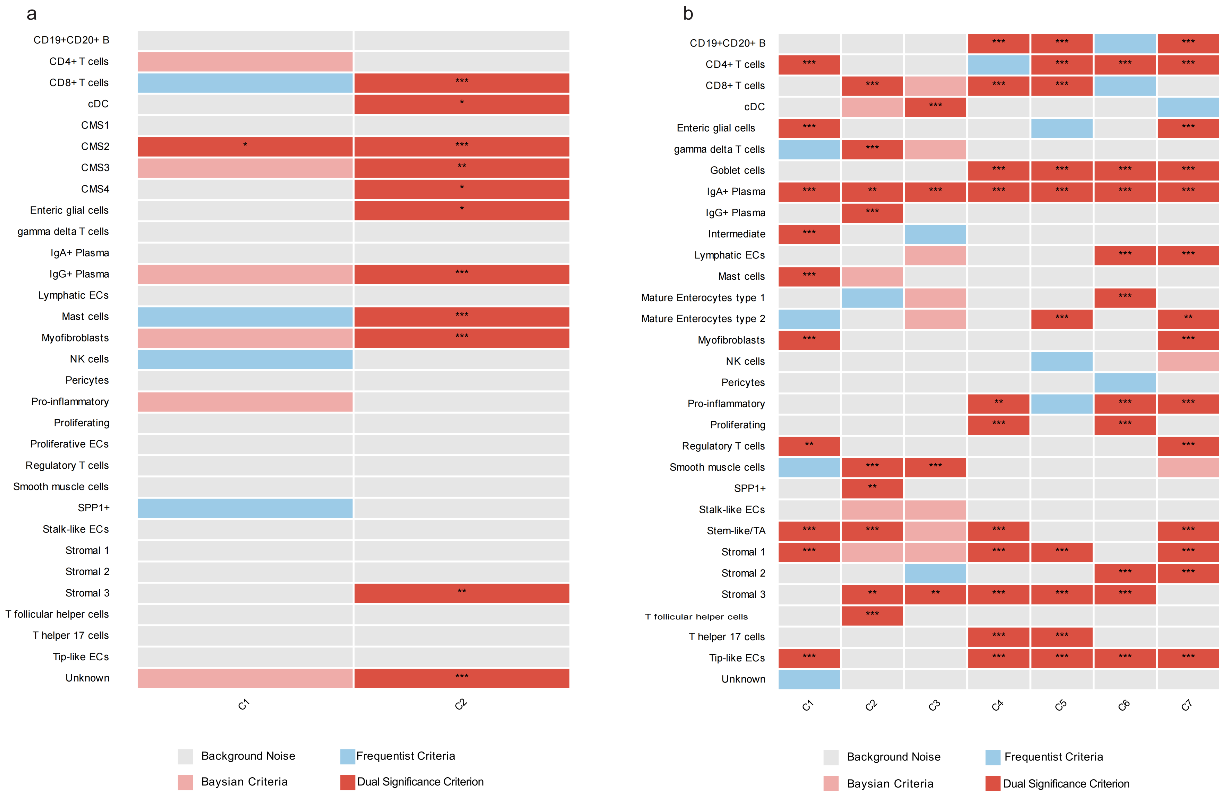

**Supplementary Figure 5: Dual-significance cell-subtype association heatmaps for colorectal cancer (CRC) across tumour and matched normal colon compartments. a**, Module–cell-subtype association matrix in the CRC tumour compartment (GSE132465) across the two inferred modules C1 and C2. Module C1 is exclusively associated with the canonical epithelial malignant subtype CMS2, whereas module C2 captures a multicellular microenvironmental program spanning $CD8^{+}$ T cells, conventional dendritic cells (cDC), $IgG^{+}$ plasma cells, mast cells, myofibroblasts, Stromal 3 cells, enteric glial cells, and the CMS2/CMS3/CMS4 malignant states. **b**, Module–cell-subtype association matrix in the matched normal-colon atlas across seven inferred modules (C1–C7), resolving homeostatic epithelial, stromal, vascular and mucosal-immune compartments. Colour coding indicates background noise (grey), Bayesian-only calls (lFSR $< 0.05$; light pink), frequentist-only calls (FGLS $q < 0.05$; light blue), and dual-significant associations (lFSR $< 0.05$ and FGLS $q < 0.05$; dark red). Asterisks indicate FGLS FDR thresholds ($^{*}q < 0.05$, $^{**}q < 0.01$, $^{***}q < 0.001$).

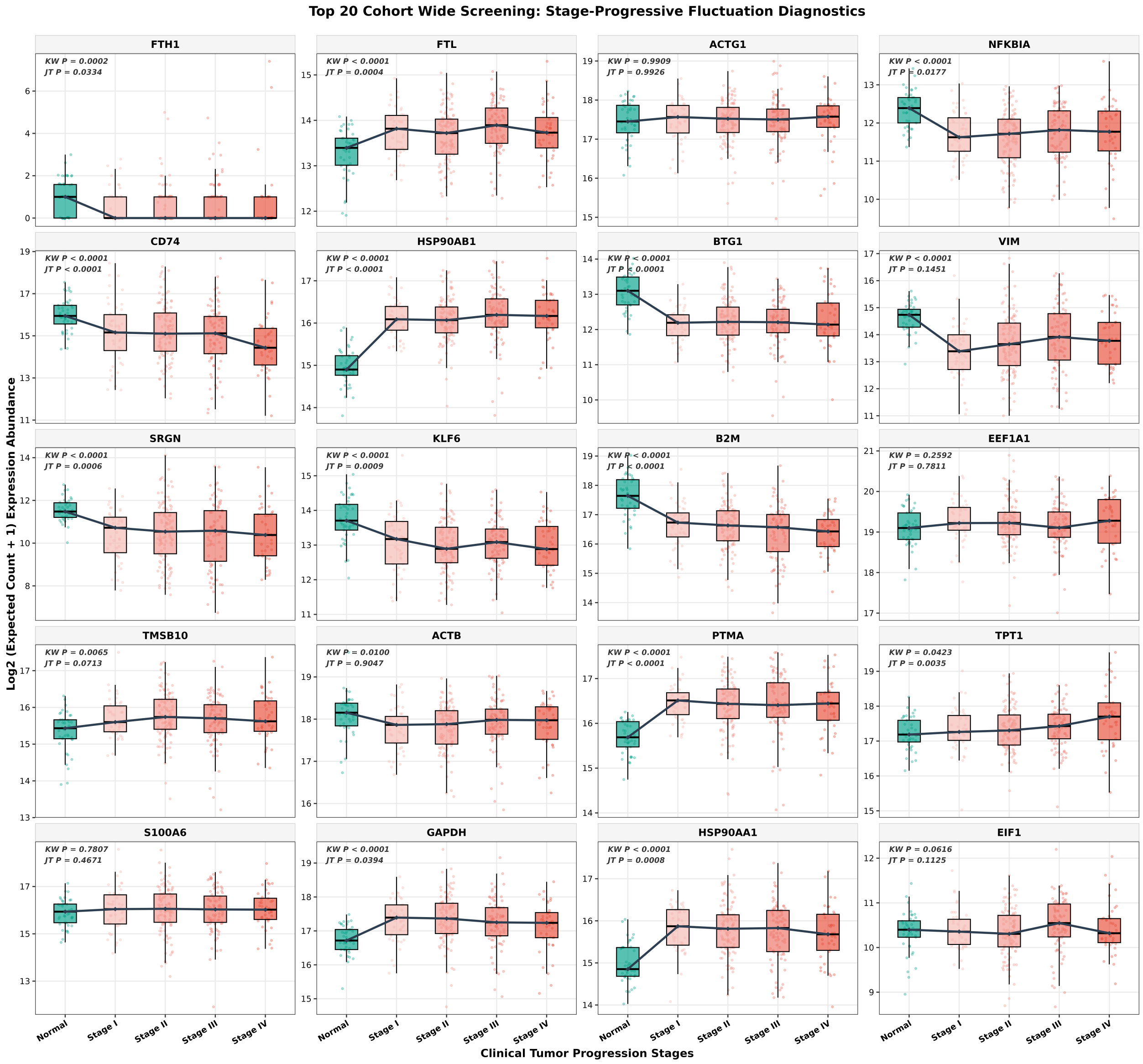

**Supplementary Figure 6: Stage-stratified expression profiles of the top 20 THREAD-prioritized CRC module genes in the independent TCGA-COAD cohort.** Box-and-jitter plots showing $\log_2$(expected count + 1) expression abundance across normal colon tissue (green) and pathological tumour stages I–IV (pink-to-red gradient) for the top 10 prioritized genes from module C1 (*FTH1*, *FTL*, *ACTG1*, *NFKBIA*, *CD74*, *HSP90AB1*, *BTG1*, *VIM*, *SRGN*, *KLF6*) and the top 10 prioritized genes from module C2 (*B2M*, *EEF1A1*, *TMSB10*, *ACTB*, *PTMA*, *TPT1*, *S100A6*, *GAPDH*, *HSP90AA1*, *EIF1*). Solid black lines connect stage-group medians to illustrate monotonic progression trajectories. Inset statistics report nominal $P$ values from the omnibus five-group Kruskal–Wallis (KW) test and the two-sided Jonckheere–Terpstra (JT) ordered trend test from Normal through Stage IV (complete Benjamini–Hochberg-adjusted $q$ values and Spearman correlation coefficients are provided in Supplementary Table 4). Center lines, medians; box limits, upper and lower quartiles; whiskers, 1.5× interquartile range.
